# Astrocyte-Oligodendrocyte interaction regulates central nervous system regeneration

**DOI:** 10.1101/2022.10.04.510850

**Authors:** Irene Molina-Gonzalez, Rebecca K. Holloway, Zoeb Jiwaji, Owen Dando, Katie Emelianova, Amy F. Lloyd, Lindsey H. Forbes, Ayisha Mahmood, Thomas Skripuletz, Viktoria Gudi, James A. Febery, Jeffrey A. Johnson, Jill H. Fowler, Tanja Kuhlmann, Anna Williams, Siddharthan Chandran, Martin Stangel, Andrew J.M. Howden, Giles E. Hardingham, Veronique E. Miron

**Author notes:** Corresponding author: Veronique E. Miron.

## Abstract

Failed regeneration of myelin around neuronal axons following central nervous system damage contributes to nerve dysfunction and clinical decline in various neurological conditions, for which there is an unmet therapeutic demand^1,2^. Here, we show that interaction between glial cells – astrocytes and mature myelin-forming oligodendrocytes – is a critical determinant of remyelination. Astrocytes support the survival of regenerating oligodendrocytes, via downregulation of the Nrf2 pathway associated with increased astrocytic cholesterol biosynthesis pathway activation. Remyelination fails following sustained astrocytic Nrf2 activation yet is restored by either cholesterol biosynthesis/efflux stimulation, or Nrf2 inhibition using the existing therapeutic Luteolin. We identify that astrocyte-oligodendrocyte interaction regulates remyelination, and reveal a drug strategy for central nervous system regeneration centred on targeting this interaction.

## Main

Damage to myelin in the central nervous system (CNS) leads to loss of neuronal axon health and function in various neurological conditions (e.g. multiple sclerosis, spinal cord injury), which can be restored by the regeneration of myelin, termed remyelination^1^. This involves recruitment and differentiation of progenitor cells into mature oligodendrocytes, which must then survive to regenerate myelin^2^. However, remyelination often fails with chronic myelin injury in association with continuous clinical decline. The unmet need for regenerative therapeutics highlights the importance of identifying which cellular interactions are critical for remyelination, and how these can be targeted to restore remyelination when it fails. Although current therapeutic strategies in development are focused on direct targeting of oligodendrocyte differentiation^3^, glial-glial interactions are increasingly recognized as important regulators of myelin health^4–8^. Astrocytes are promising therapeutic targets as they are the most abundant glial cell type in the CNS and communicate with oligodendrocytes in development to support initial myelin formation^9,10^, yet their role in remyelination is understudied and controversial^11–14^. To address this, we asked whether characterization and manipulation of astrocytes during remyelination could reveal their regenerative function and a drug strategy to restore CNS regeneration.

### Astrocytes become reactive during remyelination

We first characterized astrocyte responses during efficient CNS remyelination *in vivo* by analysing focal lesions of myelin damage in the mouse corpus callosum, induced with the myelin toxin lysolecithin (LPC). This model allows for the investigation of astrocytes during remyelination specifically, without concomitant damage, and has well-defined phases of completed myelin damage (demyelination, 3 days post-injection; DPI), oligodendrocyte differentiation and survival (7 DPI), early remyelination (10 DPI) and late remyelination (14-21 DPI)^6,15,16^ (Fig. 1a). During remyelination, densities of reactive astrocytes (GFAP+) and the proportion of total astrocytes (SOX9+) which were GFAP+ were increased compared to sham (PBS)-injected and no-lesion controls (Fig.1b-d), confirmed by analysis of additional markers of astrocyte reactivity (Extended Data Fig.1). Although total astrocyte numbers were decreased early after myelin damage (3 DPI), these recovered over the course of remyelination following proliferation (Extended Data Fig.2). In an alternative model of demyelination induced by the oligodendrocyte toxin cuprizone in the diet, we observed similar changes in astrocyte responses during remyelination (Extended Data Fig.3). These data demonstrate that astrocytes become reactive during remyelination.

**Figure 1.**
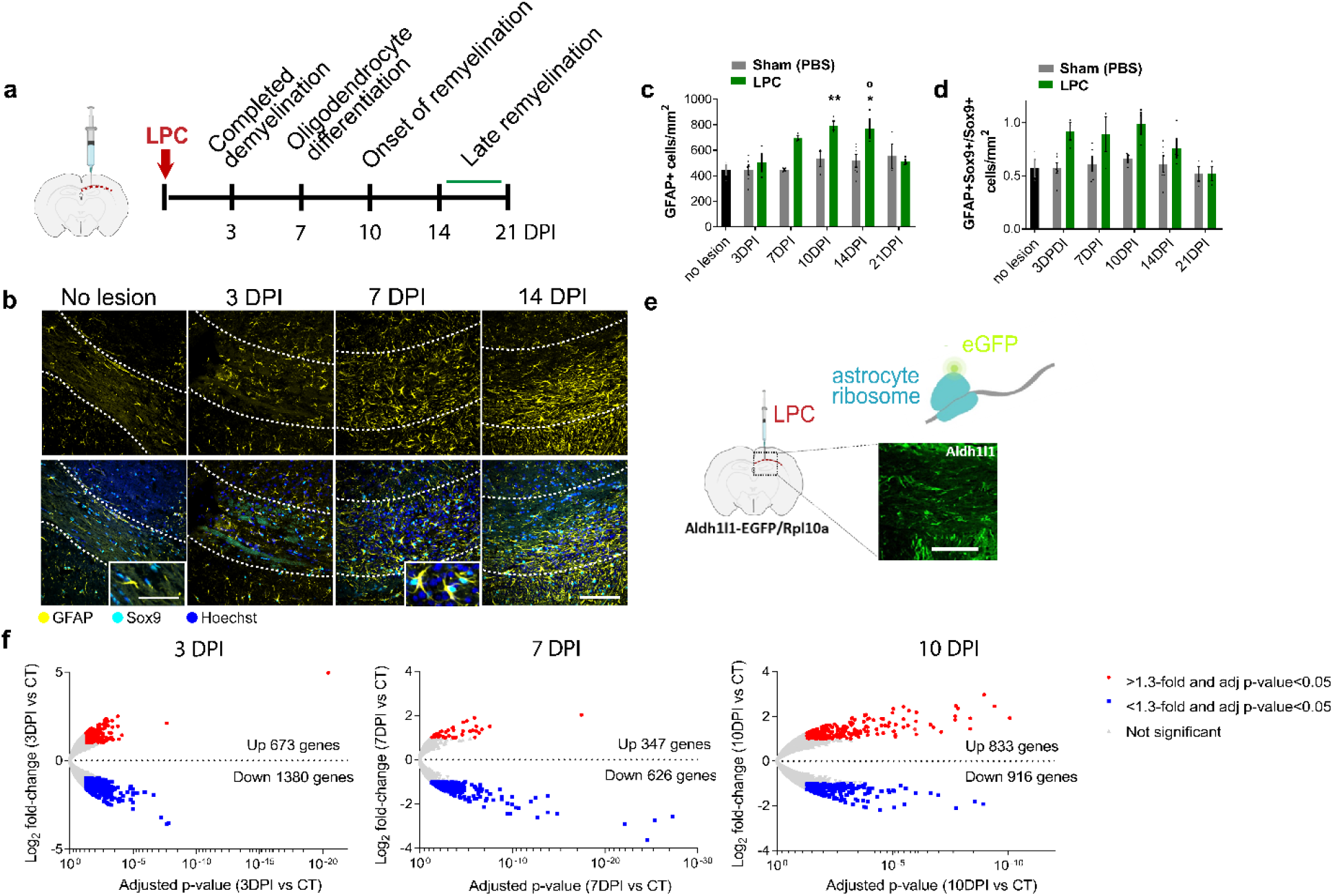
Astrocyte reactivity and translatomes dynamically change during remyelination. **a.** Astrocytes were analysed in focal LPC-demyelinated lesions of adult mouse corpus callosum at key time points representative of completed demyelination (3 DPI; days post injection), oligodendrocyte differentiation and survival (7 DPI), onset of remyelination (10 DPI) and late remyelination (14-21 DPI). **b.** Astrocytes (SOX9+; cyan) which were reactive (GFAP+; yellow) in corpus callosum (outlined) in no-lesion control and at 3, 7, and 14 DPI. Hoechst indicates nuclei in blue. Scale bar, 100 μm. Inset indicates increased astrocyte hypertrophy over time; scale bar, 50 μm. **c.** Mean GFAP+ cells/mm^2^ ± s.e.m in Sham (PBS)-injected controls and at 3, 7, 10, 14, and 21 DPI. One-way ANOVA with Tukey posthoc test versus no-lesion control (\**P*=0.0152, \*\**P*=0.0075) and sham controls (°*P*=0.0389). ANOVA summary (F=5.47 and *P*-value=0.0002). n=3-6 mice/group. **d.** Mean proportion of GFAP+SOX9+ cells/mm^2^ ± s.e.m normalized to total SOX9+ cells in no-lesion control, sham controls, and at 3, 7, 10, 14, and 21 DPI. Kruskal-Wallis test, *P-*value=0.3946. n=3-6 mice/group. **e.** TRAP of lesions in *Aldh1l1*-EGFP/Rpl10a mice allowed purification of eGFP-labelled ribosomes from astrocytes. Image of eGFP+ cells in the corpus callosum; scale bar, 50 μm. **f.** Volcano plots of adjusted P-values against Log2 fold-change following TRAP-Seq of *Aldh1l1-* EGFP/Rpl10a mice at 3 DPI, 7 DPI, and 10 DPI compared to control (CT), using a threshold of 1.3-fold and adjusted p-value of <0.05. Red indicates upregulated genes, blue represents downregulated genes, and grey indicates genes which were not significantly changed.

### Astrocytes downregulate the Nrf2 pathway and upregulate the cholesterol biosynthesis pathway during remyelination

To understand how these changes in astrocyte reactivity link to remyelination, we used translational ribosome affinity purification (TRAP) to isolate and sequence ribosome-associated mRNAs which are being actively translated in astrocytes. To achieve this, we used a transgenic mouse in which the ribosomal subunit RPl10a is tagged with eGFP driven by the pan-astrocyte-specific promoter for *Aldhl1l*^17^ (Fig.1e); we confirmed that TRAP led to enrichment of mRNAs that encode astrocyte-associated genes, and not those associated with other neural cells (Extended Data Fig.4a). Compared to no-lesion control, there were significant changes in genes throughout remyelination, with 673 upregulated and 1380 downregulated genes at 3 DPI, 347 upregulated and 626 downregulated genes at 7 DPI, and 833 upregulated and 916 downregulated genes at 10 DPI (1.3-fold change at adjusted p-value <0.05; Fig.1f; Supplemental Sheet 1). The top 25 up- or down-regulated genes were different at each time point, indicating changes in translation of genes throughout remyelination (Extended Data Fig.4). These included genes previously implicated in regulating remyelination, such as *Timp1, S1pr3, Lgals1, Lgals3, Sema3a* (3 DPI), *Hmgcs2* (7 DPI), and *Lgals3* and *Cxcl5* (10 DPI). We investigated astrocyte phenotype by assessing expression of the top 50 genes previously shown to be induced in astrocytes in models of inflammation (induced by the bacterial peptide LPS) and neuroprotection (induced by cerebral artery occlusion)^18^ and found a mixed profile at all time points (Extended Data Fig.5).

Our unbiased assessment of astrocyte phenotype identified engagement of the nuclear factor erythroid-2-related factor 2 (Nrf2) pathway at 3 DPI by Ingenuity Pathway Analysis (IPA; *‘Nrf2-mediated Oxidative Stress Response*’; Fig.2a) and Gene Ontology (GO) term analysis (‘*Wound healing involved in inflammatory responses*’; Extended Data Fig.6a). This was of particular interest since Nrf2 has previously been shown to drive anti-oxidant and neuroprotective functions in astrocytes during myelin damage^19^. Nrf2 target genes (e.g. *Hmox1*) were transiently upregulated at 3 DPI versus no-lesion control (Fig.2b-c), then subsequently downregulated at 7 DPI compared to 3 DPI (Fig.2d-e). We confirmed the transient activation of the Nrf2 pathway at 3 DPI with increased activated Nrf2+ (nuclear expression) GFAP+ cells and HMOX1+ GFAP+ cells versus controls, which subsequently decreased by 7 DPI (Fig.2f-i).

**Figure 2.**
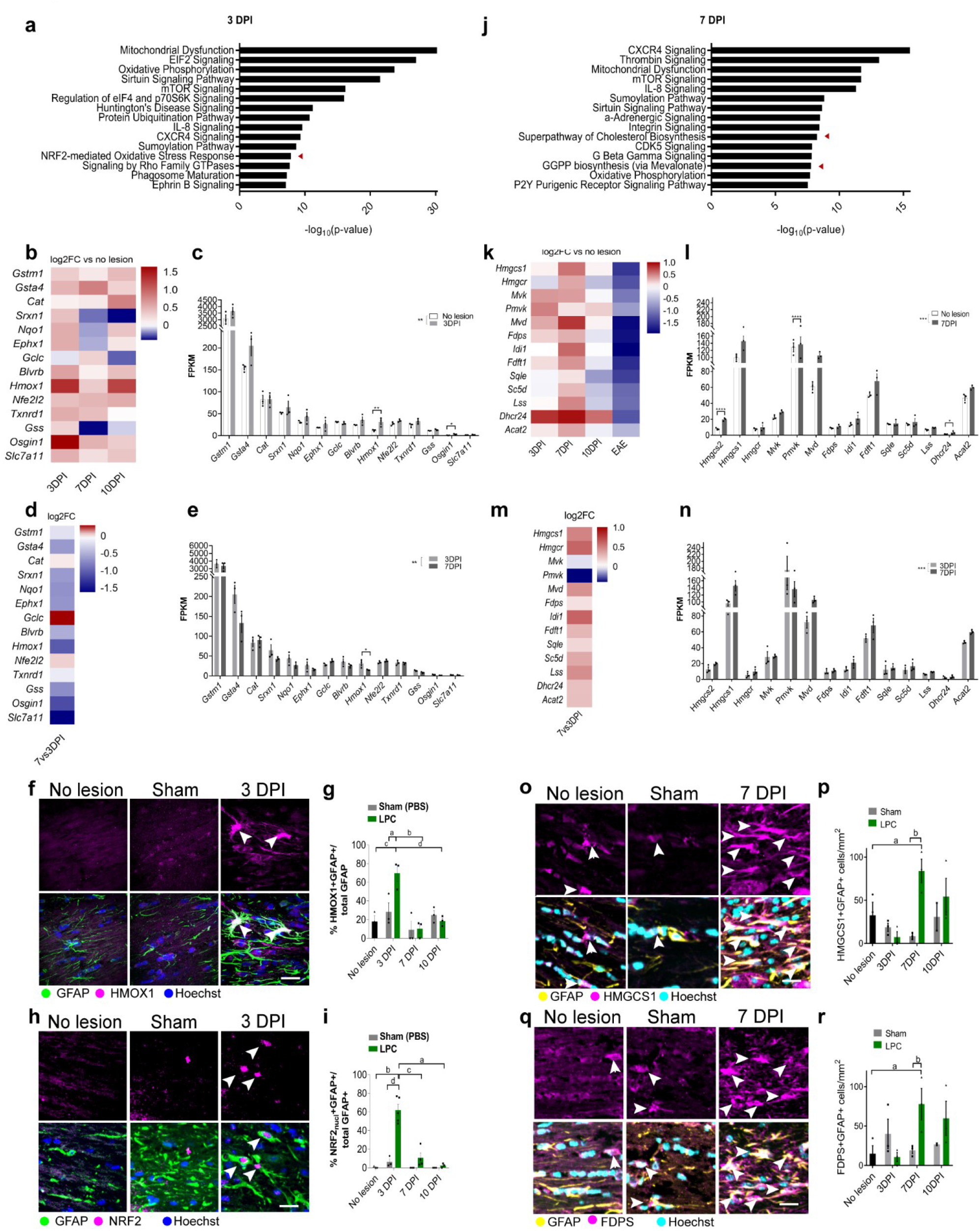
Astrocytes transiently engage the Nrf2 pathway followed by the cholesterol biosynthesis pathway during remyelination. **a.** Top 15 significantly engaged pathways in astrocytes at 3 DPI derived from Ingenuity Pathway Analysis. Arrowhead indicates the “Nrf2-mediated oxidative stress response” pathway. **b.** Heatmap of Nrf2-target genes expressed by astrocytes at 3, 7, and 10 DPI indicating Log2 fold change (FC) over no-lesion control, demonstrating upregulation at 3 DPI. **c.** Mean FPKM values ± s.e.m. of Nrf2-target genes. DE-Seq2 Benjamini–Hochberg-adjusted *P*-value, *Hmox1 P*=0.002, *Osgin1 P*=0.021. 2-tailed Paired *t* test between no lesion and 3 DPI, *P*-value=0.0054, t=3.3333. n=3 mice/ condition. **d.** Heatmap of Nrf2-target genes expressed by astrocytes at 7 DPI indicating Log2FC over 3 DPI, indicating downregulation during remyelination. **e.** Mean FPKM values ± s.e.m. of Nrf2-target genes. DE-Seq2 Benjamini–Hochberg-adjusted *P*-value, *Hmox1 P*=0.0249. 2-tailed Paired *t* test between 3 DPI and 7 DPI, *P*-value=0.0068, t=3.214 n=3 mice/ condition. **f.** GFAP+ astrocytes (green) co-stained for Nrf2-target HMOX1 (magenta) in no-lesion and sham controls and at 3 DPI. Hoechst indicates nuclei in blue. Arrows indicate double positive cells. Scale bar; 25 μm. **g.** HMOX1+GFAP+ cells/mm^2^ ± s.e.m. in no-lesion control and at 3, 7, and 10 DPI. Two-way ANOVA with Tukey’s multiple comparisons test, ^a^*P*=0.0377, ^b^*P*=0.0292, ^c^*P*=0.0280, ^d^*P=*0.0166. Two-way ANOVA summary (Interaction F(4,12)=1.838, *P*-value=0.1864; Time F(2,12)=4.66, *P*-value=0.0318; Column (condition) factor F(2,6) = 3.08, *P-*value=0.1202; Subjects (matching) F(6,12)=3.08, *P*-value=0.1324). n=3 mice/ condition. **h.** GFAP+ astrocytes (green) co-stained for Nrf2-target HMOX1 (magenta) in no lesion and sham controls and at 3 DPI. Hoechst indicates nuclei in blue. Arrows indicate double positive cells. Scale bar; 25 μm. **i**. Nrf2 (nuclear)+ GFAP+ cells/mm^2^ ± s.e.m. in no-lesion control and at 3, 7, and 10 DPI. Two-way ANOVA with Tukey’s multiple comparisons test, ^a^*P*=0.0002, ^b^*P*<0.0001, ^c^*P*=0.0004, ^d^*P*<0.0001. Two-way ANOVA summary (Interaction F(4,12)=5.897, *P*-value=0.0024; Row (Time-point) factor F(2,21) = 8.062, *P-*value=0.0025; Column (condition) factor F(2,21)=22.63, *P*-value<0.0001). n=3 mice/ condition. **j.** Top 15 significantly engaged pathways in astrocytes at 7 DPI derived from Ingenuity Pathway Analysis. Arrowheads indicate upregulation of the “Superpathway of Cholesterol biosynthesis” and “GGPP biosynthesis (via mevalonate)” pathways. **k.** Heatmap of cholesterol biosynthesis pathway genes expressed by astrocytes at 3, 7, and 10 DPI indicating Log2FC over no-lesion control, demonstrating upregulation at 7 DPI. These genes are downregulated in astrocytes in poorly-remyelinating conditions (EAE). **l.** Mean FPKM values ± s.e.m. of genes in the cholesterol synthesis pathway. DE-Seq2 Benjamini– Hochberg-adjusted *P*-value, *Hmgcs2 P*= 4×10^−7^, *Mvd P=* 5×10^−4^, *Dhcr24 P=* 0.045. 2-tailed Paired *t* test between no lesion and 7 DPI, *P*-value=0.0002, t=5.217. n=3 mice/ condition. **m.** Heatmap of cholesterol synthesis pathway genes expressed by astrocytes at 7 DPI normalized to 3 DPI indicates upregulation during the course of remyelination. **n.** Mean FPKM values ± s.e.m. of genes in the cholesterol synthesis pathway. 2-tailed Paired *t* test between no lesion and 7 DPI, *P*-value=0.0002, t=5.036. n=3 mice/ condition. **o.** GFAP+ astrocytes (yellow) co-stained with HMGCS1 (magenta) in no-lesion and sham controls and at 7 DPI. Hoechst indicates nuclei in cyan. Arrows indicate double positive cells. Scale bar; 25μm. **p.** Mean HMGCS1+ GFAP+ cells/mm^2^ ± s.e.m. in no-lesion control and at 3, 7, and 10 DPI. Two-way ANOVA with Tukey’s multiple comparisons test, ^a^*P*=0.0276, ^b^*P=*0.0015. Two-way ANOVA summary (Interaction F(4,12)=4.9575, *P*-value=0.0134; Time F(2,12)=3.75, *P*-value=0.0543; Column (condition) factor F(2,6) = 2.546, *P-*value=0.1582; Subjects (matching) F(6,12)=2.139, *P*-value=0.1236). n=3 mice/ condition. **q.** GFAP+ astrocytes (yellow) co-stained with FDPS (magenta) in no lesion and sham controls and at 7 DPI. Hoechst indicates nuclei in cyan. Arrows indicate double positive cells. Scale bar; 25μm. **r.** Mean FDPS+ GFAP· cells/mm^2^ ± s.e.m. in no-lesion control and at 3, 7, and 10 DPI. Two-way ANOVA with Tukey’s multiple comparisons test, ^a^*P*=0.0094, ^b^*P*=0.0155. Two-way ANOVA summary (Interaction F(4,12)=1.838, *P*-value=0.1864; Time F(2,12)=4.66, *P*-value=0.0318; Column (condition) factor F(2,6) = 3.08, *P-*value=0.1202; Subjects (matching) F(6,12)=2.078, *P*-value=0.1324). n=3 mice/ condition.

At 7 DPI, at the time of new oligodendrocyte generation and survival, we identified engagement of the cholesterol biosynthesis pathway by IPA (‘*Superpathway of Cholesterol Biosynthesis’, ‘GGPP biosynthesis via Mevalonate’;* Fig.2j) and GO term analysis (*‘Cholesterol metabolism’*, *‘Cholesterol homeostasis’*; Extended Data Fig.6b). Cholesterol is known to support mature oligodendrocyte survival^20^, and astrocytic cholesterol synthesis supports developmental myelination^10^ yet is dysregulated in neurological disease^21,22^; however, the role of astrocytic cholesterol in remyelination specifically is unclear. We found that genes in the cholesterol biosynthesis pathway were upregulated at 7 DPI versus no-lesion control (Fig.2k, l). Genes in the cholesterol pathway were also upregulated at 7 DPI versus 3 DPI (Fig.2m-n). In contrast, analysis of published astrocyte translatomes in the poorly remyelinating model experimental autoimmune encephalomyelitis (EAE) confirmed downregulation of these genes (Fig.2k), as previously documented^21^. Astrocytes (GFAP+) positive for enzymes involved in cholesterol synthesis, HMGCS1 and FDPS, were significantly increased at 7 DPI versus controls (Fig.2o-r), confirming engagement of this pathway.

We also observed these pathway changes in astrocytes in the cuprizone demyelination model. Densities of astrocytes with Nrf2 pathway activation were highest during the demyelination phase and early remyelination phase (3 and 5 weeks of cuprizone diet, respectively); during late-stage remyelination (7 & 10 weeks on normal diet), Nrf2-activated astrocytes were decreased while astrocytes expressing cholesterol biosynthesis enzymes were increased (Extended Data Fig.7). Our findings indicate that during remyelination, the Nrf2 pathway is de-activated in astrocytes coincident with activation of the cholesterol biosynthesis pathway. Interestingly, increased astrocytic cholesterol pathway activation occurred concomitantly with decreased activation of this pathway in the oligodendrocyte lineage at 7 DPI, suggesting potential compensation by astrocytes, whereas oligodendroglial Nrf2 activation was consistent throughout remyelination (Extended Data Fig.8).

### Manipulating astrocytic Nrf2 activation influences cholesterol pathway activation, oligodendrocyte survival and remyelination

Given that suppression of the Nrf2 pathway in the liver engages the cholesterol biosynthesis pathway^23–26^, we postulated that the downregulation of the Nrf2 pathway in astrocytes permits subsequent activation of the cholesterol biosynthesis pathway during remyelination. To test this, we prevented Nrf2 pathway downregulation by using a transgenic mouse in which Nrf2 is constitutively overexpressed in reactive astrocytes (GFAP-Nrf2), leading to persistent pathway activation^27,28^. RNA sequencing of astrocytes in this model confirmed the upregulation of the Nrf2 gene (*Nfe2l2*) and Nrf2 target genes (Fig.3a-b; Extended Data Fig.9a), and staining of tissue showed an increase in GFAP+ cells expressing the Nrf2 targets HMOX1 (Fig.3c) and NQO1 (Extended Data Fig.9b) in the corpus callosum compared to wildtype control. *Nfe2l2* upregulation was similar between astrocytes from GFAP-Nrf2 mice (fold change 2.3 versus wildtype control) and multiple sclerosis astrocytes (fold change 1.8 versus healthy control) (IPA Analysis Match^19,29^), indicating a disease-relevant modulation of astrocytic Nrf2 activation in the mice. As neural stem cells (NSCs) can also express GFAP, we verified that the NSC marker MASH1 was not expressed by >99% of nuclear Nrf2+ cells in the corpus callosum of either wildtype or GFAP-Nrf2 mice (Extended Data Fig.9c).

**Figure 3.**
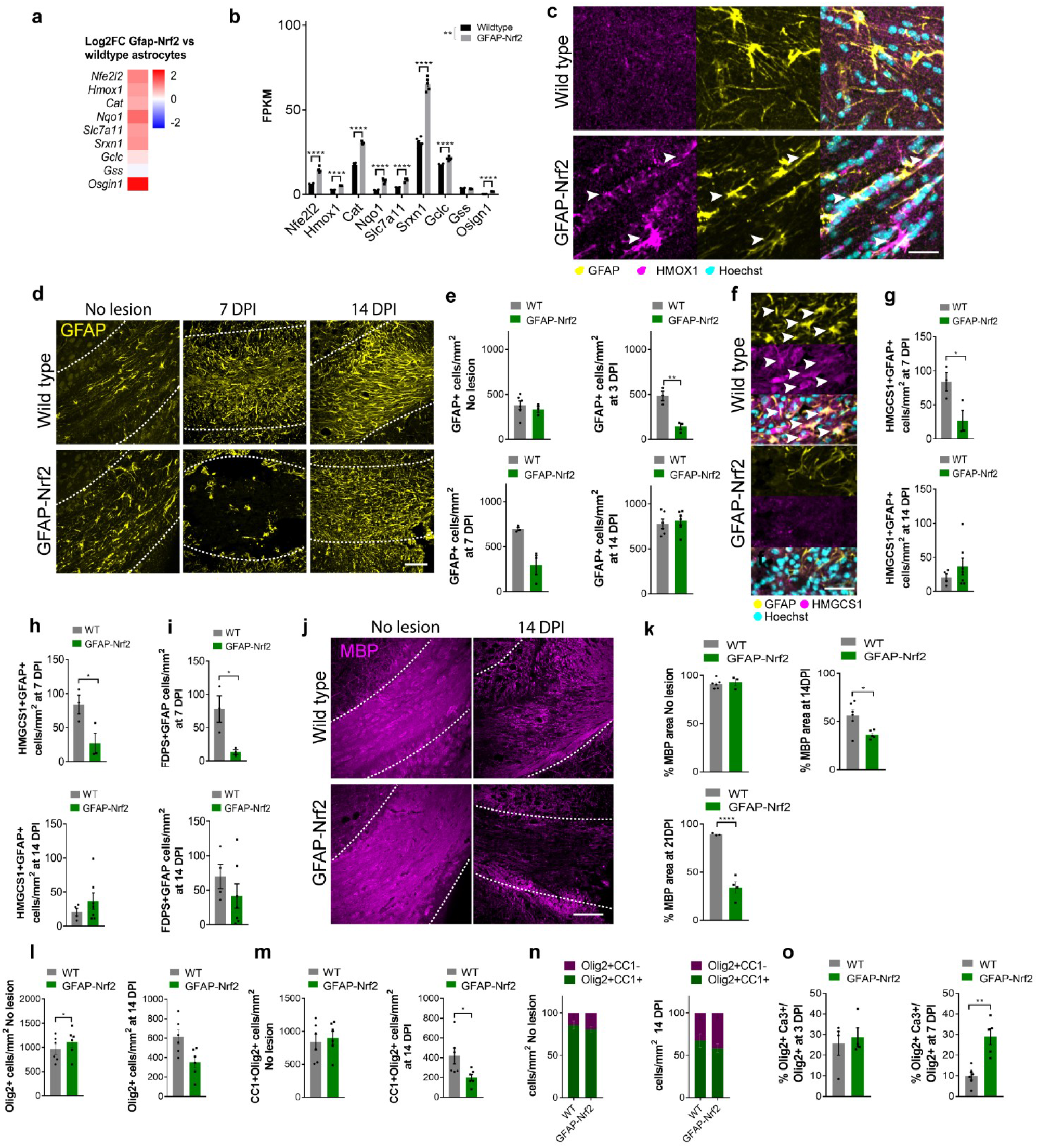
Astrocytic Nrf2 activation regulates oligodendrocyte survival and remyelination. **a.** Heatmap of Log2 fold change (FC) of Nrf2 gene (*Nfe2l2*) and Nrf2-target genes indicating upregulation in GFAP-Nrf2 astrocytes vs wildtype astrocytes. **b.** Mean FPKM values ± s.e.m. of Nrf2-target genes. DE-Seq2 Benjamini–Hochberg-adjusted *P*-value, *Nfe2l2 P*= 4.11×10^−56^, *Hmox1 P*= 2.16×10^−30^, *Cat P*= 3.88×10^−59^, *Nqo1 P*= 9.39×10^−45^, *Slc7a11 P*= 5.88×10^−37^, *Srxn1 P*= 2.27×10^−64^, *Gclc P*= 1.55×10^−9^, *Osgin1 P*= 9.43×10^−100^. 2-tailed Paired *t*-test between wild type and GFAP-Nrf2 mice, *P*-value=0.0064, t=3.66. n=5 mice/ condition. **c.** Expression of Nrf2 target HMOX1 (magenta) in astrocytes (GFAP+) in the non-lesioned corpus callosum of wild type control and GFAP-Nrf2 mice. Hoechst indicates nuclei in cyan. Arrows indicate double positive cells. Scale bar; 25μm. **d.** GFAP+ astrocytes (yellow) in wildtype and GFAP-Nrf2 mouse corpus callosum (outlined) in no-lesion control and at 7 and 14 DPI. Scale bar; 100 μm. **e.** Mean GFAP+ cells/mm^2^ ± s.e.m. in wild type (WT) and GFAP-Nrf2 mice in no lesion control and at 3, 7, and 14 DPI. Kolmogorov-Smirnov test, WT vs GFAP-Nrf2 in no lesion *P*-value: 0.3333; 2-tailed unpaired Student’s *t*-test with Welch’s correction, WT vs GFAP-Nrf2, 3 DPI *P*=0.0089 t=53.371, 7 DPI *P*=0.0591 t=3.711, 10 DPI *P*=0.5753 t=0.5791. n=3-6 mice/ condition. **f.** GFAP+ astrocytes (yellow) co-stained with HMGCS1 (magenta) in lesioned corpus callosum of wild type and GFAP-Nrf2 mice. Hoechst indicates nuclei in cyan. Arrows indicate double positive cells. Scale bar; 50 μm. **g.** Mean HMGCS1+GFAP+ cells/mm^2^ ± s.e.m. in WT and GFAP-Nrf2 mice at 7 and 14 DPI. 2-tailed unpaired Student’s *t*-test with Welch’s correction WT vs GFAP-Nrf2, 7 DPI *P=*0.0482 t=2.823, 14 DPI *P*=0.3112 t=1.1. n= 3 mice/condition. **h**. Mean percentage of GFAP+ cells positive for HMGCS1 in WT and GFAP-Nrf2 mice at 7 and 14 DPI. 2-tailed unpaired Student’s *t*-test with Welch’s correction WT vs GFAP-Nrf2, 7 DPI *P*<0.0001 t=10.11, 14 DPI *P*=0.1952 t=1.469. n=3 mice/condition. **i.** Mean FDPS+GFAP+ cells/mm^2^ ± s.e.m. in WT and GFAP-Nrf2 mice at 7 and 14 DPI. 2-tailed unpaired Student’s *t*-test with Welch’s correction WT vs GFAP-Nrf2, 7 DPI *P=*0.0398 t=3.179, 14 DPI *P=* 0.2842 t=1.148. n= 3 mice/condition. **j.** MBP staining (magenta) in the corpus callosum (outlined) in wild type and GFAP-Nrf2 mice in no lesion control and at 14 DPI. Scale bar; 100 μm. **k.** Percentage of area of corpus callosum with MBP staining ± s.e.m. in WT and GFAP-Nrf2 mice in no lesion control and at 14 and 21 DPI. 2-tailed unpaired Student’s t-test with Welch’s correction WT vs GFAP-Nrf2, no lesion *P*=0.6985 t=0.4244, 14 DPI *P*=0.0225 t=3.047, 21 DPI *P=*0.0024 t= 9.212. n=3-6 mice/condition. **l.** Mean Olig2+ cells/mm^2^ ± s.e.m. in WT and GFAP-Nrf2 mice in no lesion control and at 14 DPI. 2-tailed unpaired Student’s *t*-test with Welch’s correction WT vs GFAP-Nrf2, no lesion *P*=0.3478 t=1.174, 14 DPI *P*=0.0204 t=2.777. n=5-6 mice/condition. **m.** Mean CC1+Olig2+ cells/mm^2^ ± s.e.m in WT and GFAP-Nrf2 mice in no-lesion control and at 14 DPI. 2-tailed unpaired Student’s *t*-test with Welch’s correction WT vs GFAP-Nrf2, no lesion *P*=0.4435 t=0.865, 14 DPI *P*=0.0425 t=2.508. n=5-6 mice/condition. **n.** Proportion of Olig2+ cells that are CC1+ (green) or negative (magenta) ± s.e.m in WT and GFAP-Nrf2 mice in no lesion control and at 14 DPI. n=5-6 mice/condition. 2-way ANOVA with Bonferroni correction, WT vs GFAP-Nrf2; no lesion, CC1+Olig2+ *P*=0.8323 CC1-Olig2+ *P*=0.8322, two-way ANOVA summary (Interaction F(1,20)=1.379, *P*-value=0.2541; Row (mouse genotype) Factor F(1,20)=3.171×10^−9^, *P*-value>0.9999; Column (cell type) factor F(1,20) = 242.9, *P*-value<0.0001). 14 DPI, CC1+Olig2+ *P*=0.6945 CC1-Olig2+ *P*=0.6945, two-way ANOVA summary (Interaction F(1,20)=1.853, *P*-value=0.1885; Row Factor F(1,20)=0, *P*-value>0.9999; Column factor F(1,20) = 14.81, *P*-value=0.0010). **o.** Mean percentage of Olig2+ cells which are cleaved Caspase-3+ ± s.e.m in WT and GFAP-Nrf2 mice at 3 DPI, Kolmogorov-Smirnov test, *P*=0.7714; at 7 DPI, 2-tailed unpaired Student’s *t*-test with Welch’s correction *P*=0.0044 t=4.49. n=4-6 mice/condition.

Early after demyelination, lesioned GFAP-Nrf2 mice showed decreased densities of reactive astrocytes (GFAP+; Fig.3d-e) and total astrocytes (SOX9+; Extended Data Fig.9f-g), which recovered by 14 DPI. We confirmed that reduced SOX9+ cells did not represent changes in oligodendrocytes, as the oligodendrocyte lineage marker SOX10 was not expressed by the majority of SOX9+ cells nor by >90% of GFAP+SOX9+ cells (Extended Data Fig.9h-j). The reduction in astrocytes in GFAP-Nrf2 lesions may have been partially due to increased astrocyte apoptosis (active-Caspase-3+ GFAP+; Extended Data Fig.9k-l). We then assessed the impact of astrocytic Nrf2 activation on the cholesterol biosynthesis pathway, and found that the number of lesion GFAP+ cells that expressed cholesterol biosynthesis enzymes HMGCS1, FDPS, MVD and FDFT1 was significantly decreased in GFAP-Nrf2 mice compared to wildtype controls at 7 and 14 DPI (Fig.3f-g, Extended Data Fig.9 c-d). The percentage of lesion astrocytes expressing HMGCS1 at 7 DPI was reduced from 40.2 ± 4.6 % of GFAP+ cells in wildtype control to 7.5 ± 5.0 % in GFAP-Nrf2 mice, which remained low by 14 DPI (Fig.3g), indicating reduced engagement of the cholesterol biosynthesis pathway in surviving astrocytes in GFAP-Nrf2 mice. This was associated with an impairment in remyelination in GFAP-Nrf2 lesions at the time when it is normally robust (14 and 21 DPI), indicated by decreased expression of myelin proteins MBP (Fig.3j-k) and CNPase (Extended Data Fig.10a-b). Impaired remyelination in GFAP-Nrf2 mice was not the result of disrupted myelination at baseline as non-lesioned corpus callosum was unaffected (Fig. 3j-k), nor was demyelination more severe as the initial LPC-induced lesion was comparable to wildtype control (Extended Data Fig.10c-d).

We did however observe decreased densities of total oligodendrocyte lineage cells (Olig2+; Fig.3l) and oligodendrocytes (CC1+Olig2+; Fig.3m) at 14 DPI in GFAP-Nrf2 mice, whereas these were unaffected in non-lesioned mice (Fig.3l-m). The proportion of Olig2+ cells which were CC1+ or CC1-was not significantly affected in lesioned GFAP-Nrf2 mice (Fig.3n), suggesting that oligodendrocyte differentiation was not impacted. To determine the cause of reduced oligodendrocyte lineage cells, we investigated their responses at earlier time points. No change was observed in the proportion of oligodendrocyte progenitor cells (OPCs; nuclear Olig1+) that were proliferating at 3 or 7 DPI (Ki67+; Extended Data Fig.10e-f). However, GFAP-Nrf2 mice showed a significant increase in the proportion of oligodendrocyte lineage cells undergoing cell death at 7 DPI (Olig2+ active-Caspase-3+; Fig.3o, Extended Data Fig.10g), associated with a reduction in total Olig2+ cells (Extended Data Fig.10h). These were found to be mature oligodendrocytes, as we observed a significant increase in the proportion of CC1+ cells which were active-Caspase-3+ (Extended Data Fig.10i-j). These dysregulated oligodendrocyte responses in GFAP-Nrf2 lesions did not result from altered microglial numbers nor their phagocytosis of myelin debris, which were not significantly changed compared to control (Extended Data Fig.10k-l). These findings associate sustained Nrf2 activation in astrocytes with reduced oligodendrocyte survival and poor remyelination.

As GFAP-Nrf2 lesions showed both a reduction in astrocyte numbers and impaired astrocytic cholesterol biosynthesis pathway activation, we next tested the contribution of both processes to oligodendrocyte death during remyelination. First, we asked whether inducing astrocyte death is sufficient to reduce oligodendrocyte survival. We induced astrocyte death during remyelination using transgenic mice in which GFAP-driven expression of thymidine-kinase renders astrocytes selectively vulnerable to apoptosis following ganciclovir administration, using the cuprizone model where this paradigm has been previously optimized^12^. Astrocyte death was induced in early-phase remyelination (weeks 4-5 of cuprizone administration; Extended Data Fig.11 a-c), when the cholesterol pathway is not yet upregulated (Extended Data Fig.7e-h). This was associated with a decrease in oligodendrocytes (CC1+) and increase in apoptotic oligodendrocytes (active-Caspase-3+ CC1+) (Extended Data Fig.11b,e-f).

Second, we asked whether poor oligodendrocyte survival in GFAP-Nrf2 mice could be rescued by reinstating cholesterol biosynthesis pathway activation in astrocytes. We achieved this using a blood-brain-barrier (BBB)-permeable and molecularly specific agonist of the cholesterol transporter ABCA1 (CS-6253)^30^, previously used to increase expression of cholesterol biosynthesis genes in astrocytes^21^ as they are the primary expressors of ABCA1 (www.brainrnaseq.org). We assessed ABCA1 expression during remyelination, first by proteomic analysis of wildtype lesions which indicated increased ABCA1 levels at 7 DPI alongside several cholesterol biosynthesis pathway proteins (Fig.4a); our dataset was biased towards detecting astrocyte-associated proteins (Fig.4b) whereas microglial proteins CD68, TMEM119, and P2RY12 were not detected likely due to sensitivity thresholds. We confirmed ABCA1 gene and protein expression were upregulated in astrocytes over the course of remyelination (Fig.4 c-e), with >80% of ABCA1+ cells being GFAP+ between 7 and 14 DPI (Fig.4f). CS-6253 or vehicle control was administered to lesioned GFAP-Nrf2 mice at the time when the cholesterol biosynthesis pathway is activated in wildtype mice (7 DPI) until remyelination would normally be underway (14 DPI) (Fig.4g). CS-6253 significantly increased the percentage of cholesterol biosynthesis enzymes in astrocytes (HMGCS1+GFAP+) compared to vehicle control (Fig.4h, i). CS-6253 did not significantly impact cholesterol biosynthesis enzyme expression by oligodendrocytes or microglia (Extended Data Fig.12a-d), nor microglial lesion coverage (Extended Data Fig.12e-f). Nonetheless, the possibility of off-target effects on other cell types must be taken into consideration.

**Figure 4.**
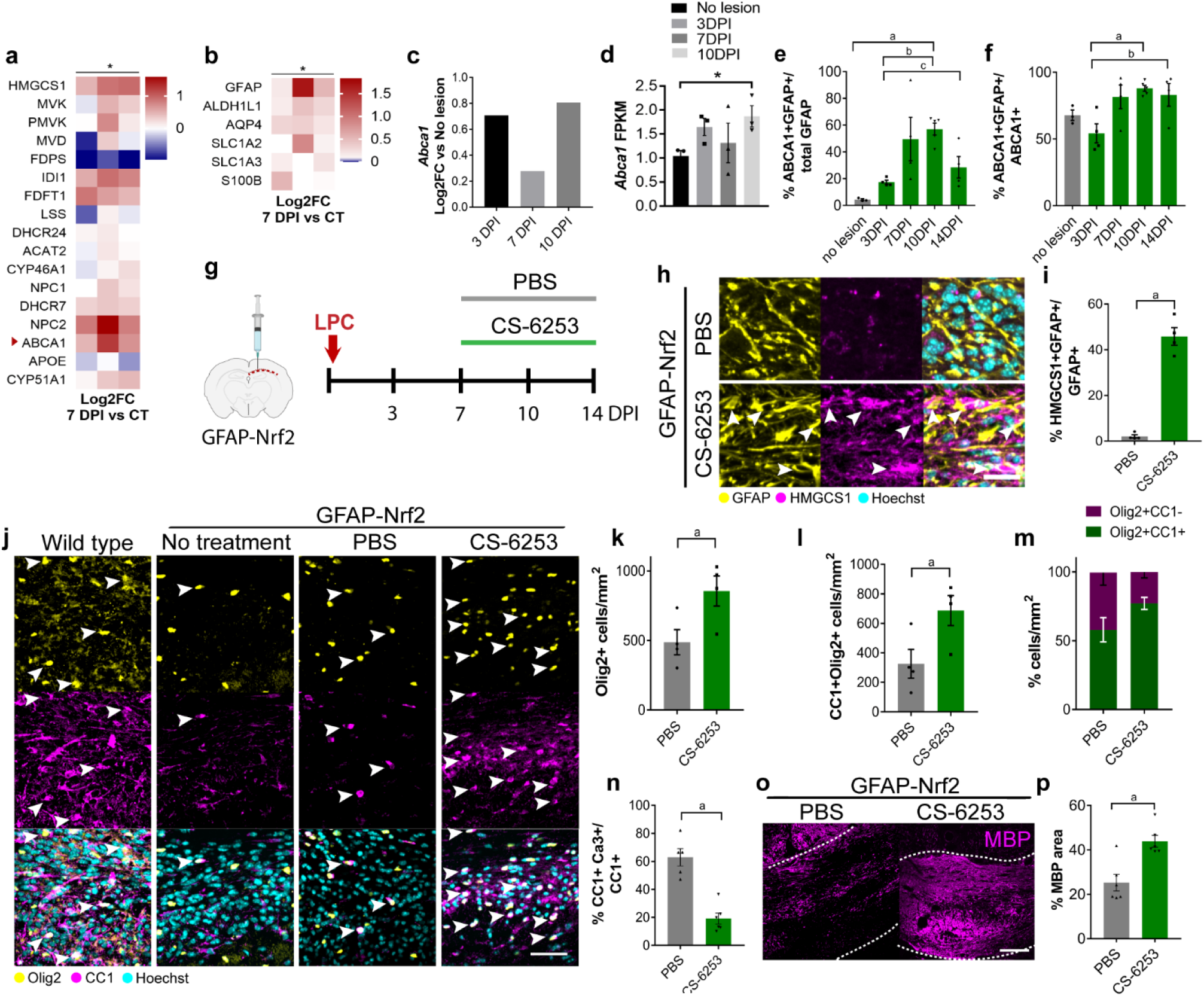
Stimulating the cholesterol biosynthesis pathway rescues oligodendrocyte survival and remyelination when astrocytic Nrf2 activation is sustained. **a.** Proteomic analysis of wild type lesions at 7 DPI vs no lesion control (CT), represented as Log2FC DPI/CT. 2-tailed Paired *t-*test between CT and 7 DPI on normalized intensities, *P*-value=0.0449, t=2.176. n=3 mice/condition. **b.** Upregulation of astrocyte-associated proteins in lesions at 7 DPI vs CT, represented as Log2FC. 2-tailed Paired *t-*test between CT and 7 DPI on normalized intensities, *P*-value=0.0217, t=3.291. n=3 mice/condition. **c.** *Abca1* Log2FC vs no lesion control at 3, 7, and 10 DPI. n=3 mice/condition. **d**. Mean FPKM value ± s.e.m. of *Abca1*. DE-Seq2 Benjamini–Hochberg-adjusted *P*-value ^a^*P*=0.042. n=3 mice/ condition. **e.** Percentage of GFAP+ cells expressing ABCA1 ± s.e.m. in no-lesion controls and at 3-14 DPI. One way ANOVA with Tukey’s multiple comparisons test; ^a^*P*=0.0237, ^b^*P*=0.0054, ^c^*P*=0.0245. ANOVA summary (F=6.456 and *P-*value=0.0031). n=3-5 mice/condition. **f.** Percentage of ABCA1+ cells expressing GFAP ± s.e.m. in no-lesion controls and at 3-14 DPI. One way ANOVA with Tukey’s multiple comparisons test; ^a^*P*=0.0126, ^b^*P*=0.0496. ANOVA summary (F=9.985 and *P-*value=0.0004). n=6 mice/condition. **g.** Focal LPC-demyelination was performed in adult GFAP-Nrf2 corpus callosum and CS-6253 or PBS was administrated daily from 7-14 DPI. **h.** Astrocytes (GFAP+; yellow) co-stained with HMGCS1 (magenta) in PBS or CS-6253 treated GFAP-Nrf2 mice at 14 DPI. Hoechst indicates nuclei in cyan. Arrows indicate double positive cells. Scale bar; 25μm. **i.** Percentage of GFAP+ cells expressing HMGCS1 ± s.e.m in PBS and CS-6253 treated GFAP-Nrf2 mice at 14 DPI, Kolmogorov-Smirnov test, ^a^*P=*0.0286. n=4 mice/condition. **j.** Oligodendrocyte lineage cells (Olig2+; yellow) which are mature (CC1+; magenta) at 14 DPI in wild type and GFAP-Nrf2 mice which were non-treated, or treated with PBS or CS-6253. Hoechst indicates nuclei in cyan. Arrows indicate double positive cells. Scale bar; 50μm. **k.** Mean Olig2+ cells/mm^2^ ± s.e.m. in PBS and CS-6253-treated GFAP-Nrf2 mice at 14 DPI, 2-tailed unpaired Student’s *t*-test with Welch’s correction, ^a^*P=*0.0413, t=2.61. n=3-4 mice/condition. **l.** Mean CC1+Olig2+ cells/mm^2^ ± s.e.m. in PBS and CS-6253-treated GFAP-Nrf2 mice at 14 DPI, 2-tailed unpaired Student’s *t*-test with Welch’s correction, ^a^*P=*0.0432, t=2.555. n=3-4 mice/condition. **m.** Proportion of Olig2+ cells which were CC1+ (green) or CC1-(magenta) ± s.e.m. in PBS and CS-6253-treated GFAP-Nrf2 mice at 14 DPI. Two-way ANOVA with Bonferroni correction PBS vs CS6253, CC1+Olig2+ *P*-value=0.1597 CC1-Olig2+ *P*-value=0.1734. Two-way ANOVA summary (Interaction F(1,12)= 7.141, *P*-value=0.0203; Row (condition) Factor F(1,12)=0.001127, *P*-value=0.9738; Column (type of cell) factor F(1,12) = 24.77, *P*-value=0.0003). n=3-4 mice/group **n**. Percentage of CC1+ cells expressing active caspase-3 ± s.e.m. in PBS and CS-6253-treated GFAP-Nrf2 mice at 14 DPI, 2-tailed unpaired Student’s *t*-test with Welch’s correction, ^a^*P*=0.0005, t=6.007. n=5-6 mice/condition. **o.** Representative images of myelin basic protein (MBP; magenta) in corpus callosum lesions (outlined) of GFAP-Nrf2 mice treated with PBS or CS-6253. Scale bar, 100 μm. **p**. Percentage of corpus callosum area covered by MBP ± s.e.m. in PBS and CS-6253-treated GFAP-Nrf2 mice at 14 DPI, Kolmogorov-Smirnov test, ^a^*P*=0.0260, n=6 mice/condition.

CS-6253 administration increased numbers of oligodendrocyte lineage cells (Olig2+; Fig.4j-k) and oligodendrocytes (CC1+Olig2+; Fig.4j, l), increased the proportion of Olig2+ cells which were CC1+ (Fig.4m), and significantly decreased apoptotic oligodendrocytes (active caspase 3+ CC1+) in GFAP-Nrf2 mice compared to vehicle control (Fig. 4n). This led to a rescue in remyelination at 14 DPI (Fig.4o-p). Altogether, these findings demonstrate that sustained Nrf2 activity in astrocytes compromises oligodendrocyte survival and remyelination, and suggest that this could be partly due to an impairment in the astrocytic cholesterol biosynthesis pathway.

Having observed that sustained astrocytic Nrf2 activation inhibits cholesterol pathway activation and remyelination, we next asked whether loss-of-function knockout of Nrf2 in astrocytes would have opposing effects. Due to the importance of astrocytic Nrf2 for neuronal health, we avoided potentially deleterious impacts of CNS-wide knockout by generating a localized knockout of *Nfe2l2* in astrocytes in the corpus callosum. *Nfe2l2* floxed mice were stereotaxically injected with an AAV5 inducing GFAP promoter-driven Cre recombinase and GFP expression (‘AAV-Cre), as done previously^31–33^, allowed to recombine over 5 days, before injecting LPC to induce a lesion. At 3 DPI in the AAV-Cre lesions, >89% of GFP+ signal was GFAP+ and >98% of GFAP+ cells were GFP+ (Extended Data Fig. 13a-e); nuclear Nrf2+GFAP+ cells were significantly decreased to 16%, compared to 63% in mice injected with a sham virus (AAV5-GFAP-eGFP; ‘AAV Sham’) (Extended Data Fig.13f-h). As the LPC model is normally associated with reduced astrocytic Nrf2 at 7 DPI, inducing this downregulation ahead of schedule by gene deletion resulted in an earlier upregulation of expression of cholesterol biosynthesis enzymes HMGCS1, FDPS, MVD, and FDFT1 in astrocytes at 3DPI, which was maintained by 7 DPI (Extended Data Fig.13i-m), confirming Nrf2 impact on this pathway. There were no significant impacts of Nrf2 conditional knockout on astrocyte number or demyelination (Extended Data Fig.13n-p). This genetically-induced early downregulation of Nrf2 caused a small but significant increase in the level of remyelination at 7 DPI, a time point prior to the robust remyelination at later time points in controls (Extended Data Fig.13q-r). The small impact on remyelination likely reflects the low number of newly generated/ apoptotic oligodendrocytes which could be influenced by astrocytic responses this early in remyelination. Altogether, these findings indicate that Nrf2 regulation in astrocytes is sufficient to influence the cholesterol pathway and remyelination.

### Astrocytes export cholesterol to oligodendrocytes in vitro to regulate their survival and remyelination

In development, newly generated oligodendrocytes are susceptible to cell death, yet cholesterol supports their survival^10,20^. We therefore asked whether astrocytes support oligodendrocyte survival by directly exporting cholesterol to oligodendrocytes. As to our knowledge, technologies to assess cholesterol transfer between specific cell types *in vivo* are non-existent, precluding *in vivo* assessment. Therefore, as a proof-of-concept, we adopted a simplified *in vitro* model using primary cultures to directly manipulate isolated cell types. To assess the role of astrocytic Nrf2 activation in this process, we treated cultured primary astrocytes with the potent Nrf2 stimulator, the triterpenoid CDDO^TFEA 34,35^(Fig.5a), and confirmed increased activated nuclear Nrf2 (Fig. 5b, c) which was abrogated using Luteolin, a flavonoid previously used reduce Nrf2 activation when hyperactivated^36^ (Fig.5b, c). CDDO^TFEA^ upregulated Nrf2 pathway gene expression, as expected, and downregulated expression of cholesterol pathway genes (Fig.5d), demonstrating suppression of the cholesterol pathway by Nrf2 activation. Select cholesterol biosynthesis pathway gene expression was impacted in CDDO^TFEA^-activated astrocytes with Luteolin, or cholesterol biosynthesis pathway/efflux stimulation with CS-6253 (Fig.5a,e-f).

**Figure 5.**
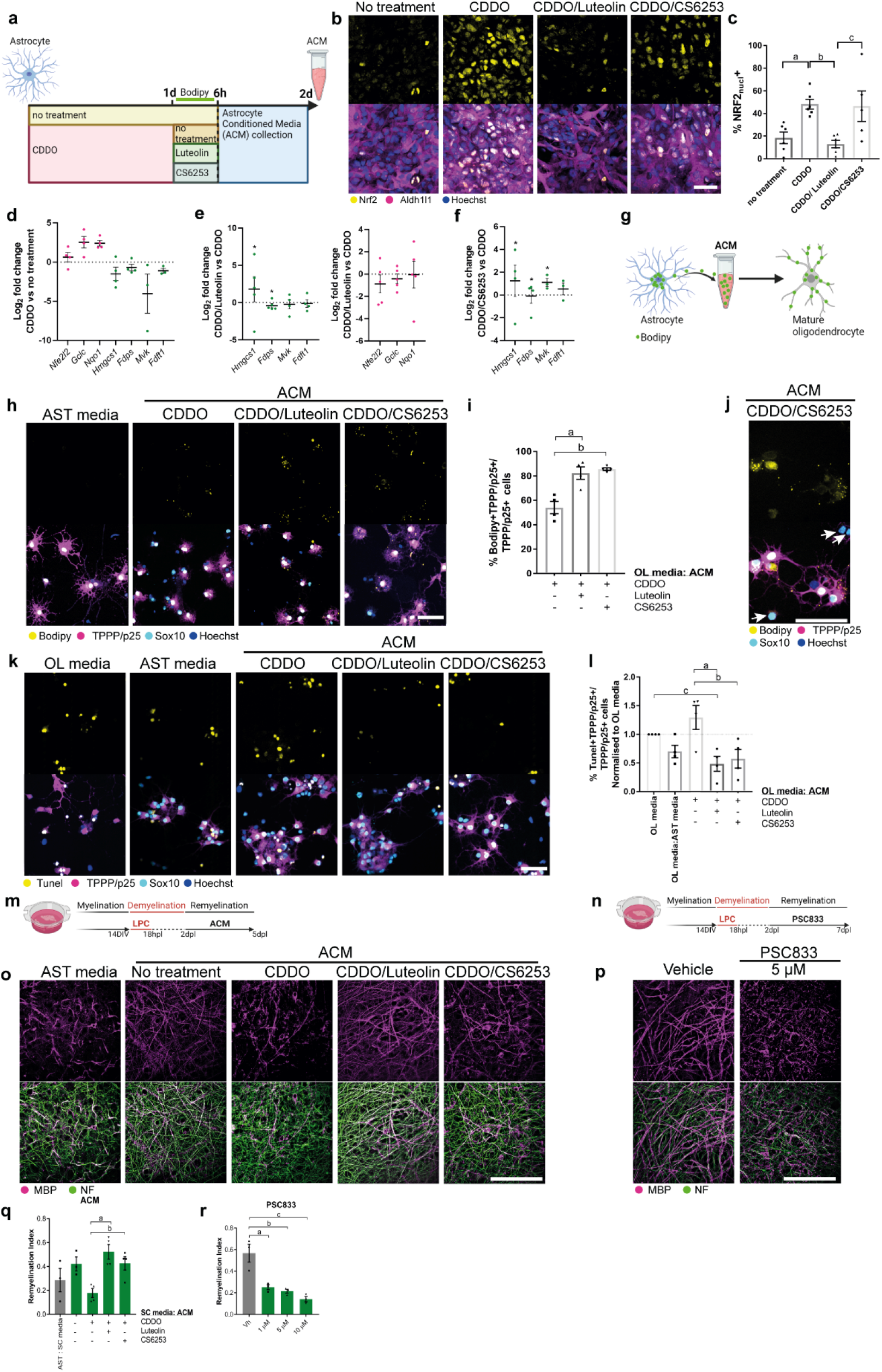
Astrocytes export cholesterol to oligodendrocytes to regulate their survival and remyelination. **a.** Primary astrocytes were either untreated or treated with the Nrf2 activator CDDO^TFEA^ for 1d, then either untreated or treated with Luteolin or CS-6253, during which time they were loaded with fluorescent cholesterol analogue Bodipy-FL-C12. Excess analogue was washed off, then exported into astrocyte conditioned media (ACM) over 18 hours. **b.** Astrocytes (Aldh1l1; magenta) expressing Nrf2 (yellow) and counterstained with Hoechst (blue) following treatment with CDDO^TFEA^, or CDDO^TFEA^ then Luteolin, or CDDO^TFEA^ then CS-6253. Scale bar, 50 μm. **c.** Mean percentage of astrocytes expressing nuclear Nrf2 ± s.e.m. One way ANOVA and Tukey’s multiple comparison test; ^a^*P*=0.0286, ^b^*P*=0.0085, ^c^*P*=0.0178. n=4 independent litters. **d.** Astrocyte expression of genes involved in cholesterol and Nrf2 signalling following CDDO^TFEA^ treatment, represented as Log2FC over no treatment condition. Comparison of all genes as a set using 2-tailed Wilcoxon test of all Nrf2 and cholesterol pathway genes between CDDO and no treatment, *P*-value=0.0313. n=3-4 independent litters. **e.** Astrocyte expression of genes after CDDO^TFEA^ treatment followed by Luteolin treatment, represented as Log2FC over CDDO^TFEA^ condition. Comparison of specific genes between CDDO vs CDDO/Luteolin by Kolmogorov-Smirnov tests, *Hmgcs1 *P*=0.0476, *Fdps *P*=0.0286. n=4-5 independent litters. **f.** Astrocyte expression of genes after CDDO^TFEA^ treatment followed by CS-6253 treatment, represented as Log2FC over CDDO^TFEA^ condition. Comparison of specific genes using CDDO vs CDDO/CS6253 by Kolmogorov-Smirnov tests, *Hmgcs1 *P*=0.0476, *Fdps *P*=0.0476, *Mvk *P*=0.0286. n=4-5 independent litters. **g.** ACM was applied to primary cultures of mature oligodendrocytes to track uptake of fluorescent cholesterol analogue, Bodipy-FL-C12, exported from astrocytes. **h.** Representative images of mature oligodendrocytes (TPPP/p25+ Sox10+; magenta/cyan) in unconditioned astrocyte media (AST) or following exposure to ACM, and uptake of Bodipy-FL-C12 (yellow). Hoechst indicates nuclei in blue. Scale bar, 50 μm. **i.** Percentage of TPPP/p25+ cells having taken up cholesterol analogue from ACM (Bodipy+) ± s.e.m.. Kruskal-Wallis test, ^a^*P*=0.0561, ^b^*P*=0.0360. n=4 independent litters. **j.** Immature oligodendrocyte lineage cells (Sox10+ TPPP/p25-) were Bodipy negative (arrows). Scale bar, 50 μm. **k.** Representative images of apoptotic (TUNEL+; yellow) TPPP/p25+ Sox10+ oligodendrocytes (magenta/cyan), in oligodendrocyte media (OL) or unconditioned astrocyte media (AST) controls, or following exposure to ACM. Scale bar, 50 μm. **l.** Mean percentage of TPPP/p25+ cells which are TUNEL+, normalized to OL media control. Kruskal-Wallis test *P-*value=0.0349, ^a^*P*=0.0083, ^b^*P=*0.0265, ^c^*P*=0.0265. n=4 independent litters. **m.** Brain explants were allowed to myelinate for 14 days *in vitro* (DIV), demyelinated with LPC, then fixed at 5 days post-LPC (dpl) when remyelination is initiated. **n.** Representative images of brain explants exposed to unconditioned astrocyte media (AST) or ACM from astrocytes following no treatment or exposure to CDDO^TFEA^, CDDO^TFEA^ then Luteolin, and CDDO^TFEA^ then CS-6253 stained for myelin basic protein (MBP; magenta) and neurofilament (NF; green). Scale bar, 50 μm. **o.** Remyelination index for AST: slice culture media (SC) control, or following exposure to ACM. One-way ANOVA with Tukey’s multiple comparisons test, ^a^*P*=0.0078, ^b^*P*=0.0634. ANOVA summary (F=5.208 and *P-*value=0.0100). n=3-4 mice/condition. **p.** Brain explants were allowed to myelinate for 14 DIV, demyelinated with LPC, then treated with the ABCA1 inhibitor PSC833 (1, 5, 10 μM) or vehicle control from 2 to 7 dpl when remyelination is underway. **q.** Representative images of brain explants treated with vehicle control or PSC833 (5 μM) and stained for myelin basic protein (MBP; magenta) and neurofilament (NF; green). Scale bar, 50 μm. **r.** Remyelination index for vehicle or PSC833-treated brain explants. One-way ANOVA with Tukey’s multiple comparisons test, ^a^*P*=0.0052, ^b^*P*=0.0026, ^c^*P*=0.0007. ANOVA summary (F=17.11 and *P-*value=0.0008). n=3 mice/condition.

Using a previously established *in vitro* method to track direct cholesterol transfer between cell types^37^, we loaded astrocytes with the fluorescent cholesterol analogue Bodipy-FL-C12 for 6 hours (Fig.5a; Extended Data Fig.14), washing off excess then allowing 18 hours for analogue efflux into astrocyte conditioned media (ACM), which was then applied to mature oligodendrocytes (Fig.5g). Astrocyte cultures were confirmed to be pure via expression of markers Aldh1l1 and GFAP (Extended Data Fig.14). Cholesterol analogue was taken up from ACM by mature oligodendrocytes (Bodipy+ TPPP/p25+; Fig.5h-j). An increased proportion of oligodendrocytes were Bodipy+ following treatment with ACM where astrocytic Nrf2 was inhibited (CDDO^TFEA^/Luteolin ACM) or the Nrf2 impact on the cholesterol synthesis pathway was corrected (CDDO^TFEA^/CS-6253 ACM), compared to ACM generated from CDDO^TFEA^-treatment alone (Fig.5h-i). Immature oligodendrocyte lineage cells (Sox10+ TPPP/p25-) were not Bodipy+ (Fig. 5j). We next assessed the effect of ACM on oligodendrocyte survival. Oligodendrocytes matured in culture show a high level of baseline apoptosis (TUNEL+ TPPP/p25+) (‘OL media’; Fig.5k). CDDO^TFEA^ ACM increased the percentage of apoptotic oligodendrocytes even further, which was not observed with CDDO^TFEA^/Luteolin or CDDO^TFEA^/CS-6253 ACM (Fig.5k,l). These results demonstrate a correlation between cholesterol analogue uptake from ACM and oligodendrocyte survival *in vitro*. To assess relevance for remyelination, we applied ACM to brain explants after LPC-induced demyelination is complete (2 days post-LPC; dpl) until initiation of remyelination (5 dpl) (Fig.5m). Whereas control explants exposed to unconditioned astrocyte media had little remyelination at this time, this was enhanced by exposure to untreated ACM (Fig.5n) as measured by remyelination index (MBP and NF colocalization normalized to NF; Fig.5o). Conversely, exposure of brain explants to CDDO^TFEA^ ACM worsened remyelination, which was rescued by CDDO^TFEA^/Luteolin or CDDO^TFEA^/CS6253 ACM (Fig.5n,o). Although we recognize the limitations of *in vitro* experiments, this provided a proof-of-concept that in principle astrocytes have the ability to export cholesterol to oligodendrocytes, this process is regulated by Nrf2, and this influences oligodendrocyte survival and remyelination *in vitro/ex vivo*. Future work is required to address this important question *in vivo* with the advent of new technologies.

We next hypothesized that astrocyte cholesterol efflux is a critical regulator of remyelination, which we tested inhibiting cholesterol efflux from astrocytes using the ABCA1 inhibitor PSC833. Due to the inability of PSC833 to cross the blood brain barrier, and the requirement for long-term treatment post-demyelination which precluded intracerebral injection, we applied the inhibitor to brain explants following demyelination (2 dpl) until remyelination is normally underway (7 dpl; Fig.5p). Remyelination was impaired following treatment with PSC833 (1, 5, 10 μM) compared to vehicle control (Fig.5q,r). These data suggest that astrocytes could regulate oligodendrocyte survival and remyelination via cholesterol efflux, yet this is impaired by Nrf2 activation.

### Astrocytic Nrf2 activation in chronic human CNS lesions is associated with decreased cholesterol pathway activation and oligodendrocyte death

As Nrf2 activation in astrocytes is known to undergo changes in the human demyelinating disease multiple sclerosis (MS)^19,38,39^, we investigated whether this was associated with altered cholesterol biosynthesis pathway activation. We analysed published single nuclei sequencing data of distinct MS lesion types in which remyelination potential is high (‘active’) or poor (‘inactive’) compared to controls who died of non-neurological causes^29^. Astrocytes were identified using the markers AQP4 and GFAP, extracted using Seurat, and re-clustered at a resolution of 0.3 with a threshold of <15% mitochondrial RNA. Eight astrocyte clusters were identified, which demonstrated differential prevalence in MS lesions vs control, and active vs inactive lesions (Extended Data Fig.15a). In particular, clusters 1 and 2 were predominantly detected in inactive lesions (Extended Data Fig.15b). Of these two, cluster 1 astrocytes had enrichment of Nrf2-associated genes compared to all other clusters (P=0.0039, Fisher’s Exact Test and Odds ratio 2.343). For instance, classical Nrf2 target genes *SLC7A11, MGST1*, and *SQSTM1* were significantly increased in cluster 1 astrocytes compared to other clusters (Extended Data Fig. 15c). These same genes were also significantly increased in GFAP-Nrf2 astrocytes to a similar extent (Extended Data Fig.15d). We found that cluster 1 astrocytes downregulated cholesterol efflux genes (Extended Data Fig.15e), thereby showing in human disease an association between Nrf2 engagement with cholesterol pathway downregulation in inactive MS lesion astrocytes. We validated these findings at the protein level (cases in Extended Data Table 1), finding that inactive lesions had the highest proportion of astrocytes positive for markers of Nrf2 activation (Nrf2, HMOX1, NQO1; Fig.6a-b, Extended Data Fig.15f-h) compared to active lesions, lesions that had already fully remyelinated, and controls. In contrast, these inactive lesions had a reduced proportion of astrocytes positive for the cholesterol biosynthesis pathway enzyme HMGCS1 compared to control and active or remyelinated lesions (Fig.6c-d). Overall, these findings indicate activation of the Nrf2 pathway and suppression of cholesterol biosynthesis pathway in inactive MS lesion astrocytes.

**Figure 6.**
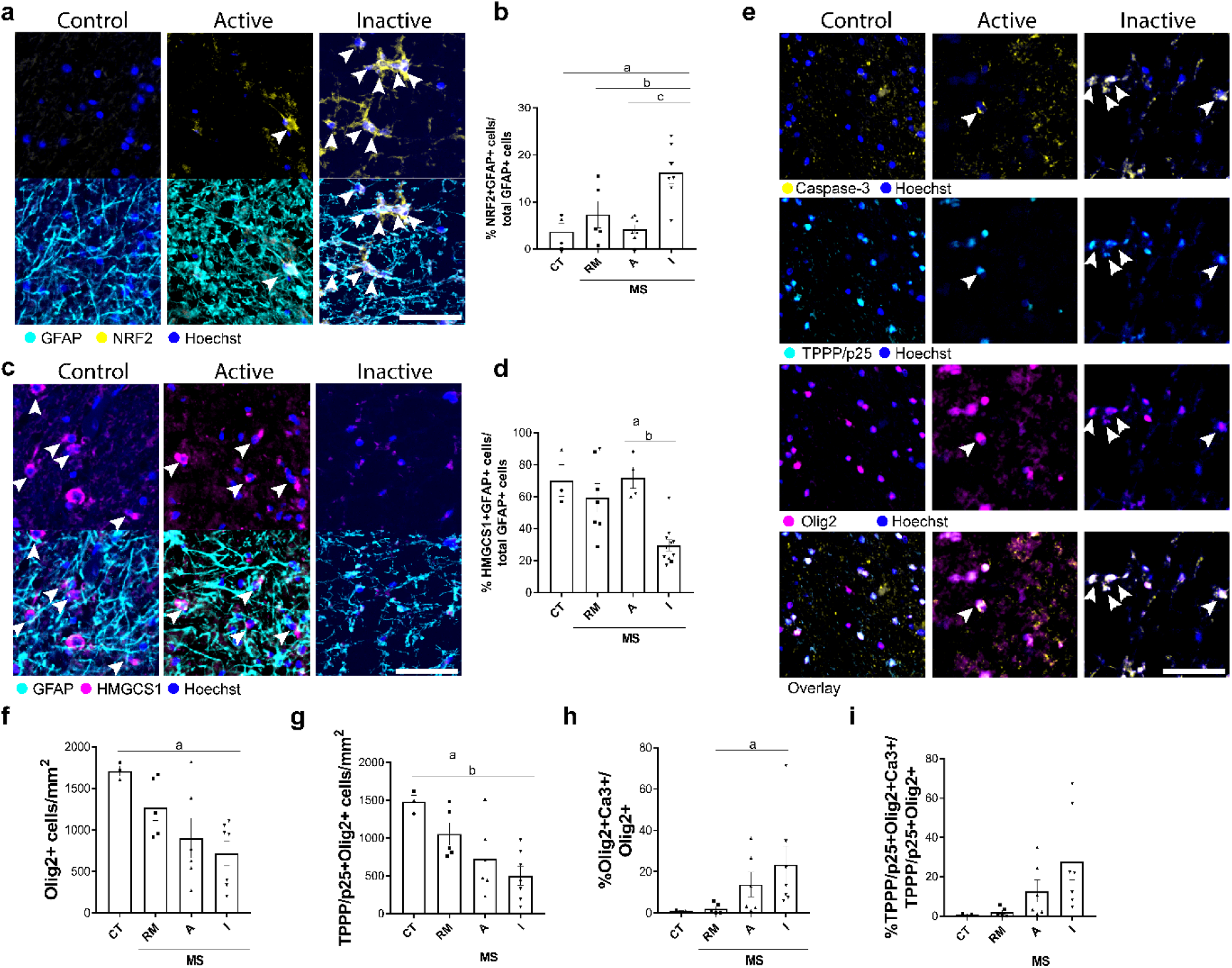
Astrocytic Nrf2 and cholesterol pathways are altered in chronic human brain lesions with poor remyelination potential and oligodendrocyte death. **a.** Astrocytes (GFAP+; cyan) co-stained for Nrf2 (yellow) in controls and active and inactive multiple sclerosis lesions. Hoechst indicates nuclei in blue. Arrows indicate double positive cells. Scale bar; 100 μm. **b.** Mean percentage of GFAP+ cells which are NRF2+ ± s.e.m. in control (CT; n=3), remyelinated (n=6), active (n=6) and inactive lesions (n=7) from multiple sclerosis (MS) cases. One-way ANOVA with Tukey’s multiple comparisons test; ^a^*P*=0.0034, ^b^*P*=0.0285, ^c^*P*=0.011. ANOVA summary (F=8.914, *P*-value=0.0007). **c.** Astrocytes (GFAP+; cyan) co-stained for HMGCS1 (magenta) in controls and active and inactive MS lesions. Hoechst indicates nuclei in blue. Arrows indicate double positive cells. Scale bar; 100 μm. **d.** Mean percentage of GFAP+ cells which are HMGCS1+ ± s.e.m. in control (CT; n=3), remyelinated (n=7), active (n=4) and inactive lesions (n=11) from MS cases. Kruskal-Wallis and Dunn’s multiple comparisons test, ^a^*P=*0.0429, ^b^*P*=0.0113. ANOVA summary (*P*-value=0.0009). **e.** Oligodendrocytes (TPPP/p25+; cyan, and Olig2+; magenta) expressing active caspase-3 (yellow), counterstained with Hoechst (blue). Positive cells indicated with arrowheads. Scale bar, 100 μm. **f.** Mean Olig2+ cells/mm^2^ ± s.e.m. in CT, and RM, A, I MS lesions. One-way ANOVA and Tukey’s multiple comparison test, ^a^*P*=0.0177. ANOVA summary (F=4.472, *P*-value=0.0173). **g**. Mean TPPP/p25+ Olig2+ cells/mm^2^ ± s.e.m. in CT, and RM, A, I MS lesions. One-way ANOVA with Tukey’s multiple comparison test, ^a^*P*=0.0401, ^b^*P*=0.0053. ANOVA summary (F=6.009, *P*-value=0.0055). **h**. Mean percentage of Olig2+ cells expressing active caspase-3/mm^2^ ± s.e.m. in CT, and RM, A, I MS lesions. Kruskal-Wallis test, ^a^*P*=0.0251. ANOVA summary (*P*-value=0.0090) **i**. Mean percentage of TPPP/p25+ Olig2+ cells expressing active caspase-3/mm^2^ ± s.e.m. in CT, and RM, A, I MS lesions. ANOVA with Tukey’s multiple comparison test, no significant. ANOVA summary (F=3.112, *P*-value=0.0539).

Given that our experimental findings suggest that sustained Nrf2 activation and poor cholesterol pathway activation in astrocytes leads to oligodendrocyte death in demyelinated lesions, we asked whether the association could be also made in MS lesions. Indeed, inactive lesions had reduced numbers of oligodendrocyte lineage cells (Olig2+) and mature oligodendrocytes (TPPP/p25+), and increased proportions of apoptotic (active Caspase-3+) Olig2+ or TPPP/p25+ cells (Fig.6e-i). These findings are consistent with oligodendrocyte death recently being shown to contribute to poor remyelination in inactive MS lesions^2^. Therefore, our results propose a link between dysregulated Nrf2 and cholesterol pathway activation in astrocytes with oligodendrocyte death and poor remyelination in humans.

### Luteolin restores oligodendrocyte survival and remyelination efficiency

Given that sustained Nrf2 activation in astrocytes is associated with remyelination failure in mouse and human, we next sought to identify a drug candidate that could target this pathway to restore the efficiency of remyelination. We tested Luteolin, which is BBB-permeable, has an established safety profile in humans^40,41^ and potently decreases Nrf2 levels when it is over-activated (e.g. in tumours or in gain-of-function mutations^40,42,43^), as observed in our *in vitro* assays (Fig.5). Luteolin was administered to lesioned GFAP-Nrf2 mice when Nrf2 activation in wildtype astrocytes normally declines (4-7 DPI) (Fig.7a); this decreased the proportion of GFAP+ cells with active nuclear Nrf2 in comparison to vehicle control (Fig.7b-c). Luteolin restored astrocyte responses at 7 DPI, with a recovery in GFAP+ cells (Fig.7b,d-e), a reduction in the proportion of GFAP+ active-Caspase-3+ cells (Fig.7d,f), and an increase in GFAP+ cells expressing the cholesterol biosynthesis enzyme HMGCS1 (Fig.7g-h). Luteolin may have off-target effects on other cell types; although Luteolin also decreased Nrf2 activation in microglia, this did not significantly impact their expression of HMGCS1, their numbers, or their phagocytosis of myelin debris (Extended Data Fig.16a-e). Direct treatment of primary microglia cultures did not impact their numbers or response to pro-inflammatory stimuli (Extended Data Fig.16f).

**Figure 7.**
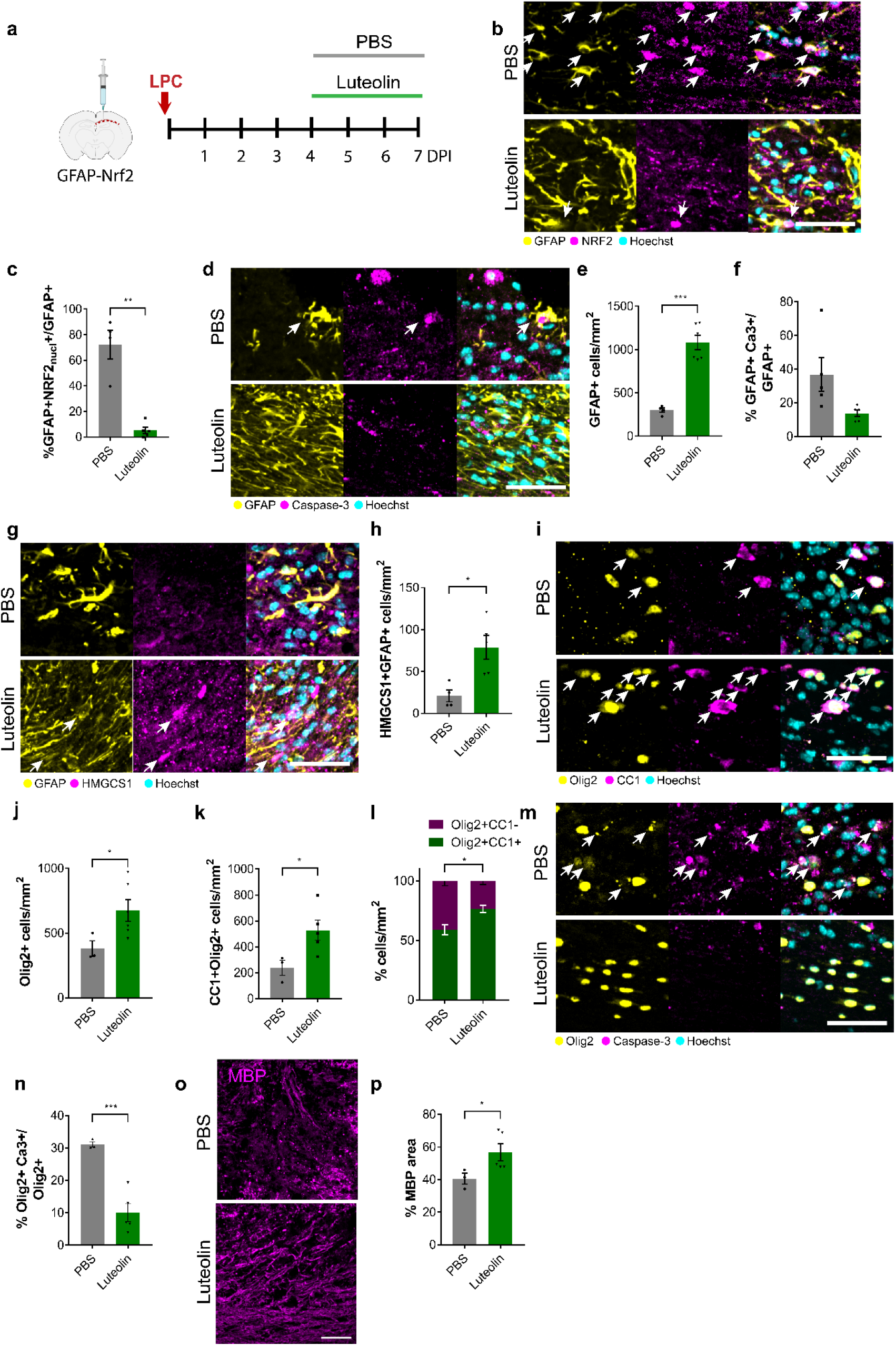
Luteolin restores oligodendrocyte survival and remyelination when astrocytic Nrf2 is sustained. **a.** Focal LPC-demyelination was performed in adult GFAP-Nrf2 corpus callosum and Luteolin or PBS control was administered daily from 4-7 DPI. **b.** Astrocytes (GFAP+; yellow) co-stained for the Nrf2 (magenta) in PBS or Luteolin-treated GFAP-Nrf2 mice at 7 DPI. Arrows indicate double positive cells. Hoechst indicates nuclei in cyan. Scale bar; 50μm. **c.** Mean percentage of GFAP+ cells which have nuclear NRF2 (NRF2_nucl_) ± s.e.m. in PBS or Luteolin-treated GFAP-Nrf2 mice at 7 DPI. 2-tailed unpaired Student’s *t*-test with Welch’s correction, \*\**P=*0.0075, t=5.851. n=3-5 mice/condition. **d**. Astrocytes (GFAP+; yellow) expressing active caspase-3 (magenta), counterstained by Hoechst (cyan) in PBS or Luteolin-treated GFAP-Nrf2 mice at 7 DPI. Double positive cell indicated by arrow. Scale bar, 50 μm. **e.** Mean GFAP+ cells/mm^2^ ± s.e.m. in PBS or Luteolin-treated GFAP-Nrf2 mice at 7 DPI. 2-tailed unpaired Student’s *t*-test with Welch’s correction, \*\*\**P=*0.0001, t=8.925. n=4-5 mice/condition. **f.** Percentage of GFAP+ cells which are positive for active caspase-3/mm^2^ ± s.e.m. in PBS or Luteolin-treated GFAP-Nrf2 mice at 7 DPI. 2-tailed unpaired Student’s *t*-test with Welch’s correction, *P=*0.0831, t=2.245. n=5 mice/condition. **g.** GFAP+ astrocytes (yellow) co-stained with HMGCS1 (magenta) in PBS or Luteolin-treated GFAP-Nrf2 mice at 7 DPI. Hoechst indicates nuclei in cyan. Double positive cells indicated with arrows. Scale bar; 50 μm. **h.** Mean HMGCS1+GFAP+ cells/mm^2^ ± s.e.m in PBS or Luteolin treated GFAP-Nrf2 mice at 7 DPI. 2-tailed unpaired Student’s *t*-test with Welch’s correction, \**P=*0.0116, t=3.642. n= 4-5 mice/condition. **i.** Oligodendrocyte lineage cells (Olig2+; yellow) which are mature (CC1+; magenta) in PBS or Luteolin-treated GFAP-Nrf2 mice at 7 DPI. Hoechst indicates nuclei in cyan. Arrows indicate double positive cells. Scale bar; 50μm. **j.** Mean Olig2+ cells/mm^2^ ± s.e.m. in PBS or Luteolin-treated GFAP-Nrf2 mice at 7 DPI, 2-tailed unpaired Student’s *t*-test with Welch’s correction, \**P=*0.0253, t=2.841. n=3-5 mice/condition. **k.** Mean CC1+Olig2+ cells/mm^2^ ± s.e.m. in PBS or Luteolin-treated GFAP-Nrf2 mice at 7 DPI, 2-tailed unpaired Student’s *t*-test with Welch’s correction, \**P=*0.0260, t=2.938. n=3-5 mice/condition. **l.** Proportion of Olig2+ cells which were CC1+ (green) or CC1-(magenta) ± s.e.m. in PBS or Luteolin-treated GFAP-Nrf2 mice at 7 DPI. Two-way ANOVA with Bonferroni correction PBS vs Luteolin, CC1+Olig2+ *P*-value=0.0088 CC1-Olig2+ *P*-value=0.0086. Two-way ANOVA summary (Interaction F(1,14)= 23.04, *P*-value=0.0003; Row (condition) Factor F(1,14)=4.67×10^−5^, *P*-value=0.9946; Column (type of cell) factor F(1,14) = 93.77, *P-*value<0.0001). n=3-6 mice/condition. **m.** Oligodendrocyte lineage cells (Olig2+; yellow) expressing active Caspase-3 (magenta) in PBS or Luteolin-treated GFAP-Nrf2 mice at 7 DPI. Hoechst indicates nuclei in cyan. Scale bar; 50μm. **n.** Mean percentage of Olig2+ cells which are active Caspase-3+ ± s.e.m in PBS or Luteolin treated GFAP-Nrf2 mice at 7 DPI, 2-tailed unpaired Student’s *t*-test with Welch’s correction, \*\*\**P=*0.0009, t=7.326. n=3-5 mice/condition. **o.** MBP staining (magenta) in the corpus callosum in PBS or Luteolin-treated GFAP-Nrf2 mice at 7 DPI. Scale bar; 50 μm. **p.** Percentage of area of corpus callosum with MBP staining ± s.e.m. in PBS or Luteolin-treated GFAP-Nrf2 mice at 7 DPI. Kolmogorov-Smirnov test, \**P*=0.0357. n=3-5 mice/condition.

We then asked how Luteolin-induced normalization of astrocyte responses impacted the oligodendrocyte lineage and remyelination. Luteolin treatment of GFAP-Nrf2 mice restored numbers of oligodendrocyte lineage cells (Olig2+; Fig.7i-j) and mature oligodendrocytes (CC1+Olig2+; Fig.7i,k), increased the proportion of CC1+ cells (Fig.7l), and decreased the proportion of Olig2+ cells which were active-caspase-3+ (Fig.7m-n). Importantly, Luteolin enhanced remyelination in GFAP-Nrf2 mice, with the percentage of MBP area significantly increased over vehicle control (Fig.7o-p). Luteolin had no impact on the number of oligodendrocyte lineage cells that express Nrf2 or HMGCS1 in the corpus callosum of GFAP-Nrf2 mice, nor did it affect oligodendrocyte lineage cell number or differentiation, when delivered to purified primary cultures (Extended Data Fig.16g-I; Extended Data Fig.16j-k). In addition, the beneficial response to Luteolin *in vivo* was limited to remyelination and not demyelination, as treatment of wildtype mice with Luteolin during the time of LPC-mediated demyelination (0-3 DPI) had no impact on the percentage of MBP area or GFAP+ cells (Extended Data Fig.16l-o). Luteolin may also act on other molecular targets; to address the specificity for Nrf2, we treated mice in which *Nfe2l2* was knocked out in astrocytes with Luteolin, and observed no additional effects on the cholesterol pathway or demyelination, nor was remyelination further increased (Extended data Fig.13t-ai), suggesting the beneficial impact of Luteolin on remyelination occurred primarily via Nrf2 modulation. In summary, we identified Luteolin as a candidate therapy to correct the dysregulated Nrf2 activation in astrocytes in the context of chronic myelin deficit, and to restore oligodendrocyte survival and remyelination efficiency.

## Discussion

Our study identifies astrocyte-oligodendrocyte interaction as a key determinant of CNS myelin regeneration (Extended Data Fig.17), complementing growing evidence for glial-glial interactions regulating myelin health^4–8^. Our finding that astrocytes support the survival of mature remyelinating oligodendrocytes has important implications for the field. First, together with mature oligodendrocyte death recently considered to contribute to remyelination failure in chronic human disease^2^, we highlight the importance of expanding the target of current regenerative drug development beyond immature cells of the oligodendrocyte lineage^3^. Second, our discovery that sustained activation of Nrf2 in astrocytes induces oligodendrocyte death and prevents remyelination counters the widely held concept that Nrf2 activation is often neuroprotective, and is consistent with myelin pathology observed in humans harbouring gain-of-function Nrf2 gene mutations^40^. This calls for further investigation into the impact of existing Nrf2-stimulating drugs on remyelination in progressive chronic disease. The protective function of astrocytic Nrf2 in reducing autoimmune-mediated myelin damage^19,44,45^ suggests distinct molecular requirements for astrocytes to support myelin health during active damage, versus during regeneration once the damage has been inflicted. Third, our discovery that astrocyte cholesterol efflux regulates oligodendrocyte survival and hence remyelination highlights oligodendroglial cholesterol uptake as a potential critical regulator of white matter health. Consistent with this finding, cholesterol is known to support oligodendrocyte survival^20^ and myelin production^46–49^ in development, and remyelination in adulthood^49,50^. We propose that this occurs via astrocyte-mature oligodendrocyte interaction in late remyelination, building on recent work demonstrating that microglia-derived cholesterol precursor influences OPC differentiation in early remyelination^50^ – when we showed astrocyte-derived cholesterol synthesis to be low. Importantly, the cholesterol biosynthesis pathway is dysregulated in the context of chronic demyelination in mice^49,50^ and humans^49^, yet is impacted both positively and negatively by drugs in MS clinical trials^48,51^. We now propose that a strategy to restore the cholesterol biosynthesis pathway in chronic disease is to correct the sustained activation of Nrf2 in astrocytes using Luteolin, to re-establish healthy astrocyte-oligodendrocyte interaction and remyelination efficiency. The established safety profile of Luteolin in humans^52–54^ and its availability as a dietary supplement^52,54^ highlight it as a promising novel regenerative therapy, in addition to drugs in clinical trial which directly impact the oligodendrocyte lineage^49,55–57^.

Our work suggesting the importance of astrocyte-oligodendrocyte interactions in remyelination complements earlier work implicating other cell types (e.g. microglia, regulatory T cells, pericytes) and pathways (e.g. muscarinic receptor and estrogen receptor signalling) in regulating OPC differentiation into remyelinating oligodendrocytes^6,8,16,56–59^. These findings, taken together with our study, point to the possibility of combining therapeutic approaches targeting oligodendrocyte generation and survival for maximal remyelination efficiency. We put forward that targeting astrocyte-oligodendrocyte interaction is a promising strategy to promote CNS remyelination in chronic disease.

## Methods

### Animals

Experiments were performed under a United Kingdom Home Office project licence issued under the Animals (Scientific Procedures) Act 1968. Mice were housed in IVCs in groups of 5 in a 12:12h light: dark cycle with unrestricted access to food and water. C57Bl/6J wildtype mice, *Aldh1l1*-EGFP/Rpl10a mice (stock no.030248), C57BL/6-*Nfe2l2*tm1.1Sred/SbisJ (stock no. 025433), and B6.Cg-Tg(Gfap-TK)7.1Mvs/J (stock no.005698) were purchased from Jackson Laboratories. GFAP-Nrf2.2 mice were generated as previously described^27^ and bred in-house with C57Bl/6 mice. Wildtype littermates from the same colony were used as controls. For *in vitro* studies, Sprague-Dawley wildtype rats were used. Genotyping was performed from ear punches using Transnetyx services.

### *In vivo* focal demyelination

2-3 month old male mice were anaesthetised with isoflurane, administered analgesia (buprenorphine at 0.1 mg/kg and carprofen at 20 mg/kg, subcutaneous injection) then demyelinating lesions were induced by stereotaxically injecting 2 μl of 2 μg lysolecithin (LPC; egg yolk, Sigma Aldrich Cat. No. L4129-100MG) diluted in phosphate-buffered saline (PBS) into the corpus callosum. Sham lesions were induced by injecting PBS. Mice were sacrificed for immunofluorescence analysis by intracardially perfusing with 4% paraformaldehyde (PFA) at 3, 7, 10, 14, and 21 days post-injection (DPI), or by carbon dioxide (CO_2_) asphyxiation at 3, 7, 10 DPI for translational ribosome affinity purification (TRAP). For the former, brains were post-fixed overnight with 4% PFA, cryoprotected in sucrose and sectioned at 12 μm thickness. The ABCA1 agonist CS-6253 (Biosynthesis Inc.)^60^ was administered to demyelinated GFAP-Nrf2 mice from 7-14 DPI. CS-6253 was dissolved in PBS and injected intraperitoneally (i.p.) daily at 30 mg/kg as previously described^21^. The Nrf2 inhibitor Luteolin^42^ was administered to demyelinated GFAP-Nrf2 mice from 4-7 DPI. Luteolin (Tocris Cat. No. 2874) was dissolved in 0.4 ml of PBS containing 1% dimethyl sulfoxide (DMSO, Sigma-Aldrich) to 500μM which was administered daily by i.p. injection at 22.9 mg/kg.

### *In vivo* cuprizone model of de- and remyelination

For assessment of astrocyte reactivity, numbers, proliferation, and expression of proteins in the Nrf2 and cholesterol biosynthesis pathways, male C57Bl/6 mice were fed with 0.3% cuprizone (Sigma Aldrich) mixed in powdered chow for 6 weeks, then fed with normal chow for weeks 7-10. Control mice were fed powdered chow without cuprizone. To ablate reactive astrocytes in the cuprizone model, GFAP-thymidine kinase mice (Jackson Laboratories) were used in which thymidine kinase from the herpes simplex virus is driven by the GFAP promoter. Mice were fed 0.2% cuprizone for 5 weeks then returned to normal chow for 1 week. Either ganciclovir (200 mg/kg at first injection, then 25 mg/kg in 200 μl of PBS) or PBS was administered to both control and transgenic mice from weeks 4-6 by daily i.p. injection.

### Conditional knockout of *Nfe2l2* in astrocytes

4-5 month-old male C57BL/6-*Nfe2l2*tm1.1Sred/SbisJ mice were anaesthetized with isoflurane inhalation. 2.1 x 1010 genome copies of AAV5-GFAP(0.7)-EGFP-T2A-iCre (Vector Labs, Cat. No. VB1131) or AAV5-GFAP(0.7)-EGFP (Vector Labs, Cat. No. VB1149) were stereotactically injected into the corpus callosum at an infusion rate of 200 nl/ml. Recombination was allowed for 5 days before inducing demyelination with LPC, as above, and sacrificed 3 or 7 days post-LPC. A subset of AAV5-GFAP(0.7)-eGFP-T2A-iCre-treated *Nfe2l2*-floxed mice were treated daily with Luteolin as above from the time of LPC injection to 3 or 7 days post-LPC.

### Immunofluorescence of *in vivo* tissue sections

Paraffin-embedded sections of brains from cuprizone-fed mice were pre-heated at 60°C for 20 min, immersed in Histoclear II and progressively rehydrated in descending ethanol gradients (100, 95, 70, 50 %) and water. Frozen sections of brains from LPC-injected mice were air dried for 15 min. Heat induced antigen retrieval was achieved by microwaving sections at medium power in 10 mM citrate buffer (pH 6.0) and heating for 20 min at 60°C, followed by PBS washes. Slices were blocked for 1 h (5% horse serum and 0.3 % Triton-X in PBS), then incubated with primary antibodies overnight at 4°C in a humid chamber. Antigen retrieval for myelin proteins such as MBP and MAG was achieved by immersing the slides in cold methanol −20°C for 20 minutes, then washing with PBS. The following antibodies were used: rat anti-myelin basic protein (MBP, 1:100, Bio-Rad antibodies MCA409S), mouse anti-myelin-associated glycoprotein (MAG, 1:100, SigmaAldrich MAB1567),chicken anti-glial fibrillary acidic protein (GFAP, 1:500, BioLegend PCK-591P), chicken anti-GFAP (1:500, Cambridge Bioscience 829401), rabbit anti-SOX9 (1:500, Sigma-Aldrich AB5535), mouse anti-Nestin (1:100, Abcam ab6142), mouse anti-vimentin (1:100, Sigma-Aldrich clone LN-6 MAB1681), rabbit anti-nuclear factor IA (NFIA, 1:250, Abcam ab228897), rabbit anti-Ki67 (1:500, Sigma-Aldrich AB9260), mouse anti-APC (CC1, 1:100, Abcam ab16794), rabbit anti-Olig2 (1:100, Sigma-Aldrich AB9610), rabbit anti-HMOX1 (1:100, Enzo ADI-SPA-895-D), rabbit anti-HMGCS1 (1:500, Invitrogen PA5-29488), rabbit anti-FDPS (1:500, Invitrogen PA5-28228), anti-MVD antibody (1:100, Abcam ab198823) and anti-FDFT1 (1:100, Abcam ab236666), rabbit anti-IBA1 (1:500, Abcam 178846), rabbit anti-ABCA1 (1:100, Novus Biologicals NB400-105), mouse anti-Sox10 (1:100, Sigma-Aldrich AMAB91297), mouse anti-Olig1 (1:1000, EMD Millipore MAB5540), rabbit anti-active Caspase-3 (1:500, BD Pharmigen™ 559565), mouse anti-CNPase (1:2000, Sigma-Aldrich AMAB91072), mouse anti-MASH1 (1:100, BD Bioscience 556604), rabbit anti-NQO1 (1:100, Abcam ab2346), rat anti-NRF2 (1:500, Cell Signalling 14596S), and chicken anti-GFP (1:100, Abcam ab13970). Following washes with PBS, Alexa secondary antibodies (1:500, Life Technologies) and Hoechst (Sigma-Aldrich) were applied for 2h at 20-25°C in a humid chamber. Slides were coverslipped with Fluoromount G (Cambridge Biosciences). Z-stacks of images were obtained using an Olympus 3i Spinning Disk confocal microscope (30x silicone objective) and SlideBook 6 software.

### Magnetic cell separation (MACS), RNA extraction, and RNA sequencing

Brains were dissociated using gentleMACS^®^Dissociator, and astrocytes were isolated using the Anti-ACSA-2 MicroBead Kit (Miltenyi Biotec). RNA-sequencing was performed using TrueSeq Stranded Total RNA V2 library preparation along with next-generation sequencing on the Illumina Novaseq 6000 platform with an approximate read-depth of 90 million reads per sample. At-least 1 μg RNA per sample was utilised, with RNA-integrity number (RIN) > 7.

### Translational ribosome affinity purification (TRAP), RNA extraction, and RNA sequencing

Corpus callosa were dissected from LPC-injected and non-lesioned mice and were processed for TRAP as done previously^17^. RNA was extracted from pre-TRAP controls (‘Input’) and TRAP samples using the Agilent Nanoprep kit. Quality of the samples was measured using an Agilent Bioanalyzer and the Agilent RNA 6000 PICO assay protocol. Samples selected had an RNA integrity number (RIN) >7, and sent to Cambridge Genomic Services for Next-Generation Sequencing. For library preparation, the SMART-Seq v4 Ultra Low input RNA kit (Takara Bio USA) was used according to the manufacturer’s instructions. The sequencing was run on a NextSeq 550 system (Illumina) on a 150-cycle high-output run with paired-end reads (30 million reads). For each TRAP sample, 75 base pair paired-end reads were mapped to the primary assembly of the mouse (mm10) reference genome contained in Ensembl release 93. Alignment was performed with STAR version 2.5.3a. Per-gene read counts were summarised using featureCounts version 1.5.2. For read-mapping and feature counting, genome sequences and gene annotations were downloaded from Ensembl version 94. Differential expression analysis was then performed using DESeq2 (R package version 1.18.1), with a significance threshold calculated at a Benjamini–Hochberg-adjusted P value of <0.05.

### Differential expression and pathway analysis

Differential expression analysis was performed using DE-Seq2 (R package version 1.18.1), with a significance threshold calculated at a Benjamini–Hochberg-adjusted P value of <0.05. In order to account for variable amounts of background mRNA contamination in TRAP-seq samples, the presence of a set of cell-type-specific genes which should not be expressed in astrocytes were quantified. For each cell-type-specific gene, its expression in FPKM (fragments per kilobase per million mapped reads) in each sample was divided by its maximum FPKM in any sample, giving for each gene a per-sample measure of background contamination (with a value between 0 and 1). Per-gene contamination values for the six genes correlated well across samples. An average was taken over the six genes to give a persample contamination measure (C) with a value between 0 and 1. For every other gene, the correlation of its gene expression with C over all samples was calculated, giving a per-gene measure (R) of the likelihood of a gene’s apparent presence in the data as being due to background mRNA contamination. To determine how likely it was to get high values of R by chance, a test gene expression dataset containing the FPKM values from our data was created with the values scrambled between samples for each gene (in which case, this test gene expression should not be correlated with C). From this test data, the R value for each scrambled gene was created, deriving a null distribution of R values. In the real dataset, a Z-score for each gene indicating how far its own R value was from the mean of this null distribution in units of standard deviations of the null distribution was calculated. Any gene with a Z-score greater than 2 was excluded from analyses.Pathway analysis was performed using Ingenuity Pathway Analysis software (QIAGEN). Genes were inputted with their FPKM (cut off >5 FPKM) and corresponding log2FC in lesions versus control. Top canonical pathways were considered significant at *P* values <0.05. Gene ontology (GO) and pathway networks were analysed using the plugin ClueGO v.2.5.4 for Cytoscape v.3.7.1 software, with the top 200 significantly upregulated genes at each time point versus control used for analysis. Statistical significance was analysed by the two-sided hypergeometric test and Bonferroni step down correction, and grouping was based on highest significance and GO-term fusion.

### Proteomics Sample Preparation, Mass Spectrometry and Analysis

Frozen corpus callosa were pulverised using the Precellys Bioruptor and bead-based lysis kit (VWR). Tissue was transferred to 1.5ml Precellys beaded tubes with 300 μl of lysis buffer (5% SDS, 10 mM TCEP, 50 mM TEAB) before being loaded into Precellys Bioruptor for a 3 min cycle at 5500 rpm at 4°C. Lysed tissue was transferred to 1.5 ml lo-bind eppendorfs and boiled at 95°C for 5 min before DNA digestion with Benzonase at 37°C for 15 min. Samples were alkylated with 20 mM iodoacetamide for 1 h at 22°C. Protein concentration was determined using EZQ protein quantitation kit (Invitrogen) as per the manufacturer instructions. Protein isolation and clean up was performed using S-TRAP™ (Protifi) columns before digestion with trypsin at 1:20 enzyme:protein ratio for 2 h at 47°C. Digested peptides were eluted from S-TRAP™ columns using 50 mM ammonium bicarbonate, followed by 0.2% aqueous formic acid and 50% aqueous acetonitrile containing 0.2% formic acid. Eluted peptides were dried down overnight before resuspension in 5% formic acid ready for injection onto the Orbitrap Exploris (Thermo Fisher) mass spectrometer using data independent acquisition (DIA). Peptides (2 μg/ sample) were injected onto a nanoscale C18 reverse-phase chromatography system (UltiMate 3000 RSLC nano, Thermo Scientific) and electrosprayed into an Orbitrap Exploris Mass Spectrometer (Thermo Fisher). For liquid chromatography the following buffers were used: buffer A (0.1% formic acid in Milli-Q water (v/v)) and buffer B (80% acetonitrile and 0.1% formic acid in Milli-Q water (v/v). Samples were loaded at 10 μL/min onto a trap column (100 μm × 2 cm, PepMap nanoViper C18 column, 5 μm, 100 Å, Thermo Scientific) equilibrated in 0.1% trifluoroacetic acid (TFA). The trap column was washed for 3 min at the same flow rate with 0.1% TFA then switched in-line with a Thermo Scientific resolving C18 column (75 μm × 50 cm, PepMap RSLC C18 column, 2 μm, 100 Å). Peptides were eluted from the column at a constant flow rate of 300 nl/min with a linear gradient from 3% buffer B to 6% buffer B in 5 min, then from 6% buffer B to 35% buffer B in 115 min, and finally to 80% buffer B within 7 min. The column was then washed with 80% buffer B for 4 min and re-equilibrated in 3% buffer B for 15 min. Two blanks were run between each sample to reduce carry-over. The column was kept at a constant temperature of 50°C. The data was acquired using an easy spray source operated in positive mode with spray voltage at 2.445 kV, and the ion transfer tube temperature at 250°C. The MS was operated in DIA mode. A scan cycle comprised a full MS scan (m/z range from 350-1650), with RF lens at 40%, AGC target set to custom, normalized AGC target at 300, maximum injection time mode set to custom, maximum injection time at 20 ms, microscan set to 1, and source fragmentation disabled. MS survey scan was followed by MS/MS DIA scan events using the following parameters: multiplex ions set to false, collision energy mode set to stepped, collision energy type set to normalized, HCD collision energies set to 25.5, 27 and 30 %, orbitrap resolution 30000, first mass 200, RF lens 40 %, AGC target set to custom, normalized AGC target 3000 %, microscan set to1, maximum injection time 55 ms. Data for both MS Scan and MS/MS DIA Scan events were acquired in profile mode. Raw mass spectrometry data was processed using Spectronaut (Biognosys) version 14.10.201222.47784 with the DirectDIA option. The following parameters were selected: cleavage rules were set to Trypsin/P, maximum peptide length 52 amino acids, minimum peptide length 7 amino acids, maximum missed cleavages 2 and calibration mode automatic. Carbamidomethylation of cysteine was set as a fixed modification while the following variable modifications were selected: oxidation of methionine, deamidation of asparagine and glutamine and acetylation of the protein N-terminus. The FDR threshold for both precursor and protein was set at 1%. Profiling was disabled. DirectDIA data were searched against a mouse database from Uniprot release 2020 06. This database consisted of all manually annotated mouse SwissProt entries along with mouse TrEMBL entries with protein level evidence and a manually annotated homologue within the human SwissProt database. Fold changes in protein abundance were calculated using normalised protein intensities. Normalisation was performed by dividing the intensity for each protein by the summed protein intensity for all proteins identified within an individual sample.

### Primary neural cell cultures

Mixed glial cultures were derived from the cortices of P0-P3 Sprague-Dawley rats of both sexes. Microglia were isolated by collecting the floating fraction of 10 day-old mixed glial cultures following 1 h on a rotary shaker at 250 rpm at 37°C, and plated on poly-D-lysine-coated 16-well glass chamberslides (Lab-TEK) at 5×10^4^ cells per well in Dulbecco’s Modified Essential Media (DMEM) with glucose (4.5 g/L), l-glutamine, pyruvate, 10% fetal calf serum, and 1% penicillin/streptomycin. Microglia were either left untreated or treated with IFNγ (20ng/ml) and LPS (0127:B8, 100 ng/ml) overnight, and Luteolin or vehicle control. Oligodendrocyte progenitor cells (OPCs) were isolated from mixed glial cultures following overnight agitation following microglia depletion as above, with astrocyte depletion in the floating fraction achieved by differential adhesion. OPCs were plated at 2×10^4^ cells per well in PDL-coated plastic chamberslides (Lab-TEK) in DMEM containing glucose (4.5 g/L0, L-glutamine, pyruvate, SATO (16 μg/ml putrescine, 400 ng/ml L-thyroxine, 400 ng/ml triiodothyroxine, 60 ng/ml progesterone, 5 ng/ml sodium selenite, 100 μg/ml bovine serum albumin fraction V, 10 μg/ml insulin, 5.5 μg/ml halo-transferrin (all from Sigma-Aldrich)), 0.5% fetal calf serum (GIBCO), 1% penicillin/streptomycin. For 2 days, cells were exposed to 10 ng/ml platelet-derived growth factor and 10 ng/ml fibroblast growth factor-2. OPCs were matured to oligodendrocytes by withdrawal of growth factors from media for 5 days. Cells were treated with unconditioned astrocyte media or astrocyte conditioned media (ACM) in a 1:1 ratio with OPC media treated with Luteolin (Tocris) or vehicle control (DMSO) for 2 days.

Astrocytes were detached from flasks following microglia and OPC depletion with 0.25% Trypsin-EDTA for 5-10 min at 37°C. Following trypsin inactivation with fresh media, cells were separated by gentle pipetting and plated in 16-well glass chamberslides (Nunc) at 5×10^4^ cells/well in DMEM with 10% FBS and 1% penicillin/streptomycin. 1 day post-plating, astrocytes were either left untreated or treated with CDDO^TFEA^ (100 nM). 24 h later, CDDO^TFEA^-treated astrocytes were washed thoroughly with fresh media and a subset of experiments involved subsequent treatment with Luteolin (50 μM) or CS-6253 (1 μM). BODIPY-FL-C12 (2 μM) was fed to astrocytes in all conditions for 6 hours then wells were washed thoroughly with fresh media to remove any residual compound not taken up by astrocytes; for experiments where astrocyte conditioned media was to be applied to oligodendrocytes, serum was halved to 5%. After 18 hours, astrocyte conditioned media was collected and applied to oligodendrocytes, OPCs, or brain explants at a ratio of 1:1 with respective culture media.

### Quantitative reverse transcription real-time polymerase chain reaction

Primary astrocytes were washed with PBS then incubated for 10 min with RNA lysis buffer. RNA extraction was achieved using the Agilent Nanoprep kit according to the manufacturer’s instructions. Complementary DNA (cDNA) was generated using SuperScriptTM VILOTM cDNA synthesis kit (Invitrogen). The FastStart Universal SYBR Green Master mix (ROX; Roche) was used on a QuantStudio 5 Real-Time polymerase chain reaction (PCR) system. Primers used were as follows: *Hmgcs1* (5’-ATG GGG CTC GTG CAT AGT AA-3’ and 5’-ACT CTC AGT GCT CCC CGT TA-3’); *Fdps* (5’-GCA CTG ACA TCC AGG ACA AC-3’ and 5’-AGC CAC TTT TTC TGG GTC CT-3’); *Mvk* (F: 5’-CTCAAGGACGGGGTCTCC −3’ and R: 5’-GGCCCACTTGTTGATTGACT-3’); *Fdft1* (5’-TCC CTG ACG TCC TCA CCT AC-3’ and 5’-CCC CTT CCG AAT CTT CAC TA-3’); *Nrf2* (5’-CAGCTCAAGGGCACAGTGC-3 and 5’-GTGGCCCAAGTCTTGCTCC-3’); *Gclc* (5’-CCAACCATCCGACCCTCTG-3’ and 5’-TGTTCTGGCAGTGTGAATCC-3’); *Nqo1* (5’-CCTTCCGAGTCATCTCTAGC-3’ and 5’-AGCAAGGTCTTCTTATTCTGGA-3’); *Gapdh* (F: 5’-GGGTGTGAACCACGAGAAAT-3’ and R: 5’-CCTTCCACAATGCCAAAGTT-3’); *Aldh1l1* (F: 5’-CTTTGACCTTGGGTGCCT-3’ and R: 5’-ATCTGCTTTCCCATCCTTGT-3’).

### Immunocytochemistry

Cells were fixed with 4% PFA for 10 min then wash in PBS and stored in PBS at 4°C. Cells were blocked for 1 hour and incubated with primary antibodies at 4°C overnight, washed in PBS, then incubated with secondary antibodies for 1 h at room temperature prior to counterstaining with Hoechst and coverslipping with Fluoromount-G. Primary antibodies included rat anti-Nrf2 (1:500, Cell Signalling 14596S), rabbit anti-Aldh1l1 (1:500, Abcam ab190298), chicken anti-GFAP (1:1000, Cambridge Bioscience 829401), rat anti-MBP (1:250, Bio-Rad antibodies MCA409S), rabbit anti-TPPP/p25 (1:1000, Abcam ab92305), mouse anti-Sox10 (1:100, Sigma-Aldrich AMAB91297), rabbit anti-IBA1 (1:500, Abcam 178846), and mouse anti-iNOS (1:500, BD Biosciences 610329). Following washes with PBS, Alexa secondary antibodies (1:500, Life Technologies) and Hoechst (Sigma-Aldrich) were applied for 1 h. For cell death detection, a TUNEL assay was performed prior to staining according to the manufacturer’s instructions (Promega G3250).

### *Ex vivo* organotypic mouse brain explants

Cerebellum and hindbrain from P0-P2 mouse pups of both sexes were sagittally sectioned (300 μm) on a McIlwain tissue chopper and plated onto Millipore-Millicell-CM mesh inserts (Fisher Scientific) in 6 well culture plates at 6 explants per brain per insert. Culture media consisted of 50% minimal essential media, 25% heat inactivated horse serum, 25% Earle’s balanced salt solution (all from GIBCO), glucose (6.5 mg/ml, Sigma-Aldrich), 1% penicillin-streptomycin (Life Technologies), 1% Glutamax (Life Technologies) and 1% HEPES (Invitrogen). Demyelination was induced at 14 days *in vitro* by application of lysolecithin (LPC, 0.5 mg/ml, Sigma-Aldrich) for 18 hours, then washed off. Cultures were exposed to primary astrocyte conditioned media (ACM) after demyelination by treatment with 1:1 explant media:ACM or explant media:astrocyte media as a control, from 2 to 5 days post LPC application. In separate experiments, cultures were treated with the ABCA1 antagonist PSC833 (1-10 μM; Tocris Bioscience) or vehicle control (DMSO) from 2 to 7 days post LPC application. Explants were fixed with 4% PFA for 10 minutes, washed with PBS and stored at 4°C until staining. Explants were permeabilized and blocked for 1 h in 5% horse serum and 0.3% Triton-X-100 in PBS, and primary antibodies were applied overnight at 4°C with gentle shaking in a humid chamber, which included rat anti-MBP (1:250; AbD Serotec, MCA409S) and mouse anti-neurofilament (1:1000; EnCor, MCA-9B12). Alexa secondary antibodies (1:500, Life Technologies) were applied for 2 hours then counterstained with Hoechst and mounted with Fluoromount-G (Southern Biotech). Z-stacks of explants were acquired on an Olympus Spinning Disk confocal microscope with Slidebook software 6.

### Human tissue

Post-mortem tissue from multiple sclerosis (MS) patients and control cases that died of non-neurological causes were obtained via a UK prospective donor scheme with full ethical approval from the UK Multiple Sclerosis Tissue Bank (MREC/02/2/39) and their use was in accord with the terms of the informed consents. Diagnosis of MS was confirmed by neuropathological means by F. Roncaroli (Imperial College London) and clinical history was provided by R. Nicholas (Imperial College London). Snap frozen unfixed tissue blocks (2 × 2 × 1 cm^3^) were cut at 10 μm and stored at −80°C. MS lesions were classified according to the International Classification of Neurological Disease using Luxol Fast Blue (LFB) staining and CD68+ immunoreactivity. Healthy control tissue showed intact myelin, no myelin debris, and few CD68+ cells. Fully remyelinated lesions showed intermediate intensity of LFB, little to no myelin debris, and few CD68+ cells. Active lesions showed a diffuse LFB border with demyelination, and abundance of myelin debris throughout the lesion, and high numbers of CD68+ cells. Inactive lesions showed significant demyelination, no myelin debris, tissue destruction, and low CD68+ cell numbers. Tissue blocks from 3 controls and 8 MS cases were analysed. Sections were fixed in 4% PFA for 1 h, washed with Tris-buffered saline (TBS) and permeabilized in methanol for 10 min at −20°C.

Following three washes in 0.001% TritonX-100 in TBS, sections were microwaved at medium power in Vector Unmasking Solution for 5 min, heated for 20 min in the oven at 60°C and cooled down for 10 min. After three washes with 0.001% TritonX-100, sections were blocked for 5 min in Peroxide Bloxall (Vector Labs), then in blocking buffer for 1 h (10% normal horse serum, 0.5% TritonX-100 in TBS). Primary antibodies were sequentially applied overnight at 4°C, and included rabbit anti-HMGCS1 (1:500, Invitrogen PA5-29488), rabbit anti-HMOX1 (1:100, Enzo ADI-SPA-895-D), rabbit anti-TPPP/p25 (1:100, Abcam ab92305), rabbit anti-active Caspase-3 (1:500, BD Pharmigen™ 559565), goat anti-NQO1 (1:100 Abcam ab2346), mouse anti-Olig2 (1:100, EMD Millipore MABN50), chicken anti-GFAP (1:100, Cambridge Bioscience 829401), and rat anti-Nrf2 (1:100, Cell Signalling 14596S). After three washes in 0.001% TritonX-100, sections were incubated with secondary antibody (HRP ImmPress, Vector Labs) for 1 h at room temperature. Following three washes with 0.001% TritonX-100, slides were incubated with Opal fluorophore-conjugated tyramide signal amplification (TSA; 1:100, Akoya Biosciences). After the application of tyramides, slices were microwaved at medium power in Vector Unmasking Solution for 8 min, cooled for 20 minutes then washed in 0.001% TritonX-100. For detection of GFAP, HMOX1 and HMGCS1, Alexa-conjugated secondary antibodies (1:500, Life Technologies) were applied for 2 h after tyramides. On the last secondary antibody application, slides were counterstained with Hoechst and coverslipped with Fluoromount-G. Entire tissue sections were imaged using a Zeiss AxioScan Z.1 SlideScanner at a single z-plane. Three regions of interest per lesion were chosen in a blinded manner, cell densities were analysed (also blinded) and averaged per sample per lesion type.

### Human brain single nuclei RNA sequencing analysis

Published single nuclei RNA sequences from control and multiple sclerosis brains were downloaded from NCBI (Accession number PRJNA544731)^29^. This was based on sequencing of 9 control cases, 8 MS cases with active lesions, and 4 MS cases with inactive lesions. Cellranger (version 3.1.0) count was used to map reads from each sample to GRCh38 using default parameters, and output from each sample was aggregated with Cellranger Aggr. The aggregated filtered feature barcodes matrix was imported into Seurat (version 4.0.4) and filtered based on quality metrics. Cells with fewer than 200 or greater than 2500 RNA features were excluded, as were cells with >15% mitochondrial RNA. Features which were present in <5 cells were discarded. Read counts were normalised using Log1p transformation. Variance stabilizing transformation was used to identify the top 2000 most variable features. Data was scaled to the cell count total and centred, and effects of percent mitochondrial RNA and number of UMIs was regressed out. A shared nearest neighbour graph was constructed with the top 10 principal components from the scaled data to identify clusters. Astrocytes were identified based on expression of astrocytic marker genes AQP4 and GFAP visualized by t-SNE plot. Annotated astrocytes were extracted and re-clustered using the method described above to identify sub-clusters. Non-protein coding genes were removed from cluster gene lists. Cluster gene lists were computed by finding differentially expressed genes between cluster 1 and all other cells using Seurat function FindMarkers, setting minimum percent of cells expressing a feature in either comparison group to 5% and the log fold change threshold to 0. For enrichment of Nrf2-target genes, a minimum percent threshold of gene abundance was set to 5%. Mouse astrocyte transcriptomes from our dataset were compared to published single nuclei sequences from multiple sclerosis lesions using Analysis Match from OmicSoft expression analysis (Ingenuity Pathway Analysis).

### Imaging processing and quantification

ImageJ was used to convert maximum projection images into thresholded 8-bit RGB tiffs for astrocyte and oligodendrocyte counts, and to separate nuclei by watershed segmentation. Thresholded images were used to measure the percentage area of corpus callosum covered by MBP or CNPase staining over total lesion area of the image. Imaris 9.0.0 software was used to quantify astrocytes positive for HMGCS1, FDPS, and HMOX1 in z-stacks using ImarisColoc and ImarisSurfaces, where thresholding was achieved using absolute intensity and the size of each ‘surface’ was adjusted so that each was the size of a cell. Two regions of interest were analysed and averaged per stain, repeated for three sections per animal, from which a mean per animal was obtained. For remyelination index quantification, images were cropped in 5 z-planes and imported into Volocity software (Perkin Elmer), in which the voxels for each channel (NFH and MBP) were measured as well as the co-localisation between both channels. The co-localisation value was normalised to the NFH voxel counts to account for variations in axonal density^61^.

### Statistics

Sample size was calculated by two-sided 95% confidence interval using the normal approximation method using OpenEpi software. Statistical analysis was performed using GraphPad Prism 7 software. The Shapiro-Wilk test was used to assess Gaussian distribution, and then statistical analysis was performed using either parametric tests when normally distributed (2-tailed unpaired Student’s t-test or ANOVA with Tukey’s multiple comparison test), or non-parametric tests when not (Kolmogorov-Smirnov test or Kruskal-Wallis test). When two independent variables were compared (e.g. treatment and time) a two-way ANOVA was applied. Data is represented as mean ± standard error of means (s.e.m.). *P* values <0.05 were considered statistically significant and are indicated in the figure legends.

## Data availability

The data that support the findings of this study are available from the corresponding author on request.

## Acknowledgments

This work was funded by a studentship from the United Kingdom Multiple Sclerosis Society (to V.E.M.; grant no. 54), a Medical Research Council and United Kingdom Multiple Sclerosis Society Career Development Award (to V.E.M.; grant no. MR/M020827/1), a Medical Research Council Senior Non-Clinical Fellowship (to V.E.M.; grant no. MRC/V031260/1), and funds from the Medical Research Council Centre for Reproductive Health (grant no. MR/N02256/1). The cuprizone studies were supported by the German Research Foundation (T.K.; grant no. SFB-TR128-B7) and the Hannover Medical School (Hochschulinterne Leistungsförderung; M.S.). We thank the United Kingdom Multiple Sclerosis Society Tissue Bank for providing tissue. We thank A. Rooney, N. McNamara, S. Kent, L. Ryan, L. Zoupi, S. Jaekel, and M. Blesa for helpful discussions, and F. Roncaroli, R. Nicholas, and C. Watkins for technical support. Diagrams were created using BioRender.com.

## Author contributions

I.M-G co-designed the study, carried out the experiments, analysed and interpreted the data, made the figures, and co-wrote the manuscript. Z.J. assisted in TRAP protocols. O.D. and K.E. performed RNA sequencing analysis. R.K.H., L.F., and A.M. assisted with surgical procedures, tissue processing and cryosectioning, cultures, and immunostaining. A.F.L. and A.H. performed proteomics analysis. V.G., T.S. and M.S. carried out astrocyte depletion experiments. J.A.J. made GFAP-Nrf2 mice. J.A.F. and J.H.F. assisted in breeding and genotyping of transgenic mice. T.K. performed cuprizone model experiments. A.W. provided human tissue pathological mapping. S.C. and G.H. co-supervised the project and contributed to experimental design, interpretation, and manuscript editing. V.E.M. supervised the project, guided experimental design and interpretation, and co-wrote the manuscript.

## Materials and correspondence to

Veronique E. Miron

