## Extended Data for "Astrocyte-Oligodendrocyte interaction regulates central nervous system regeneration"

### Molina-Gonzalez et al:

#### Extended Data Table of Contents

Extended Data Figures & Legends 1-17

Extended Data Table 1

Supplementary Sheet 1 Legend

**Extended Data Figure 1. Reactivity of astrocytes during remyelination.**

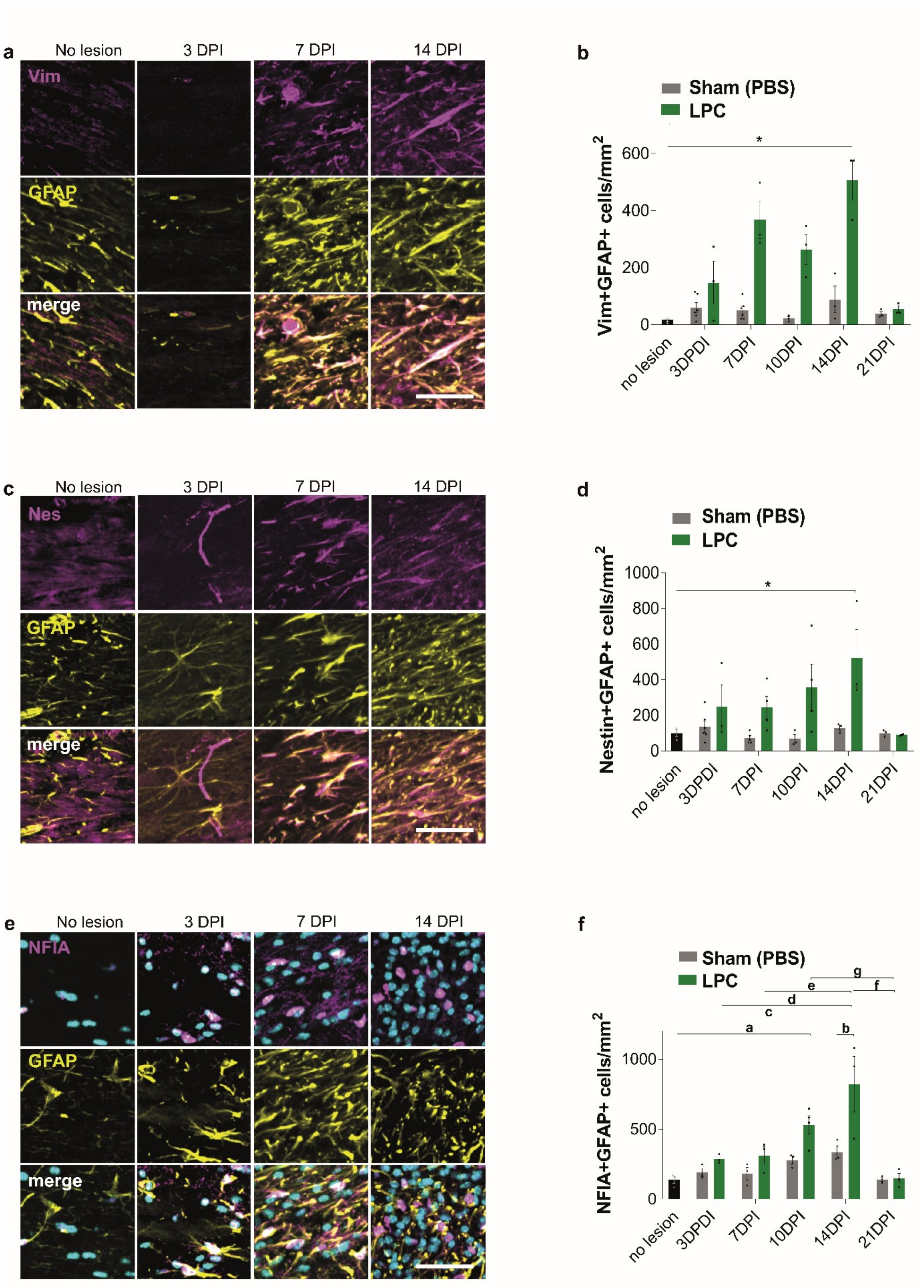

**a.** Staining for GFAP (yellow) and vimentin (Vim) (magenta) in no lesion control and at 3, 7, and 14 DPI. Scale bar; 50 µm.
**b.** Vimentin (Vim)+ GFAP+ cells/mm$^2$ ± s.e.m. in no-lesion and Sham (PBS) controls and at 3, 7, 10, 14, and 21 DPI. Kruskal-Wallis test and Dunn's test correction, **P*=0.0390. ANOVA summary *P*=0.0045. n=3-6 mice/group.
**c.** Staining for GFAP (yellow) and nestin (Nes; magenta) in no-lesion control and at 3, 7, and 14 DPI. Scale bar; 50 µm.
**d.** Nestin+ GFAP+ cells/mm$^2$ ± s.e.m. in no-lesion and Sham (PBS) controls and at 3, 7, 10, 14, and 21 DPI. One-way ANOVA with Tukey's multiple comparison test, **P*=0.0309. ANOVA summary *P*=0.0043 F=3.49. n=3-6 mice/group.
**e.** Staining for GFAP (yellow) and NFIA (magenta) in no-lesion control and at 3, 7, and 14 DPI. Hoechst indicates nuclei in cyan. Scale bar; 50 µm.
**f.** NFIA+ GFAP+ cells/mm$^2$ ± s.e.m. in no-lesion and Sham (PBS) controls and at 3, 7, 10, 14, and 21 DPI. One-way ANOVA and Tukey's multiple comparison test, $^{a}P=0.0151$, $^{b}P=0.0034$, $^{c}P<0.0001$, $^{d}P=0.0011$, $^{e}P=0.0019$, $^{f}P<0.0001$, $^{g}P=0.0185$. ANOVA summary *P*<0.0001 F=8.487. n=3-4 mice/group.

**Extended Data Figure 2. Astrocyte numbers and proliferation during remyelination.**

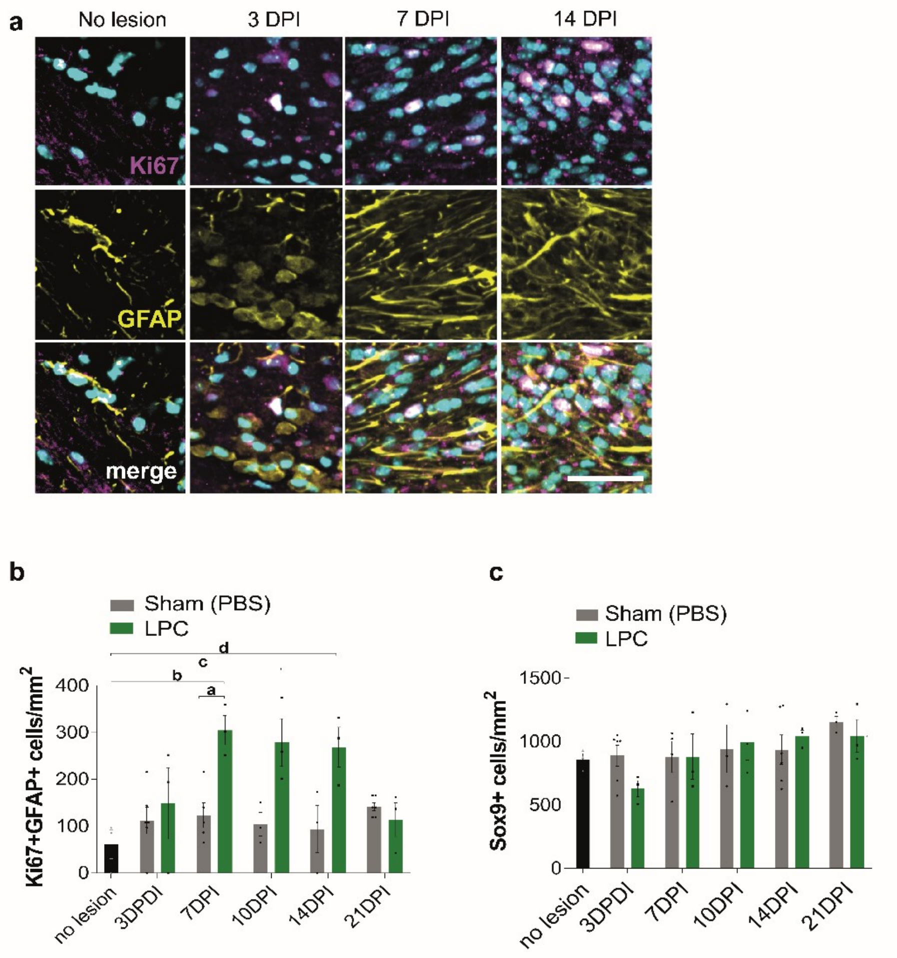


**a.** Proliferative astrocytes identified with Ki67 (magenta) and GFAP+ (yellow) in no-lesion control and at 3, 7, and 14 DPI. Hoechst indicates nuclei in cyan. Scale bar; 100 µm.

**b.** Mean Ki67+GFAP+ cells/mm$^2$ ± s.e.m. in Sham (PBS) and no-lesion control and at 3, 7, 10, 14, and 21 DPI. One-way ANOVA with Tukey's multiple comparison test, $^aP$=0.0490, $^bP$=0.0092, $^cP$=0.0255, $^dP$=0.0410. ANOVA summary $P$=0.0009 F=4.427. n=3-6 mice/group.

**c.** Mean SOX9+ astrocytes/mm$^2$ ± s.e.m. in Sham (PBS) and no-lesion control and at 3, 7, 10, 14, and 21 DPI. Kruskal-Wallis test and Dunn's test correction, not significant, $P$=0.3946. n=3-6 mice/group.

**Extended Data Figure 3. Astrocyte reactivity in the cuprizone model of remyelination.**

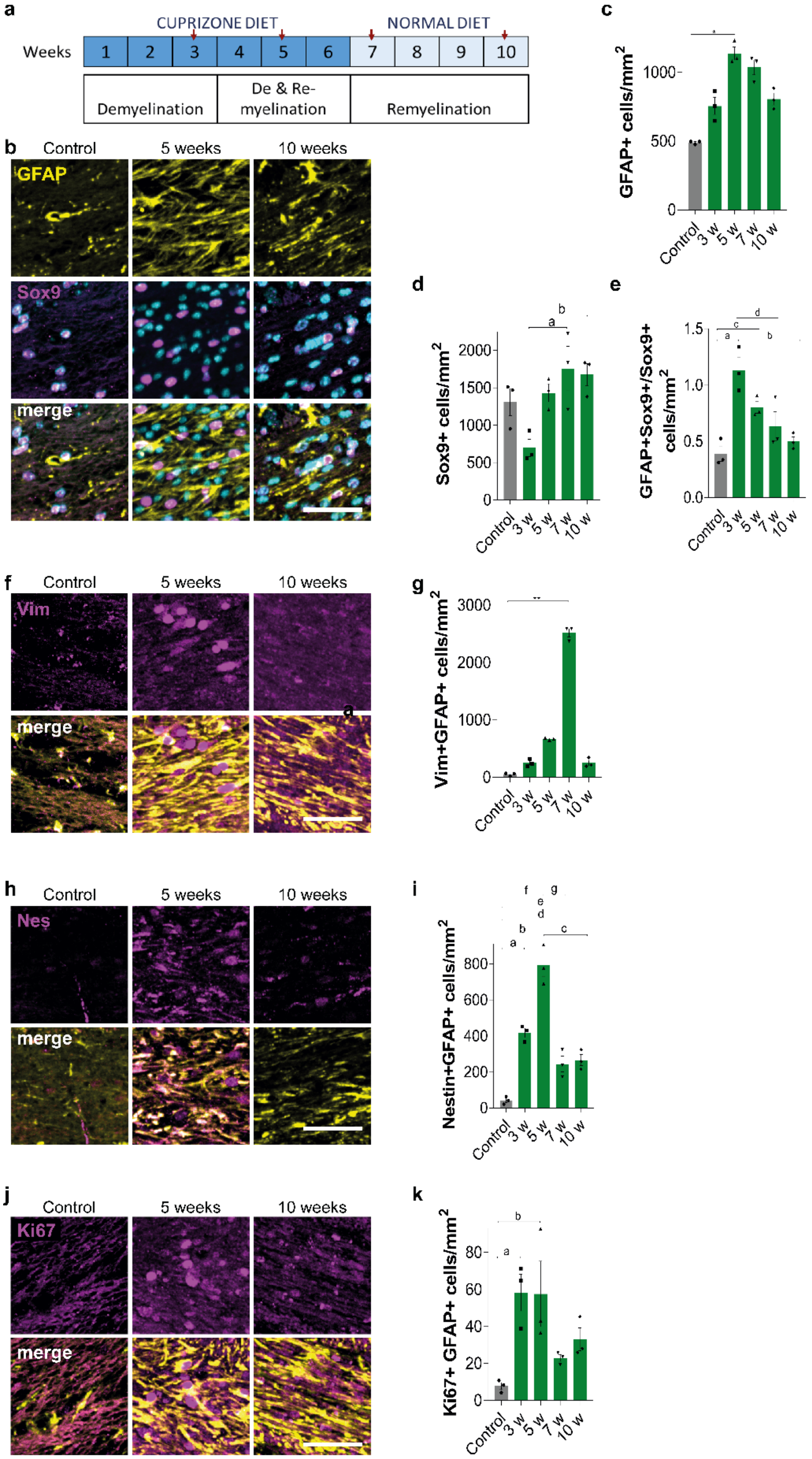

**a.** Schematic representation of the cuprizone-induced model of demyelination, where red arrowheads indicate key time-points analysed: 3 weeks (demyelination), 5 weeks (concomitant demyelination and early remyelination), 7-10 weeks (late remyelination).

**b.** Astrocytes stained with GFAP (yellow) and SOX9 (magenta) in no-lesion control and at 5 and 10 weeks. Hoechst indicates nuclei in cyan. Scale bar; 50 µm.

**c.** Mean GFAP+ cells/mm$^2$ ± s.e.m. in control and at 3, 5, 7, and 10 weeks (w). Kruskal-Wallis test with Dunn's test correction, **P*= 0.0189. ANOVA summary *P*=0.0001. n=3 mice/group.

**d.** Mean SOX9+ cells/mm$^2$ ± s.e.m. in control and at 3, 5, 7, and 10 w. One-way ANOVA with Tukey's multiple comparison test, $^a$*P*= 0.0152, $^b$*P*= 0.0242. n=3 mice/group.

**e.** Mean GFAP+SOX9+/SOX9+ cell ratio ± s.e.m. in control and at 3, 5, 7, and 10 w. One-way ANOVA with Tukey posthoc test, $^a$*P*=0.0010, $^b$*P*=0.0092, $^c$*P*=0.0033, $^d$*P*=0.0165. ANOVA summary *P*=0.0011 F=10.99. n=3 mice/group.

**f.** Astrocytes stained with GFAP (yellow) and Vim (magenta) in no-lesion control and at 5 and 10 w. Scale bar; 50 µm.

**g.** Mean Vim+GFAP+ cells/mm$^2$ ± s.e.m.. in control and at 3, 5, 7, and 10 w. Kruskal-Wallis test with Dunn's test correction, ***P*=0.0099. ANOVA summary *P*<0.0001. n=3 mice/group.

**h.** Astrocytes stained with GFAP (yellow) and Nestin (Nes; magenta) in no-lesion control and at 5 and 10 w. Scale bar; 50 µm.

**i.** Mean Nes+GFAP+ cells/mm$^2$ ± s.e.m.. in control and at 3, 5, 7, and 10 w. One-way ANOVA and Tukey's multiple comparison test, $^a$*P*=0.0004, $^b$*P*<0.0001, $^c$*P*<0.0001, $^d$*P*=0.0160, $^e$*P*=0.0291, $^f$*P*=0.0004, $^g$*P*<0.0001. ANOVA summary *P*<0.0001 F=50.85. n=3 mice/group.

**j.** Proliferative astrocytes stained with GFAP (yellow) and Ki67 (magenta) in no-lesion control and at 5 and 10 w. Scale bar; 50 µm.

**k.** Mean Ki67+GFAP+ cells/mm$^2$ ± s.e.m.. in control and at 3, 5, 7, and 10 w. One-way ANOVA with Tukey's multiple comparison test, $^a$*P*=0.0252, $^b$*P*=0.0274. ANOVA summary *P*=0.0150 F=5.284. n=3 mice/group.

**Extended Data Figure 4. Significantly upregulated and downregulated genes in astrocytes during remyelination.**

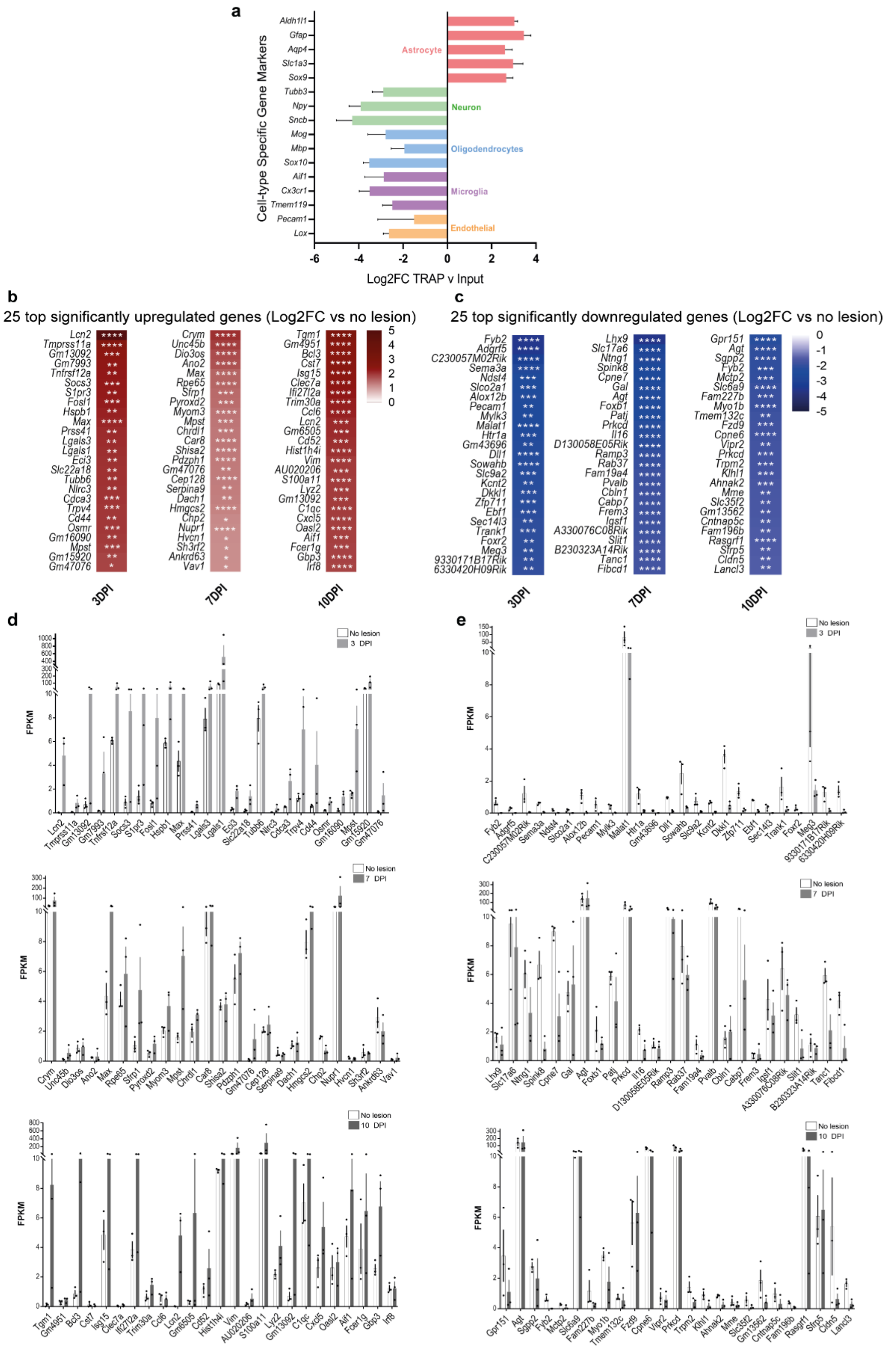

**a.** Log2 Fold-change (FC) in TRAP versus pre-TRAP (input) samples demonstrates enrichment of genes associated with astrocytes (*Aldh1l1, Gfap, Aqp4, Slc1a3, Sox9*) but not those associated with neurons (*Tubb3, Npy, Sncb*), oligodendrocytes (*Mog, Mbp, Sox10*), microglia (*Aif1, Cx3cr1, Tmem119*), or endothelial cells (*Pecam1, Lox*).
**b.** Heat map of 25 top significantly upregulated genes at 3, 7, and 10 DPI relative to no-lesion control represented as Log2 Fold change (FC). DE-Seq2 Benjamini–Hochberg-adjusted *P*-value. *P<0.05, **P<0.01, ***P<0.001, ****P<0.0001.
**c.** Heat map of 25 top significantly downregulated genes at 3, 7, and 10 DPI relative to no-lesion control represented as Log2 Fold change (FC). DE-Seq2 Benjamini–Hochberg-adjusted *P*-value. *P<0.05, **P<0.01, ***P<0.001, ****P<0.0001.
**d.** Mean FPKM values ± s.e.m. of the 25 top significantly upregulated genes at 3, 7, and 10 DPI.
**e.** Mean FPKM values ± s.e.m. of the 25 top significantly downregulated genes at 3, 7, and 10 DPI.

**Extended Data Figure 5. Astrocytes present a mixed inflammatory and neuroprotective signature during remyelination.**

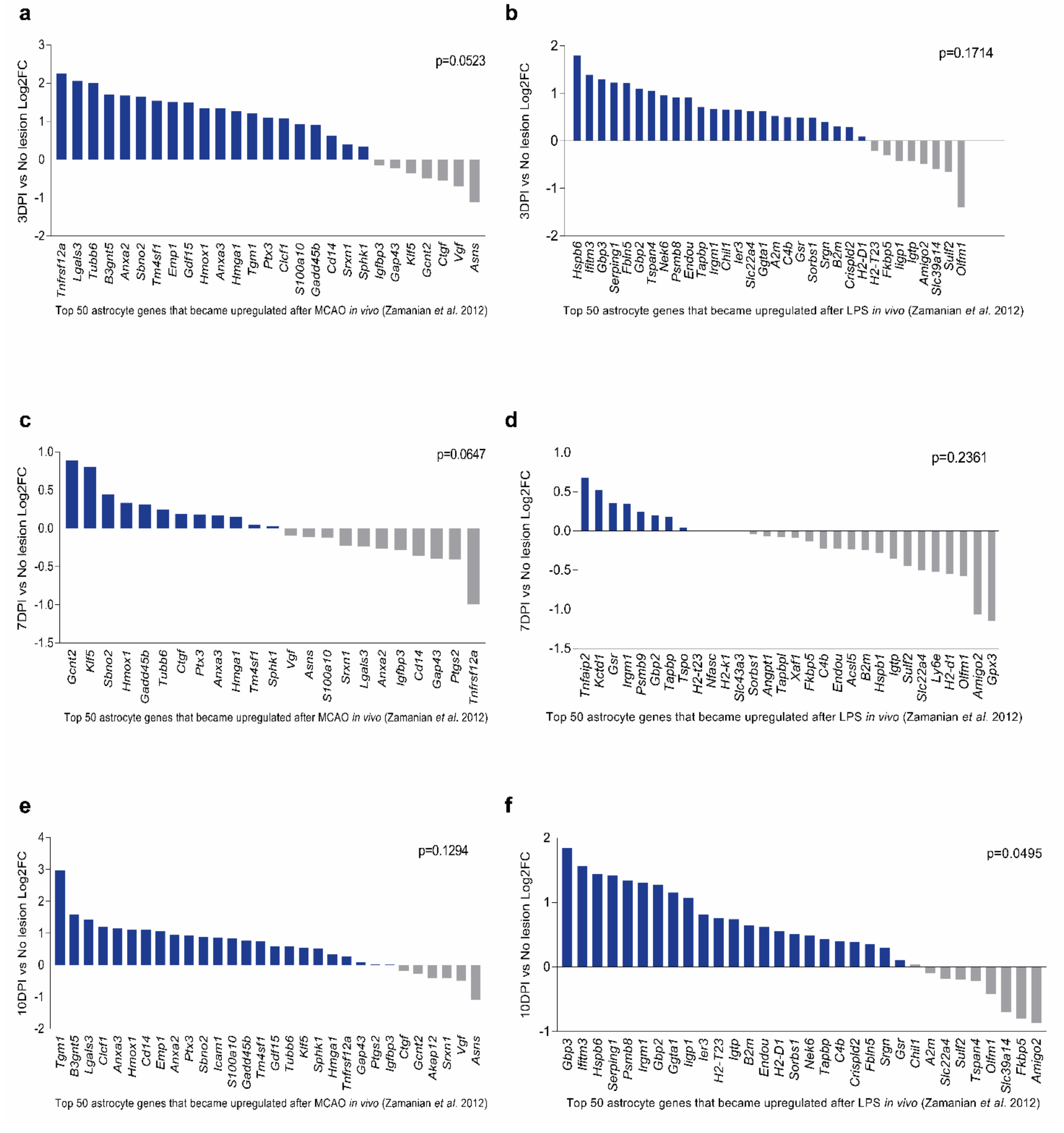


Log2 Fold change (FC) compared to no-lesion control at 3 DPI (a, b), 7 DPI (c, d) and 10 DPI (e, f) for the top 50 genes exclusively upregulated in ‘A2’ astrocytes (after cerebral artery occlusion; a, c, e) and exclusively upregulated in ‘A1’ astrocytes (after LPS injection; b, d, f). Genes expressed >0.5 FPKM were included in the analysis. Paired *t*-tests are between the average FPKM compared to no-lesion control.

**Extended Data Figure 6. Gene Ontology pathway analysis of astrocytes during remyelination.**

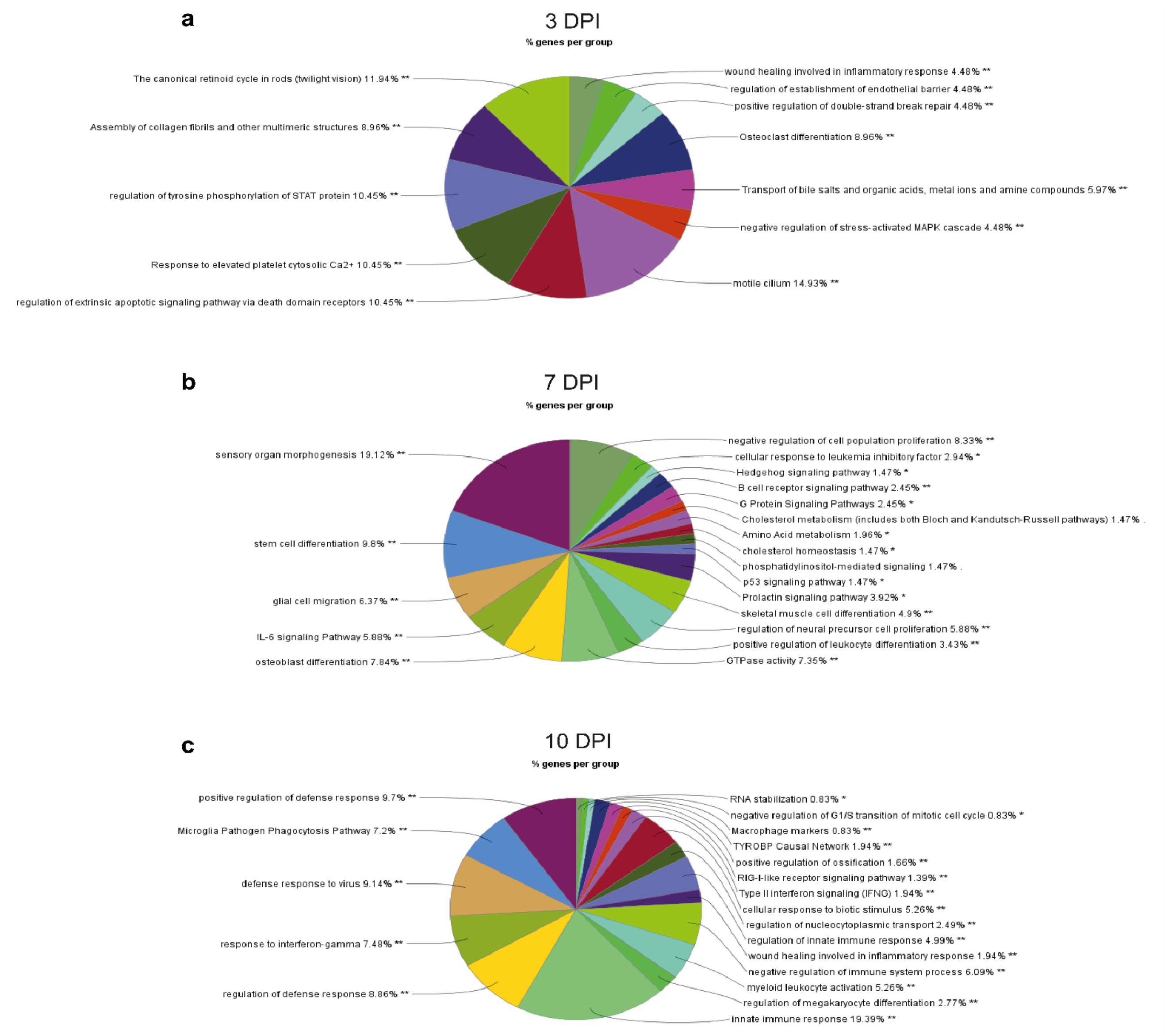


Pie charts showing enriched pathways of the most significantly upregulated Gene Ontology (GO) Terms with a 2-fold change and adjusted p-value of <0.05 at 3 DPI (**a**), 7 DPI (**b**), and 10 DPI (**c**). GO terms are grouped using functional grouping (percentage of genes in each GO group) and based on highest significance. Each term was defined with a minimum of 3 genes.

**Extended Data Figure 7. Nrf2 and cholesterol pathway activation in astrocytes in the cuprizone remyelination model.**

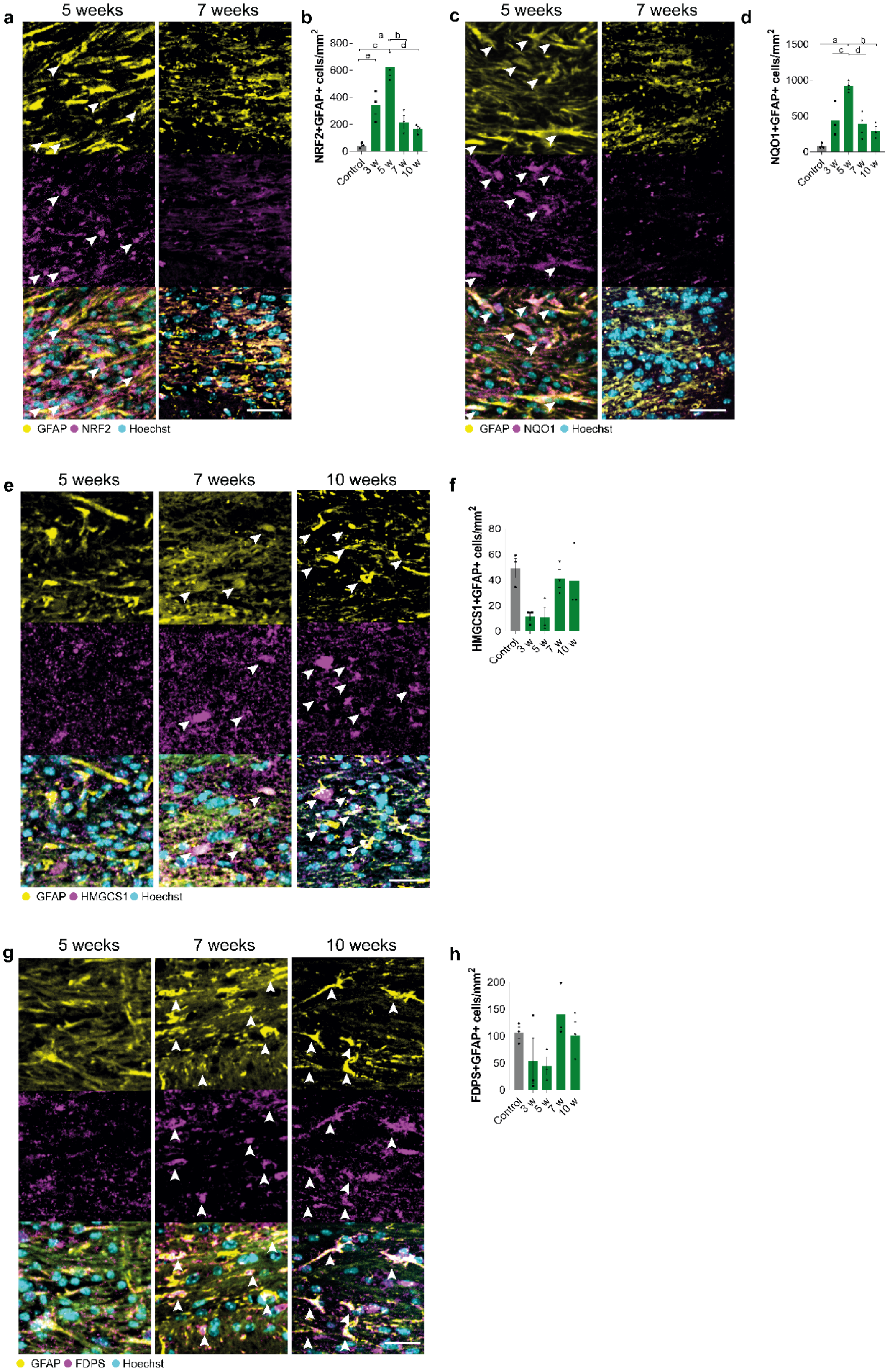

- **a.** Representative images of astrocytes (GFAP+; yellow) expressing NRF2 (magenta) counterstained with Hoechst (cyan), in the corpus callosum of mice during cuprizone-induced demyelination and early remyelination (5 weeks) or during late remyelination on normal diet (7 weeks). Arrows indicate double positive cells. Scale bar, 25µm.
- **b.** Mean densities of NRF2+ GFAP+ astrocytes ± s.e.m. in control mice, during cuprizone-induced demyelination (3, 5 weeks) and during remyelination on normal diet (7, 10 weeks). One-way ANOVA; a: $P$=0.0121, b: $P$ =0.0008, c: $P$ <0.0001, d: $P$ =0.0003, e $P$=0.0081. ANOVA summary $P$<0.0001 F=21.94 n=3 mice/group.
- **c.** Representative images of astrocytes (GFAP+; yellow) expressing Nrf2 target NQO1 (magenta), counterstained with Hoechst (cyan), in the corpus callosum at 5 and 7 weeks of cuprizone paradigm. Arrows indicate double positive cells. Scale bar, 25µm.
- **d.** Mean densities of NQO1+ GFAP+ astrocytes ± s.e.m. in control mice and from 3-10 weeks. One way ANOVA; a: $P$=0.0004, b: $P$ =0.0035, c: $P$ =0.0219, d: $P$ =0.0109. ANOVA summary $P$=0.0007 F=12.25. n=3 mice/group.
- **e.** Representative images of astrocytes (GFAP+; yellow) expressing HMGCS1 (magenta), counterstained with Hoechst (cyan), in the corpus callosum at 5 and 7 weeks of cuprizone paradigm. Arrows indicate double positive cells. Scale bar, 25µm.
- **f.** Mean densities of HMGCS1+ GFAP+ astrocytes ± s.e.m. in control mice and from 3-10 weeks. One-way ANOVA summary $P$=0.0333 F=4.045. n=3 mice/group.
- **g.** Representative images of astrocytes (GFAP+; yellow) expressing FDPS (magenta), counterstained with Hoechst (cyan), in the corpus callosum at 5 and 7 weeks of cuprizone paradigm. Arrows indicate double positive cells. Scale bar, 25µm.
- **h.** Mean densities of FDPS+ GFAP+ astrocytes ± s.e.m. in control mice and from 3-10 weeks. One-way ANOVA summary $P$=0.1494 F=2.145. n=3 mice/group.

**Extended Data Figure 8. Oligodendrocyte lineage cell activation of Nrf2 and cholesterol biosynthesis pathways during remyelination.**

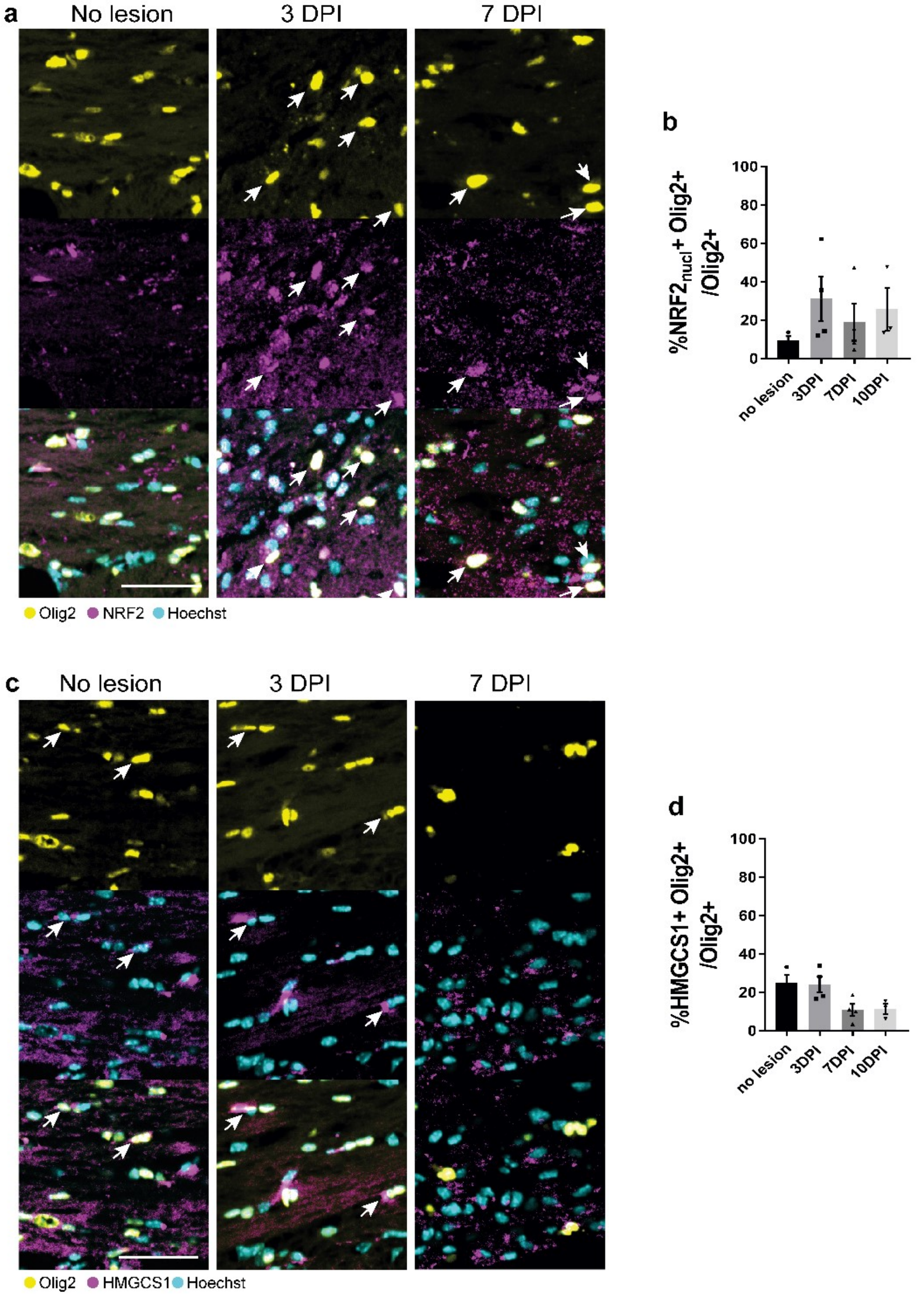


- **a.** Representative images of oligodendrocyte lineage cells (Olig2+; yellow) expressing NRF2 (magenta), counterstained with Hoechst (cyan), in the corpus callosum in non-lesioned control and at 3 and 7 DPI. Arrows indicate double positive cells. Scale bar, 50 µm.
- **b.** Mean percentage of Olig2+ cells expressing activated (nuclear) NRF2 ± s.e.m. in non-lesioned corpus callosum and at 3, 7, and 10 DPI. One-way ANOVA summary $P$=0.8304 F=5.0470. n=4 mice/group.
- **c.** Representative images of oligodendrocyte lineage cells (Olig2+; yellow) expressing HMGCS1 (magenta), counterstained with Hoechst (cyan), in the corpus callosum in non-lesioned control and at 3 and 7 DPI. Arrows indicate double positive cells. Scale bar, 50 µm.

**d.** Mean percentage of Olig2+ cells expressing HMGCS1 ± s.e.m. in non-lesioned corpus callosum and at 3, 7, and 10 DPI. One-way ANOVA summary $P$=0.0317 F=4.427. n=4 mice/group.

**Extended Data Figure 9. Astrocyte responses in GFAP-Nrf2 demyelinated lesions.**

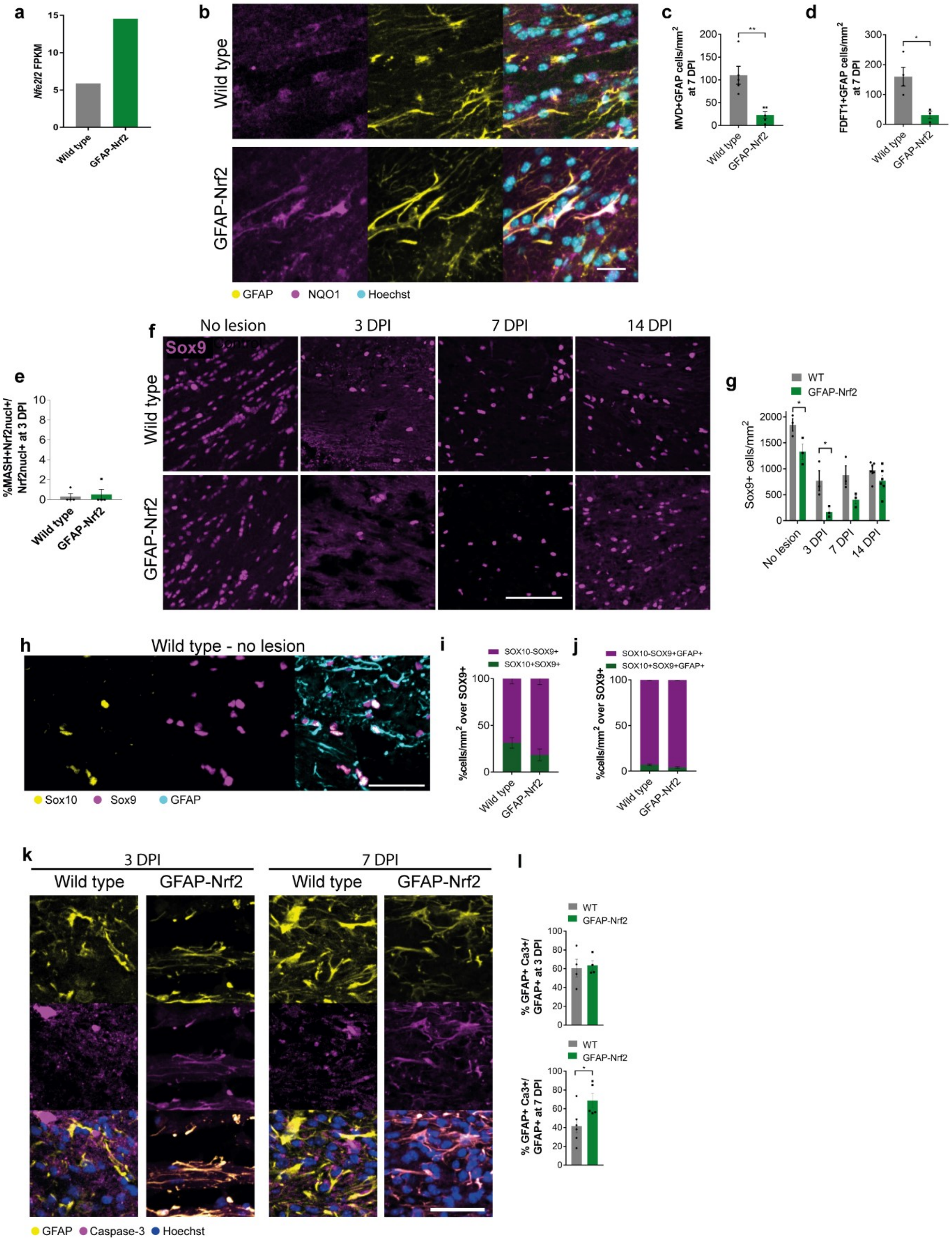


**a.** Expression of the Nrf2 gene *Nfe2l2* in wildtype and GFAP-Nrf2 astrocytes as represented by Fragments Per Kilobase per Million mapped reads (FPKM).

**b.** Astrocytes (GFAP+; yellow) positive for the Nrf2 target NQO1 (magenta) in non-lesioned corpus callosum of wild type control and GFAP-Nrf2 mice. Hoechst indicates nuclei in cyan. Scale bar, 25 µm.

**c.** Mean MVD+GFAP+ cells/$mm^2$ ± s.e.m. in WT and GFAP-Nrf2 mice at 7 and 14 DPI. 2-tailed unpaired Student's *t*-test with Welch's correction wild type vs GFAP-Nrf2, 7 DPI $P$=0.0087 t=4.137. n= 4-5 mice/condition.

**d.** Mean FDFT1+GFAP+ cells/$mm^2$ ± s.e.m. in WT and GFAP-Nrf2 mice at 7 and 14 DPI. 2-tailed unpaired Student's *t*-test with Welch's correction wild type vs GFAP-Nrf2, 7 DPI $P$=0.0196 t=3.815. n= 3-4 mice/condition.

**e.** Percentage of nuclear Nrf2+ cells expressing the neural stem cell marker MASH1 at 3 DPI in wild type and GFAP-Nrf2 corpus callosum. Kolmogorov-Smirnov test $P$>0.9999. n=4 mice/group.

**f.** SOX9+ astrocytes in wild type and GFAP-Nrf2 corpus callosum (outlined) in no-lesion control and at 3, 7 and 14 DPI. Scale bar, 100 µm.

**g.** Mean SOX9+ cells/$mm^2$ ± s.e.m in wild-type (WT) and GFAP-Nrf2 mice in no-lesion control and at 3, 7, and 14 DPI. Two-way ANOVA with Bonferroni correction wild type vs GFAP-Nrf2, a: $P$=0.0427, b: $P$=0.0151. Two-way ANOVA summary (Interaction F(3,22)= 1.321, $P$-value=0.2927; Row (time-point) Factor F(3,22)=28.03, $P$-value<0.0001; Column (mouse genotype) factor F(1,22) = 26.65, $P$-value<0.0001). n=3-6 mice/ group.

**h.** Rare examples of GFAP+ (cyan) SOX9+ (magenta) cells expressing SOX10 (yellow). Scale bar, 50 µm.

**i.** Percentage of SOX9+ cells ± s.e.m. which are SOX10+ (green) or SOX10- (purple) in wild type and GFAP-Nrf2 mice. Two-way ANOVA with Bonferroni correction wild type vs GFAP-Nrf2, no significant. Two-way ANOVA summary (Interaction F(1,6)= 3.916, $P$-value=0.0952; Row (mouse genotype) Factor F(1,6)=0, $P$-value>0.9999; Column (type of cell) factor F(1,6) = 59.38, $P$-value=0.0003). n=3 mice/group.

**j.** Percentage of SOX9+ cells ± s.e.m. which are GFAP+SOX10+ (green) or GFAP+SOX10- (purple) in wildtype and GFAP-Nrf2 mice. Two-way ANOVA with Bonferroni correction wild type vs GFAP-Nrf2, no significant. Two-way ANOVA summary (Interaction F(1,6)= 10.38, $P$-value=0.0181; Row (mouse genotype) Factor F(1,6)=0.002923, $P$-value=0.9586; Column (type of cell) factor F(1,6) = 9153, $P$-value<0.0001). n=3 mice/group.

**k.** GFAP+ astrocytes (yellow) expressing active-Caspase-3+ (magenta) in wild type and GFAP-Nrf2 mouse corpus callosum (outlined) in no-lesion control and at 3, and 7 DPI. Hoechst indicates nuclei in blue. Scale bar, 25 µm.

**l.** Mean percentage of active-Caspase-3 (Ca3)+ GFAP+ cells over total GFAP+ cells ± s.e.m. in WT and GFAP-Nrf2 mice at 3 and 7 DPI. 2-tailed unpaired Student's *t*-test with Welch's correction wild type vs GFAP-Nrf2 at 3 DPI $P$=0.7964 t=0.2736. Kolmogorov-Smirnov at 7 DPI $P$=0.0260. n=4-6 mice/group.

**Extended Data Figure 10. Oligodendrocyte and microglial responses in GFAP-Nrf2 demyelinated lesions.**

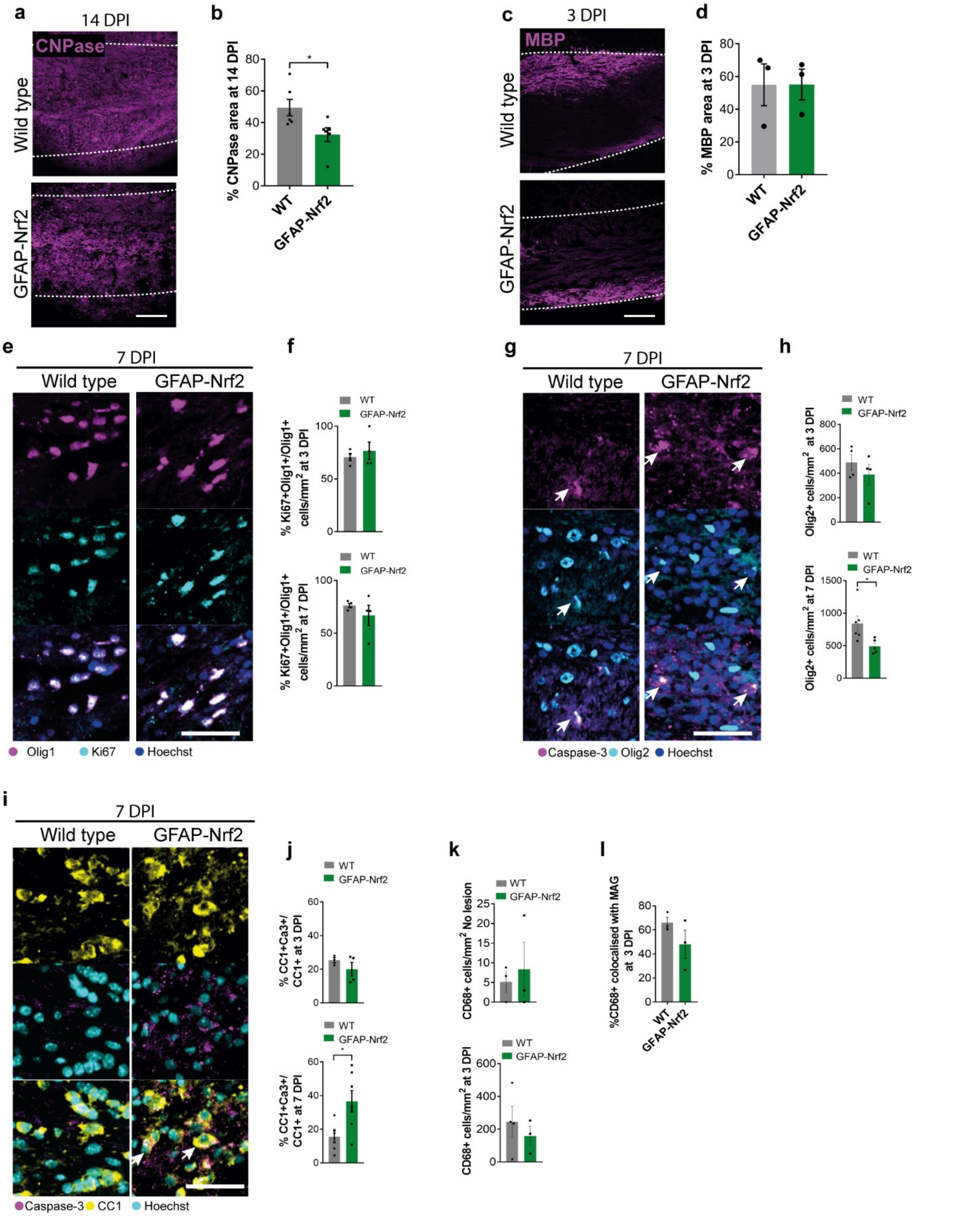

**a.** CNPase staining (magenta) in the corpus callosum (outlined) in wild type and GFAP-Nrf2 mice at 14 DPI. Scale bar, 100 µm.

**b.** Percentage of area of corpus callosum with CNPase staining ± s.e.m. in wild type (WT) and GFAP-Nrf2 mice at 14 DPI. 2-tailed unpaired Student's *t*-test with Welch's correction, **P*=0.0299 t=2.53. n=6 mice/group.

**c.** MBP staining (magenta) in the corpus callosum (outlined) in wild type and GFAP-Nrf2 mice at 3 DPI. Scale bar, 100 µm.

**d.** Percentage of area of corpus callosum with MBP staining ± s.e.m. in WT and GFAP-Nrf2 mice at 3 DPI. 2-tailed unpaired Student's *t*-test with Welch's correction, *P*=0.9933 t=0.009007, n=3 mice/group.

**e.** Oligodendrocyte precursors (Olig1+, magenta) which are proliferating (Ki67; cyan) in WT and GFAP-Nrf2 mice at 7 DPI. Hoechst indicates nuclei in blue. Scale bar, 25 µm.

**f.** Mean percentage of Olig1+ cells which are Ki67+ ± s.e.m. in WT and GFAP-Nrf2 mice at 3 and 7 DPI. 2-tailed unpaired Student's *t*-test with Welch's correction, 3 DPI *P*=0.5547 t=0.6443, 7 DPI *P*=0.4051 t=0.9545. n=3-4 mice/group.

**g.** Oligodendrocyte lineage cells (Olig2+; cyan) which are apoptotic (active Caspase-3+; magenta) in WT and GFAP-Nrf2 mouse corpus callosum at 7 DPI. Hoechst indicates nuclei in blue. Scale bar, 25 µm.

**h.** Total number of oligodendrocyte lineage cells (Olig2+) cells/$mm^2$ ± s.e.m. in WT and GFAP-Nrf2 mice at 3 and 7 DPI. 2-tailed unpaired Student's *t*-test with Welch's correction, 3 DPI *P*=0.3854 t=0.9403, 7 DPI **P*=0.0222 t=3.054. n=4-6 mice/group.

**i.** Oligodendrocytes (CC1+; yellow) which are apoptotic (active Caspase-3+; magenta) in WT and GFAP-Nrf2 mouse corpus callosum at 7 DPI. Hoechst indicates nuclei in blue. Scale bar, 25 µm.

**j.** Mean percentage of oligodendrocytes (CC1+) which are active Caspase-3+ ± s.e.m. in WT and GFAP-Nrf2 mice at 3 and 7 DPI. 2-tailed unpaired Student's *t*-test with Welch's correction, 3 DPI *P*=0.2910 t=1.22, 7 DPI **P*=0.0176 t=2.897. n=34-7 mice/group.

**k**. Mean number of CD68+ microglia/macrophages ± s.e.m. per $mm^2$ in non-lesioned mice and 3 DPI lesions in WT and GFAP-Nrf2 mice. 2-tailed unpaired Student's *t*-test with Welch's correction, no lesion *P*=0.7004 t=0.4303, 3 DPI *P*=0.4742 t=0.7773. n=3 mice/group.

**l**. Mean percentage of CD68 area co-localized with myelin associated glycoprotein (MAG) ± s.e.m. at 3 DPI in WT and GFAP-Nrf2 lesions. 2-tailed unpaired Student's *t*-test with Welch's correction, *P*=0.2632 t=1.43. n=3 mice/group.

**Extended Data Figure 11. Astrocyte depletion is associated with oligodendrocyte death during remyelination.**

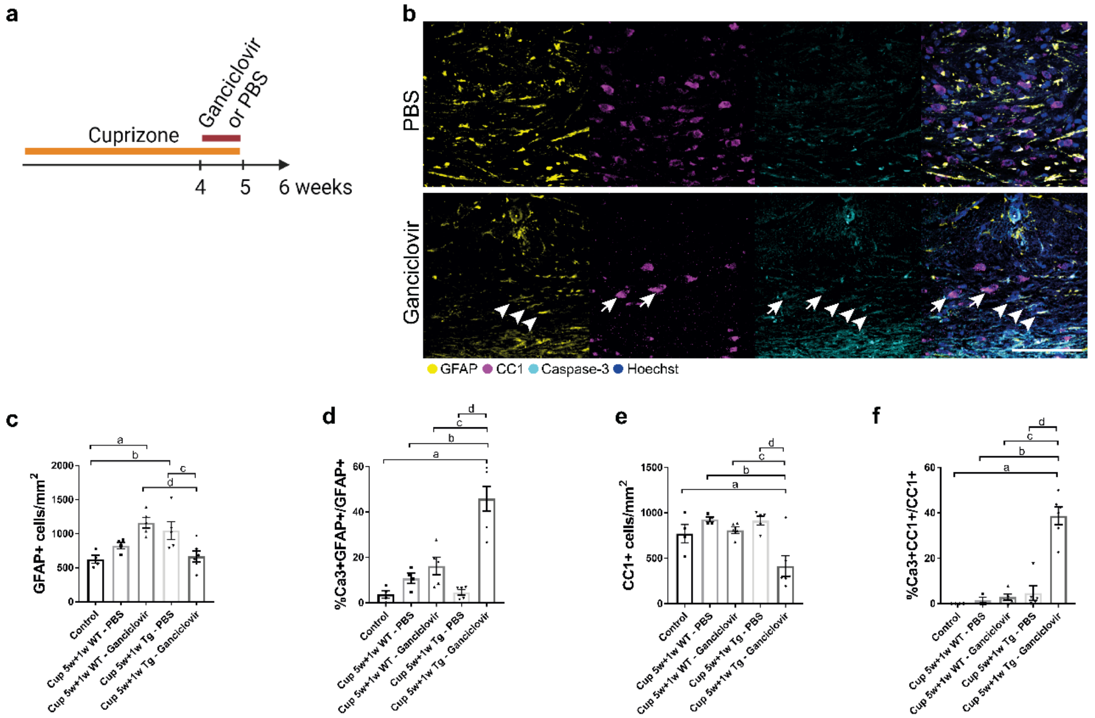


**a.** Cuprizone diet was provided to wildtype mice or GFAP-thymidine kinase mice for 5 weeks to induce demyelination. Ganciclovir was administered from weeks 4-5 at the onset of remyelination to induce astrocyte death. PBS vehicle administration served as a control. Mice were sacrificed at 6 weeks after 1 week on normal diet.

**b.** Astrocyte (GFAP+; yellow) and oligodendrocyte (CC1+; magenta) expressing the apoptotic marker active caspase-3 (cyan), and counterstained with Hoechst (blue). Apoptotic astrocytes are indicated with arrowheads, whilst apoptotic oligodendrocytes are indicated with arrows. Scale bar, 50µm.

**c.** Mean number of GFAP+ astrocytes ± s.e.m. per mm$^2$ in control diet mice, wildtype (WT) mice fed with cuprizone and treated with PBS or ganciclovir, and transgenic mice (Tg) fed with cuprizone treated with PBS or ganciclovir. One way ANOVA with Tukey's multiple comparison test, a: $P$=0.0054, b: $P$ =0.034, c: $P$ =0.0362, d: $P$ =0.0047. ANOVA summary (F=6.887 and $P$-value=0.0013). n=4 mice/group.

**d.** Mean percentage of GFAP+ astrocytes positive for active caspase-3 ± s.e.m. in the above conditions. One way ANOVA with Tukey's multiple comparison test; a-d, $P$<0.0001. ANOVA summary (F=7.699 and $P$-value=0.0007). n=4 mice/group.

**e.** Mean number of CC1+ oligodendrocytes ± s.e.m. in the above conditions. One way ANOVA with Tukey's multiple comparison test; a: $P$=0.0383, b: $P$ =0.002, c: $P$ =0.0117, d: $P$ =0.0013. ANOVA summary (F=36.73 and $P$-value<0.0001). n=4 mice/group.

**f.** Mean percentage of CC1+ oligodendrocytes positive for active caspase-3 ± s.e.m. in the above conditions. One way ANOVA with Tukey's multiple comparison test; a-d $P$<0.0001. ANOVA summary (F=24.1 and $P$-value<0.0001). n=4 mice/group.

**Extended Data Figure 12. Oligodendrocyte and microglial responses to CS-6253 treatment in GFAP-Nrf2 mice.**

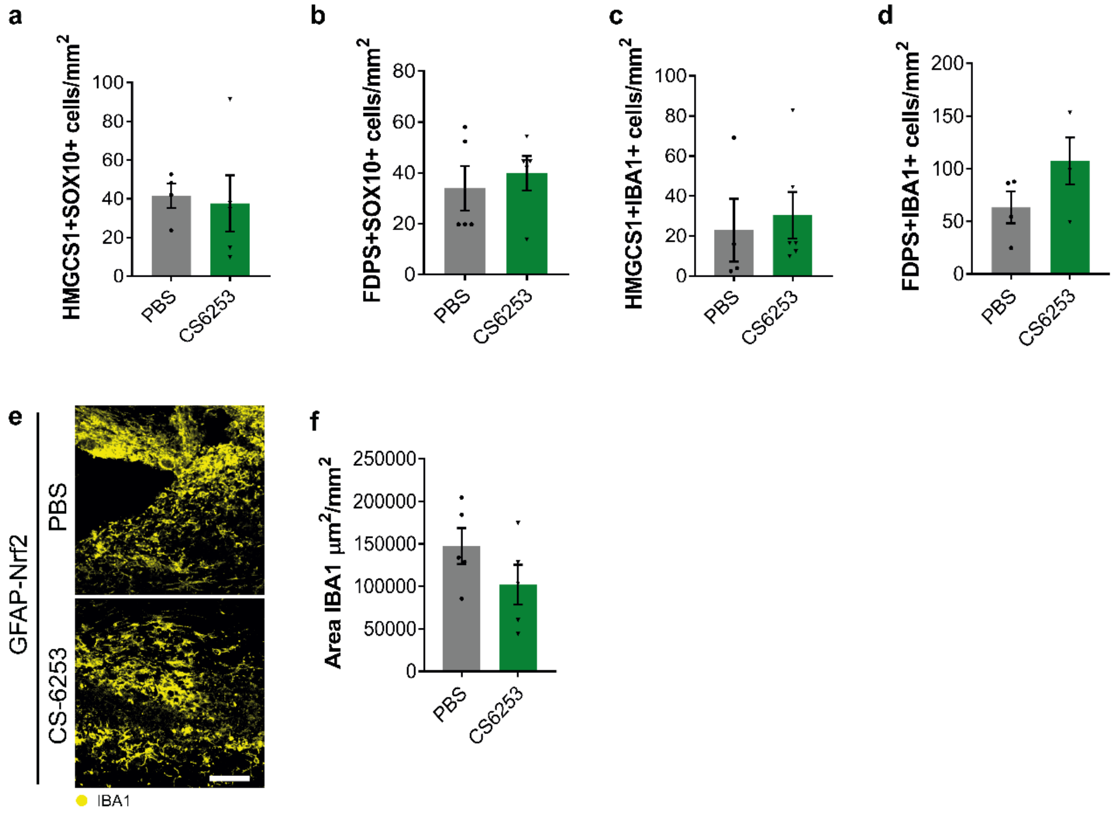


**a.** Mean number of oligodendrocyte lineage cells (SOX10+) expressing HMGCS1 per mm$^2$ ± s.e.m. in GFAP-Nrf2 mice treated with PBS or CS-6253. 2-tailed unpaired Student's *t*-test with Welch's correction, *P*=0.8128 t=0.2486. n=3 mice/group.

**b.** Mean number of FDPS+SOX10+ cells per mm$^2$ ± s.e.m. in GFAP-Nrf2 mice treated with PBS or CS-6253. 2-tailed unpaired Student's *t*-test with Welch's correction, *P*=0.6108 t=0.5308. n=3 mice/group.

**c.** Mean number of microglia/macrophages (IBA1+) expressing HMGCS1 per mm$^2$ ± s.e.m. in GFAP-Nrf2 mice treated with PBS or CS-6253. 2-tailed unpaired Student's *t*-test with Welch's correction, *P*=0.7150 t=0.3826. n=3 mice/group.

**d.** Mean number of FDPS+IBA1+ cells ± s.e.m. per mm$^2$ in GFAP-Nrf2 mice treated with PBS or CS-6253. 2-tailed unpaired Student's *t*-test with Welch's correction, *P*=0.1587 t=1.642. n=3 mice/group.

**e.** IBA1 staining in PBS or CS-6253 treated GFAP-Nrf2 mice. Scale bar, 50µm.

**f.** Mean area of IBA1 staining (µm$^2$) per mm$^2$ ± s.e.m. in PBS or CS-6253 treated GFAP-Nrf2 mice. 2-tailed unpaired Student's *t*-test with Welch's correction, *P*=0.1886 t=1.438. n=5 mice/group.

**Extended Data Figure 13. Conditional knockout of Nrf2 in astrocytes**

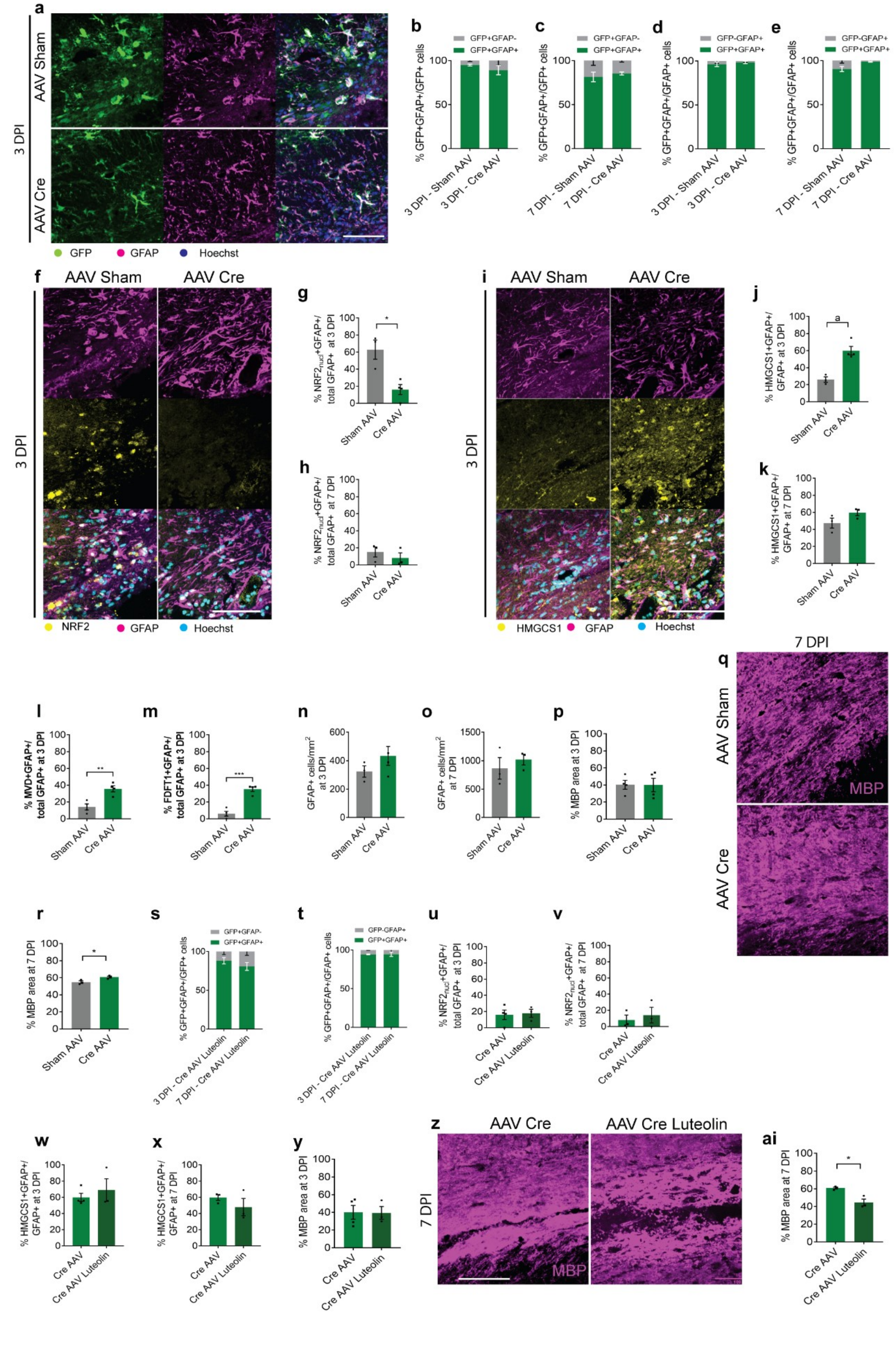

**a.** GFP expression (green) colocalized with GFAP (magenta) and counterstained with Hoechst (blue), following injection of *Nfe2l2* floxed mice with AAV5-GFAP(0.7)-eGFP-T2A-iCre ('AAV-Cre') or AAV5-GFAP(0.7)-eGFP ('AAV Sham') into the corpus callosum. Scale bar, 100 μm.
**b**. Specificity of viral transduction indicated as mean percentage of GFP+ signal which was colocalized with GFAP ± s.e.m. at 3 DPI following injection with Sham AAV or Cre AAV. N=4 mice per condition.
**c**. Specificity of viral transduction indicated as mean percentage of GFP+ signal which was colocalized with GFAP ± s.e.m. at 7 DPI following injection with Sham AAV or Cre AAV. N=3 mice per condition.
**d**. Efficiency of viral transduction indicated as mean percentage of GFAP+ cells which were GFP+ ± s.e.m. at 3 DPI following injection with Sham AAV or Cre AAV. N=4 mice per condition.
**e**. Efficiency of viral transduction indicated as mean percentage of GFAP+ cells which were GFP+ ± s.e.m. at 7 DPI following injection with Sham AAV or Cre AAV. N=3 mice per condition.
**f**. Astrocytes (GFAP+; magenta) expressing nuclear Nrf2 (yellow) and counterstained with Hoechst (blue) in AAV Sham and AAV Cre mice at 3 DPI. Scale bar, 100 μm.
**g.** Mean percentage of GFAP+ cells with nuclear Nrf2 at 3 DPI in Sham AAV and Cre AAV mice ± s.e.m. 2-tailed unpaired Student's t-test, $^{a}P$ =0.0104 t=5.186. N=4 mice per condition.
**h.** Mean percentage of GFAP+ cells with nuclear Nrf2 at 7 DPI in Sham AAV and Cre AAV mice ± s.e.m.. 2-tailed unpaired Student's test, not significant *P*=0.6235 t=0.531. N=3 mice per condition.
**i.** Astrocytes (GFAP+; magenta) expressing HMGCS1 (yellow) and counterstained with Hoechst (blue) in AAV Sham and AAV Cre mice at 3 DPI. Scale bar, 100 μm.
**j.** Mean percentage of GFAP+ cells expressing HMGCS1 at 3 DPI in Sham AAV and Cre AAV mice ± s.e.m.. Kolmogorov-Smirnov test, $^{a}P$ =0.0571. N=3-4 mice per condition.
**k.** Mean percentage of GFAP+ cells expressing HMGCS1 at 7 DPI in Sham AAV and Cre AAV mice ± s.e.m.. 2-tailed unpaired Student's test, not significant *P*=0.3226 t=1.125. N=3 mice per condition.
**l.** Mean percentage of GFAP+ cells expressing MVD at 7 DPI in Sham AAV and Cre AAV mice ± s.e.m.. 2-tailed unpaired Student's *t*-test, *P*=0.0052 t=4.274. N=4 mice per condition.
**m.** Mean percentage of GFAP+ cells expressing FDFT1 at 7 DPI in Sham AAV and Cre AAV mice ± s.e.m.. 2-tailed unpaired Student's *t*-test, *P*=0.0003 t=7.337. N=4 mice per condition.
**n.** Mean astrocyte numbers (GFAP+) in AAV Sham and AAV Cre mice at 3 DPI ± s.e.m.. 2-tailed unpaired Student's test, not significant *P*=0.2192 t=1.42. N=3 mice per condition.
**o.** Mean astrocyte numbers (GFAP+) in AAV Sham and AAV Cre mice at 7 DPI ± s.e.m..2-tailed unpaired Student's test, not significant *P*=0.5200 t=0.7291. N=3 mice per condition.
**p.** Mean percentage of MBP area at 3 DPI (indicating demyelination) in Sham AAV and AAV Cre mice ± s.e.m.. 2-tailed unpaired Student's test, not significant *P*=0.9759 t=0.03147. N=4 mice per condition.
**q.** Mean percentage of MBP area (indicating remyelination) in AAV Sham and AAV Cre mice at 7 DPI Scale bar, 100 μm.
**r.** Mean percentage of MBP area in AAV Sham and AAV Cre mice at 7 DPI± s.e.m.. 2-tailed unpaired Student's t-test, $^{a}P$ =0.0366 t=3.09. N=3 mice per condition.
**s.** Mean percentage of GFP+ signal colocalized with GFAP in AAV-Cre + Luteolin treated mice at 3 and 7 DPI ± s.e.m.. N=3 mice per condition.
**t.** Mean percentage of GFAP+ cells expressing GFP in AAV-Cre + Luteolin treated mice at 3 and 7 DPI ± s.e.m.. N=3 mice per condition.
**u.** Mean percentage of GFAP+ cells with nuclear Nrf2 expression at 3 DPI in AAV-Cre ± Luteolin treated mice ± s.e.m.. 2-tailed unpaired Student's test, not significant *P*=0.9752 t=0.03237. N=3 mice per condition.
**v.** Mean percentage of GFAP+ cells with nuclear Nrf2 expression at 7 DPI in AAV-Cre ± Luteolin treated mice ± s.e.m.. 2-tailed Student's test, not significant. *P*=0.6235 t=0.531**.** N=3 mice per condition.
**w.** Mean percentage of GFAP+ cells expressing HMGCS1 at 3 DPI in AAV-Cre ± Luteolin treated mice ± s.e.m.. Kolmogorov-Smirnov test, not significant *P*=0.8857. N=3-4 mice per condition.
**x.** Mean percentage of GFAP+ cells expressing HMGCS1 at 7 DPI in AAV-Cre ± Luteolin treated mice ± s.e.m.. 2-tailed unpaired Student's test, not significant *P*=0.3254 t=1.12. N=3 mice per condition.

**y.** Mean percentage area of MBP at 3 DPI (indicating demyelination) in AAV-Cre ± Luteolin treated mice ± s.e.m.. 2-tailed Student's test, not significant. $P$=0.9545 t=0.05997. N=3-4 mice per condition.

**z.** MBP immunostaining at 7 DPI in AAV-Cre or AAV-Cre+Luteolin-treated mice. Scale bar, 100 µm.

**ai.** Mean percentage area of MBP at 7 DPI (indicating remyelination) in AAV-Cre ± Luteolin treated mice ± s.e.m.. 2-tailed unpaired Student's t-test, [a]$P$ =0.0146 t=4.118. N=3 mice per condition.

**Extended Data Figure 14. Primary astrocyte uptake of fluorescent cholesterol analogue.**

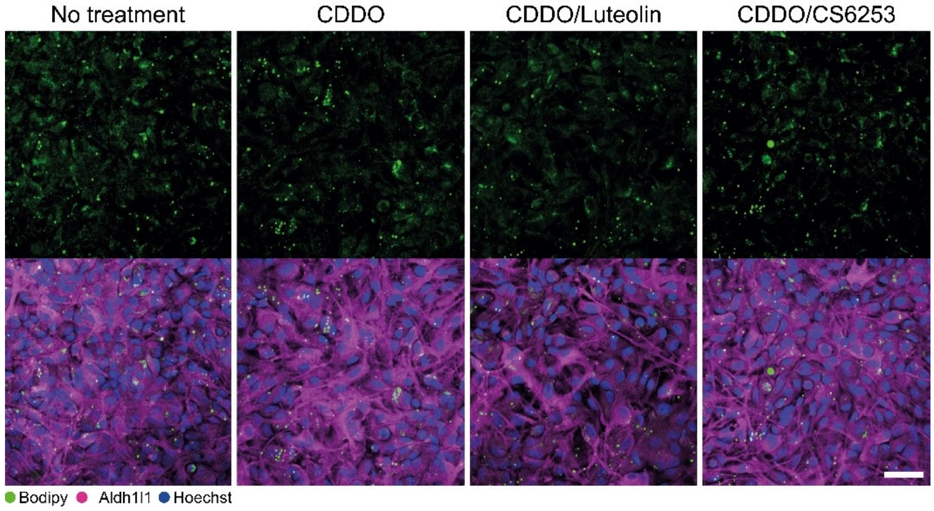


Primary astrocytes (Aldh1l1+; magenta) having taken up Bodipy-FL-C12 fluorescent cholesterol analogue (green), counterstained with Hoechst (blue). Astrocytes were either untreated, or treated with $CDDO^{TFEA}$ to induce Nrf2 activation, then either untreated or treated with the Nrf2 inhibitor Luteolin or the cholesterol biosynthesis pathway inducer/efflux stimulator CS-6253. Scale bar, 50μm.

**Extended Data Figure 15. Astrocyte responses in multiple sclerosis lesions.**

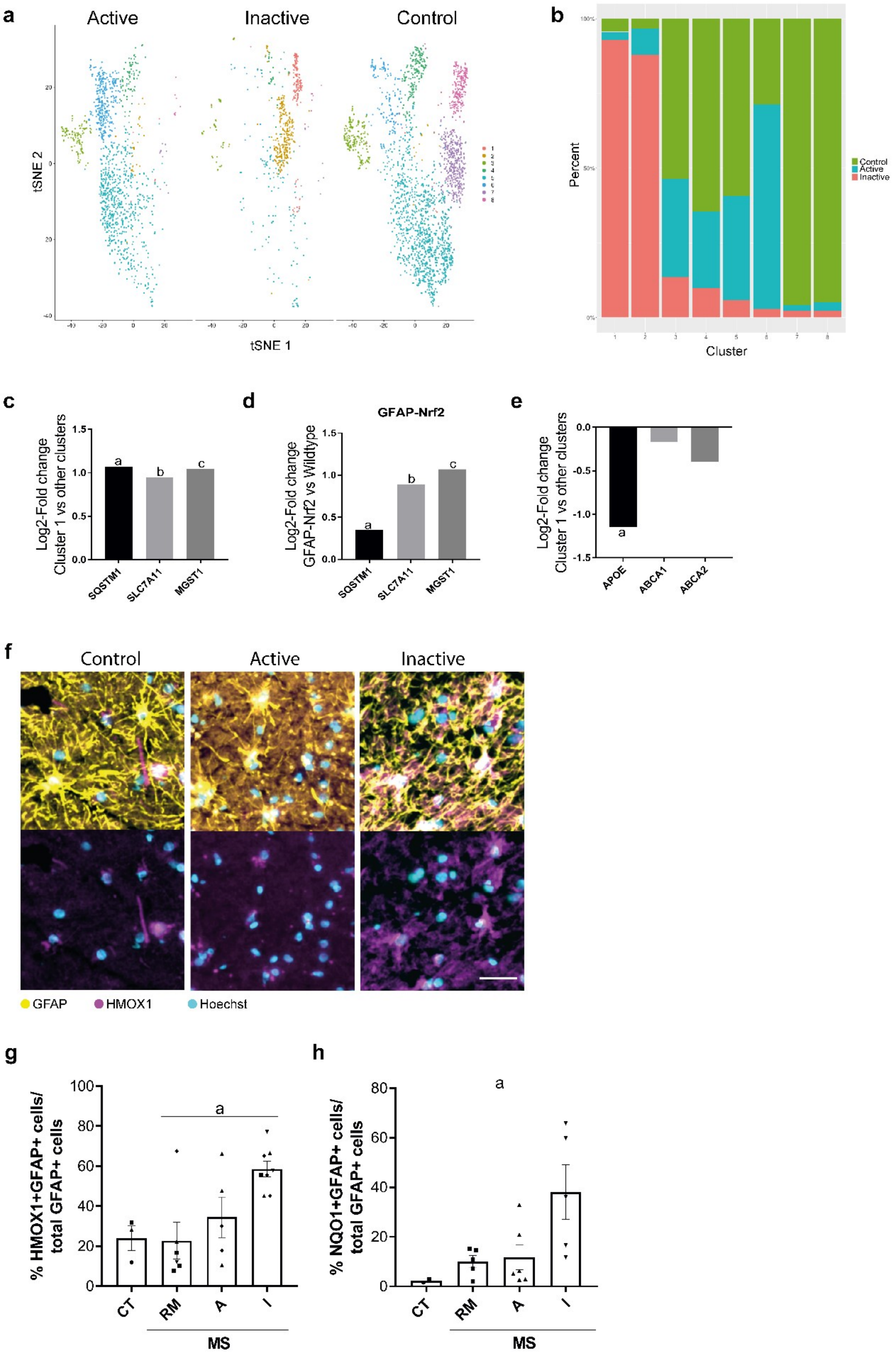

**a.** tSNE plot of single nuclei sequencing of astrocytes from control white matter, or active and inactive multiple sclerosis (MS) lesions. Legend indicates cluster colour assignments.

**b.** Percent frequency distribution of astrocyte clusters among control, and active and inactive MS lesions.

**c.** Log2 Fold Change of expression of Nrf2-target genes in Cluster 1 over other clusters. $^{a}P$=7.83x10$^{-8}$, $^{b}P$=1.91x10$^{-9}$, $^{c}P$=1.26x10$^{-11}$.

**d.** GFAP-Nrf2 mouse astrocyte expression of Nrf2 target genes upregulated in human Cluster 1 astrocytes. Represented as Log2-fold change vs wild type control astrocytes. DESeq2, $^{a}P$=0.0137, $^{b}P$=5.88x10$^{-37}$, $^{c}P$=1.84x10$^{-16}$.

**e.** Log2 Fold Change of expression of cholesterol export genes in Cluster 1 over other clusters. $^{a}P$=1.97 x 10$^{-3}$.

**f.** Astrocytes (GFAP+; yellow) expressing HMOX1 (magenta) counterstained with Hoechst (cyan) in control human white matter, and active and inactive multiple sclerosis lesions. Scale bar, 25µm.

**g.** Mean percentage of GFAP+ cells expressing HMOX1 ± s.e.m. in control (CT), fully remyelinated (RM), active (A) and inactive (I) multiple sclerosis (MS) lesions. n=3 controls and 5-8 MS lesions per type. Kruskal-Wallis and Dunn's multiple comparisons test, $^{a}P$=0.0419.

**h.** Mean percentage of GFAP+ cells expressing NQO1 ± s.e.m. in CT, RM, A, and I MS lesions. n=3 controls and 5-8 MS lesions per type. Kruskal-Wallis and Dunn's multiple comparisons test, $^{a}P$=0.0353.

**Extended Data Figure 16. Responses to Luteolin treatment.**

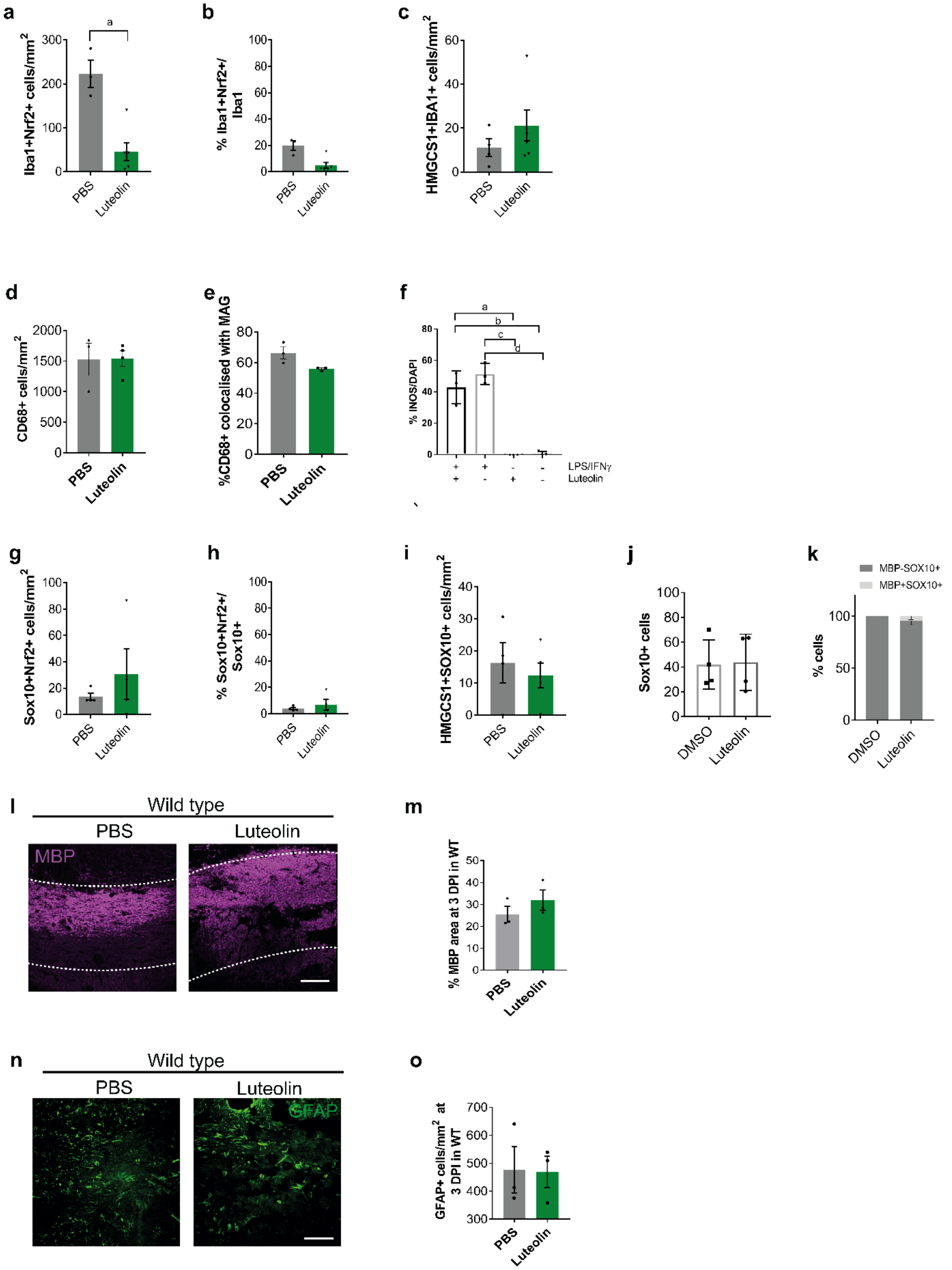

**a.** Mean number of microglia/macrophages (IBA1+) expressing Nrf2 per $mm^2$ ± s.e.m. in PBS or Luteolin treated GFAP-Nrf2 mice. Kolmogorov-Smirnov test, *P*=0.0238. n=3 mice/group.

**b.** Percentage of IBA1+ cells expressing Nrf2 per $mm^2$ ± s.e.m. in PBS or Luteolin treated GFAP-Nrf2 mice. Kolmogorov-Smirnov test, not significant *P*=0.0952. n=3 mice/group.

**c.** Mean number of IBA1+ HMGCS1+ cells per $mm^2$ ± s.e.m. in PBS or Luteolin treated GFAP-Nrf2 mice. 2-tailed unpaired Student's t-test, not significant *P*=0.0952 t=1.248. n=3 mice/group.

**d.** Mean number of microglia/macrophages (CD68+) per $mm^2$ ± s.e.m. in PBS or Luteolin treated GFAP-Nrf2 mice. 2-tailed unpaired Student's t-test, not significant *P*=0.9544 t=0.06005. n=3 mice/group.

**e.** Percentage of CD68 area co-localized with myelin associated glycoprotein (MAG) per $mm^2$ ± s.e.m. in PBS or Luteolin treated GFAP-Nrf2 mice. 2-tailed unpaired Student's t-test, not significant *P*=0.0573 t=2.645. n=3 mice/group.

**f.** Primary microglia cultures were treated with LPS and IFNγ and assessed for expression of iNOS. A subset of cells were treated with Luteolin. One way ANOVA with Tukey's multiple comparison test, a: *P*=0.0002, b: *P*=0.0002, c: *P*<0.0001, d: *P*<0.0001. ANOVA summary (*P*<0.0001 F=55.75). n=3 independent litters.

**g.** Mean number of oligodendrocyte lineage cells (SOX10+) per $mm^2$ ± s.e.m. in PBS or Luteolin treated GFAP-Nrf2 mice. 2-tailed unpaired Student's t-test, not significant. *P*=0.4857. n=3 mice/group.

**h.** Percentage of SOX10+ cells expressing Nrf2 per $mm^2$ ± s.e.m. in PBS or Luteolin treated GFAP-Nrf2 mice. Kolmogorov-Smirnov test, not significant *P*=0.5193 t=0.7207. n=3 mice/group.

**i.** Mean number of SOX10+ HMGCS1+ cells per $mm^2$ ± s.e.m. in PBS or Luteolin treated GFAP-Nrf2 mice. 2-tailed unpaired Student's t-test, not significant *P*=0.6182 t=0.5303. n=3 mice/group.

**j.** Numbers of oligodendrocyte lineage cells (SOX10+) ± s.e.m. in primary cultures treated with Luteolin or DMSO vehicle control. 2-tailed unpaired Student's t-test, not significant *P*=0.9049 t=0.1247. n=3 independent litters.

**k.** Percentage of cultured SOX10+ cells ± s.e.m. which were MBP+ (light grey) or MBP- (dark grey). Two-way ANOVA with Bonferroni correction wild type vs GFAP-Nrf2, no significant. Two-way ANOVA summary (Interaction $F(1,24)$= 1.173, *P*-value=0.2895; Row (condition) Factor $F(1,24)$=0, *P*-value>0.9999; Column (type of cell) factor $F(1,24)$ = 498.9, *P*-value<0.0001). n=3 litters.

**l.** Wildtype lesioned mice at 3 DPI following treatment with Luteolin or PBS from 0-3 DPI, immunostained for myelin debris using myelin basic protein (MBP). Scale bar, 100μm.

**m.** Percentage of lesion area covered by MBP myelin debris ± s.e.m. at 3 DPI in wildtype (WT) mice following treatment with Luteolin or PBS. 2-tailed unpaired Student's t-test, not significant *P*=0.3304 t=1.115. n=3 mice/group.

**n.** Wildtype lesioned mice at 3 DPI following treatment with Luteolin or PBS from 0-3 DPI, immunostained for GFAP. Scale bar, 100μm.

**o.** Mean number of GFAP+ cells ± s.e.m. at 3 DPI in WT mice following treatment with Luteolin or PBS. 2-tailed unpaired Student's t-test, not significant *P*=0.9447 t=0.07441. n=3 mice/group.

**Extended Data Figure 17. Summary Diagram.**

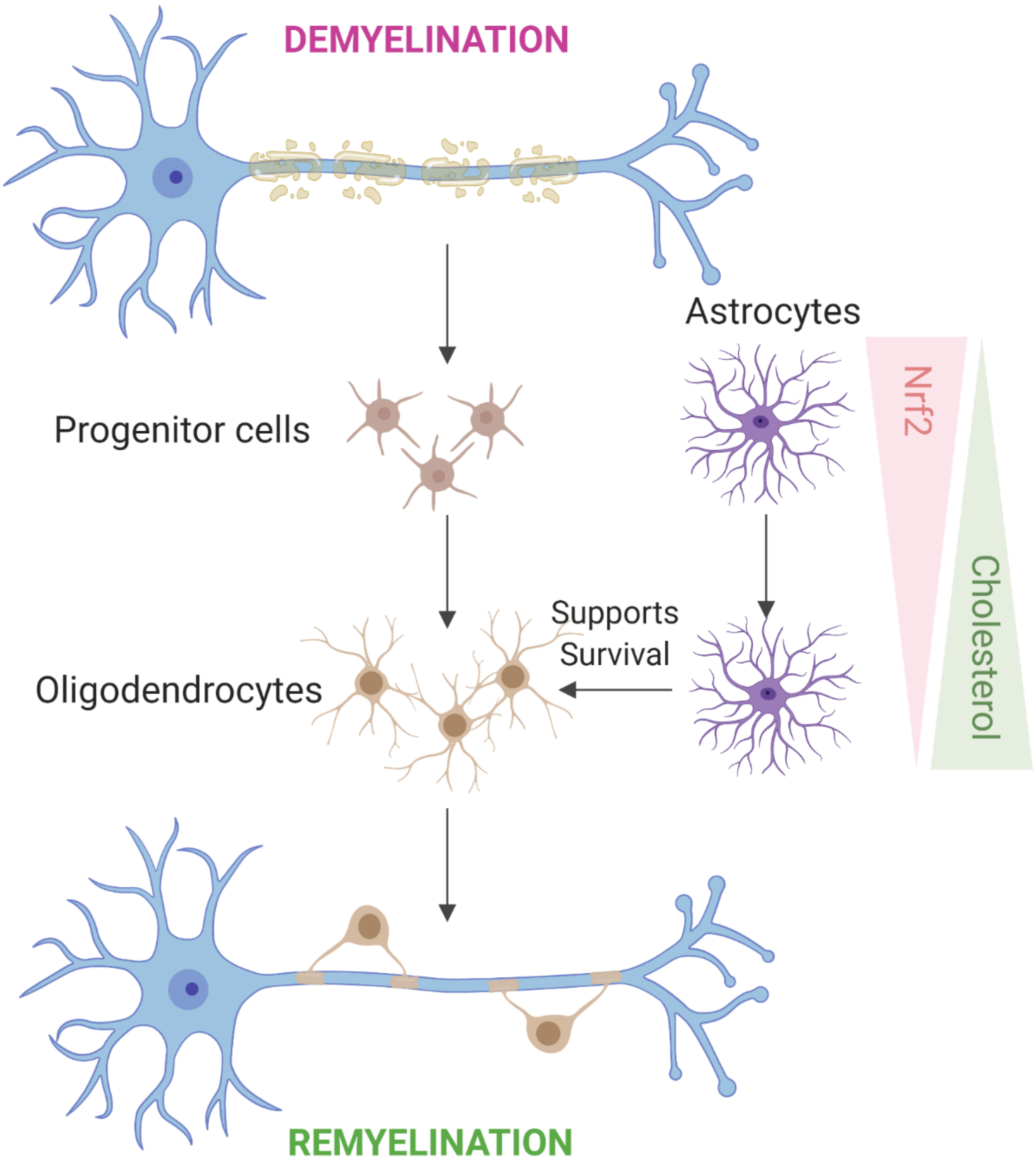

### Extended Data Table 1. Clinical information on human samples.

Clinical information on human brain tissue samples.MS=multiple sclerosis; SP= secondary progressive; PP= primary progressive; F=female; M=male; NA=not available; A=Active; I=Inactive; R=Remyelinated.

| | | | | | | | Lesions | | |
|---|---|---|---|---|---|---|---|---|---|
| | **ID** | **MS type** | **Sex** | **Age at death** | **Cause of death** | **Disease duration (years)** | **A** | **I** | **R** |
| **MS** | **MS121** | SP | F | 49 | MS | 14 | 2 | - | - |
| | **MS122** | SP | M | 44 | Bronchopneumonia | NA | 1 | 2 | - |
| | **MS100** | SP | M | 46 | NA | 8 | 1 | 3 | 1 |
| | **MS207** | SP | F | 46 | NA | 25 | 1 | - | 1 |
| | **MS176** | PP | M | 37 | Intestinal obstruction | 27 | - | 1 | 3 |
| | **MS136** | SP | M | 40 | NA | 9 | 2 | - | 2 |
| | **MS242** | SP | M | 57 | Sepsis | 19 | - | 3 | 2 |
| | **MS230** | SP | F | 42 | NA | 19 | 3 | - | - |
| **Control** | **CO14** | - | M | 64 | Cardiac failure | - | - | - | - |
| | **CO25** | - | M | 35 | Carcinoma of the tongue | - | - | - | - |
| | **CO28** | - | F | 60 | Ovarian cancer | - | - | - | - |

**Supplemental Sheet 1 Legend. TRAP sequencing of astrocytes during remyelination.**
Fragments per Kilobase Million (FPKM) and Log2 fold change of genes identified from TRAPseq of Aldh1l1-Rpl10-eGFP LPC-demyelinated lesions at 3, 7, and 10 days post injection (DPI) and no lesion controls.
